# The discriminatory power of the T cell receptor

**DOI:** 10.1101/2020.11.16.384495

**Authors:** Johannes Pettmann, Anna Huhn, Enas Abu-Shah, Mikhail Kutuzov, Daniel B. Wilson, Michael L. Dustin, Simon J. Davis, P. Anton van der Merwe, Omer Dushek

**Author notes:** Corresponding author: Omer Dushek. These authors contributed equally (Johannes Pettmann, Anna Huhn, Enas Abu-Shah).

## Abstract

T cells use their T-cell receptors (TCRs) to discriminate between lower-affinity self and higher affinity non-self pMHC antigens. Although the discriminatory power of the TCR is widely believed to be near-perfect, technical difficulties have hampered efforts to precisely quantify it. Here, we describe a method for measuring very low TCR/pMHC affinities, and use it to measure the discriminatory power of the TCR, and the factors affecting it. We find that TCR discrimination, although enhanced compared with conventional cell-surface receptors, is imperfect: primary human T cells can respond to pMHC with affinities as low as K_D_ ~1 mM. The kinetic proofreading mechanism fit our data, providing the first estimates of both the time delay (2.8 s) and number of biochemical steps (2.67) that are consistent with the extraordinary sensitivity of antigen recognition. Our findings explain why self pMHC frequently induce autoimmune diseases and anti-tumour responses, and suggest ways to modify TCR discrimination.

## Introduction

T cells use their T-cell receptors (TCRs) to discriminate between lower-affinity self and higher-affinity non-self peptides presented on Major Histocompatibility Complexes (pMHCs). This ability is the cornerstone of adaptive immunity and defects in this process can lead to autoimmunity. Although the strength of discrimination is widely believed to be near-perfect for the TCR (1–8), systematic measurements to quantify it have not been performed.

Early influential studies using three murine TCRs suggested a sharp affinity threshold for T cell activation (9–14). Using T cells from the OT-I, 3L.2, and 2B4 transgenic TCR mice, it was shown that subtle changes to their cognate peptides, which apparently produced modest 3-5-fold decreases in affinity, abolished T cell responses even when increasing the peptide concentration by as much as 100,000-fold (9–15). Although this near-perfect discrimination based on affinity could be explained by a kinetic proof-reading (KP) mechanism (16), it could not also account for the ability of T cells to respond to few pMHC ligands (high sensitivity) (17, 18). Consequently, there has been a focus on identifying mechanisms that can simultaneously explain near-perfect discrimination and high sensitivity (1–8, 15). However, near-perfect discrimination is inconsistent with evidence that T cells can respond to lower affinity self-antigens (19, 20) and moreover, that T cell-mediated autoimmunity is associated with increased expression of self-antigens (21, 22). There is thus a discrepancy between the current notion of near-perfect TCR discrimination and data on the role of T cell recognition of self-pMHC in human disease.

A key challenge in assessing discrimination is the accurate measurements of very weak TCR/pMHC affinities, with K_D_ ranging from 1 to >100 *µ*M (23). A highly sensitive method for analysing molecular interactions is surface plasmon resonance (SPR) but even with this method, accurate measurements are difficult to make, especially at 37*^◦^*C. In the case of OT-I for example, measurements were performed at 37*^◦^*C but high affinity bi-phasic binding was observed (11), which has not been observed for other TCRs and may represent protein aggregates that often form at the high concentrations necessary for making these measurements. It follows that the reported small 3-fold change in affinity between the activating OVA and non-activating E1 ligands (11) may be a consequence of multivalent interactions. Indeed, more recent studies found the expected low-affinity mono-phasic binding for OT-I/OVA (24, 25) and no detectable binding for OT-I/E1 (24). This raises the possibility that E1 does not activate T cells not because of near-perfect discrimination but simply because it does not bind the TCR. These studies highlight the challenges of accurately measuring TCR/pMHC affinities and underline their importance in our understanding of antigen discrimination.

Here, we introduce a new SPR protocol that can accurately determine ultra-low TCR/pMHC affinities at 37*^◦^*C into the K_D_ ~ 1 mM regime. We found that T cell responses were gradually lost as the affinity was decreased without a sharp affinity threshold and remarkably, responses were detected to ultra-low affinity pMHCs. By introducing a quantitative measure of discrimination, we are able to not only analyse our data but also analyse the published literature finding that the discriminatory power of the T cell receptor is imperfect yet remains above the baseline produced by other conventional surface receptors.

## Results

### Measurements of ultra-low TCR/pMHC affinities at 37***^◦^***C

To assess discrimination, we first generated ligands to the anti-tumour 1G4 (26) and anti-viral A6 (27) TCRs recognising peptides on HLA-A*02:01. The standard SPR protocol is based on injecting the TCR at increasing concentrations over a pMHC-coated surface (Fig. 1A,B) with the resulting steady-state binding response plotted over the TCR concentration (Fig. 1C). This curve is fitted by a 2-parameter Hill function to determine B_max_ (the maximum response when all pMHC are bound by TCR) and the K_D_, which is the TCR concentration where binding is half the B_max_. Therefore, an accurate determination of K_D_ requires an accurate determination of B_max_.

**Figure 1:**
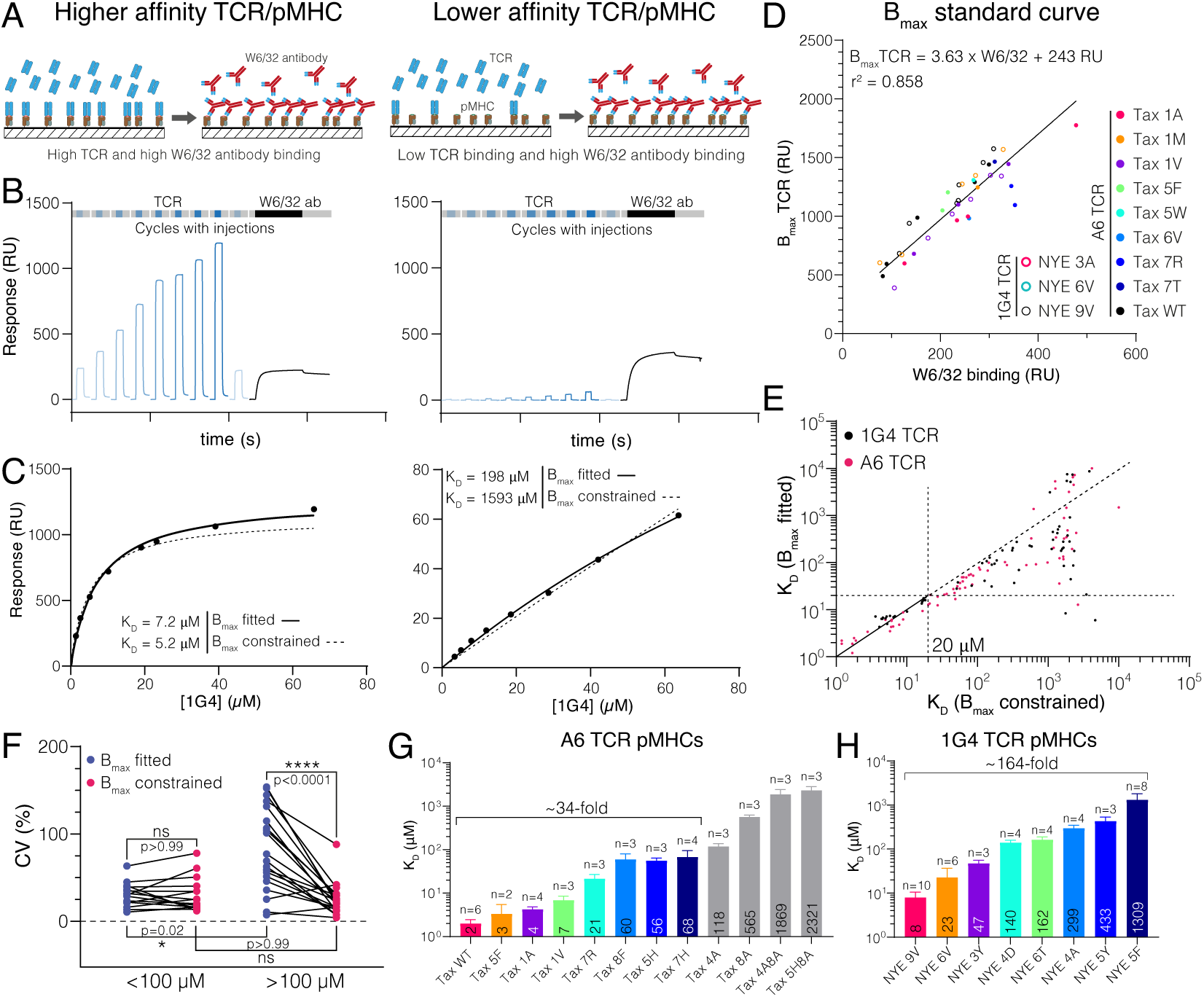
Measuring ultra-low TCR/pMHC affinities using SPR at 37*^◦^*C using a constrained B_max_ method. **(A-C)** Comparison of 1G4 TCR binding to a higher (left panels, 9V) and lower (right panels, 5F) affinity pMHC. **(A)** Schematic comparing TCR and W6/32 binding. **(B)** Example SPR sensograms showing injections of different TCR concentrations followed by the W6/32 antibody. **(C)** Steady-state binding response from **(B)** over the TCR concentration (filled circles) fitted to determine K_D_ when B_max_ is either fitted (standard method) or constrained (new method). **(D)** Empirical standard curve relating W6/32 binding to fitted B_max_ obtained using higher affinity interactions. **(E)** Correlation of K_D_ obtained using the fitted and constrained methods. Each dot represents an individual measurement (n=136; 63 for 1G4 TCR, 73 for A6 TCR). **(F)** Coefficient of variation for higher (<100 *µ*M) or lower affinity (>100 *µ*M) interactions. **(G)** Selected pMHC panel for A6 TCR. **(H)** Selected pMHC panel for 1G4 TCR. Mean values of K_D_ are indicated in bars and ligands used for functional experiments in the main text are coloured.

In the case of the 1G4 TCR binding to its cognate NY-ESO-1 peptide, this protocol produces K_D_ = 7.2 *µ*M (Fig. 1A-C, left column). However, the binding response curves do not saturate for lower affinity pMHCs (Fig. 1A-C, right column). Because of this the fitted B_max_ and therefore the fitted K_D_ may not be accurate. Saturating the binding curves by increasing the TCR concentration is limited by the tendency of soluble recombinant proteins, including the TCR, to accumulate aggregates at high concentrations, which precludes accurate SPR measurements.

To determine B_max_ for low affinity pMHCs, we generated a standard curve using the conformation-sensitive, pan-HLA-A/B/C antibody (W6/32) that only binds correctly folded pMHC (28). By injecting the W6/32 antibody at the end of each experiment (Fig. 1B, black line) we were able to plot the fitted B_max_ from higher affinity interactions (where binding saturated) over the maximum W6/32 binding (Fig. 1D). We observed a linear relationship even when including different TCRs binding different pMHC across multiple protein preparations immobilised at different levels. This strongly suggested that W6/32 and the TCR recognise the same correctly-folded pMHC population and justified the use of the standard curve to estimate B_max_. We noted that W6/32 antibody binding was generally lower than TCR binding (e.g. Fig 1B and a slope of *>* 1 in Fig. 1D), which is unexpected because the molecular weight of the antibody is larger than the TCR. A likely explanation is that by injecting the antibody at a single concentration, we have not saturated antibody binding. This is mitigated by ensuring that the same W6/32 antibody concentration is used and that B_max_ is only interpolated within the standard curve.

We next fitted K_D_ values for 136 interactions using the standard method where B_max_ is fitted and the new method where B_max_ is constrained to the value obtained using the standard curve (Fig. 1E). In the new method, the only fitted parameter is K_D_. Both methods produced similar K_D_ values for higher affinities, validating the method. In contrast, large (100-fold) discrepancies appeared for lower affinity interactions, with the fitted B_max_ method consistently underestimating the K_D_. These large discrepancies were observed despite both methods providing a similar fit (e.g. Fig. 1C, right). This suggested that for the fitted B_max_ method, different combinations of B_max_ and K_D_ can provide a fit of similar quality so that the fitted K_D_ can exhibit large variations for the same interaction (also known as ‘over-fitting’). We explored this by comparing the precision of both methods using the coefficient of variance (CV) of multiple measurements of the same TCR/pMHC combination. We found a similar CV for higher affinity interactions (<100 *µ*M K_D_) and for lower affinity interactions when B_max_ was constrained, but an increased CV for low affinity interaction when B_max_ was fitted (Fig. 1F). Therefore, the standard method has lower precision for low affinity interactions as a result of over-fitting.

We next used the new SPR method to accurately measure ultra-low affinities in order to identify panels of pMHCs that spanned the full physiological affinity range required to quantitate TCR discrimination (Fig. 1G,H).

### Primary human T cells do not display a sharp affinity threshold and respond to ultra-low affinity antigens

To quantify discrimination, we introduced the 1G4 TCR into quiescent naïve or memory CD8^+^ T cells and then co-cultured them with autologous monocyte-derived dendritic cells (moDCs) pulsed with each peptide (Fig. 2A). Using surface CD69 as a marker for T cell activation, we found that lowering the affinity gradually reduced the response without the sharp affinity threshold suggested by near-perfect discrimination and, remarkably, responses were seen to ultra-low affinity peptides, such as NYE 5F (K_D_ =1309 *µ*M; see Fig. 2B,C). To rule out preferential loading and/or stability of ultra-low affinity peptides, we pulsed the TAP-deficient T2 cell lines with all peptides and found similar HLA upregulation, suggesting comparable loading and stability (Fig. S1). We defined pMHC potency as the concentration of peptide required to reach 15 % activation (P_15_) in order to include lower-affinity pMHCs and found that it produced excellent correlations with K_D_ (Fig. 2D,E).

**Figure 2:**
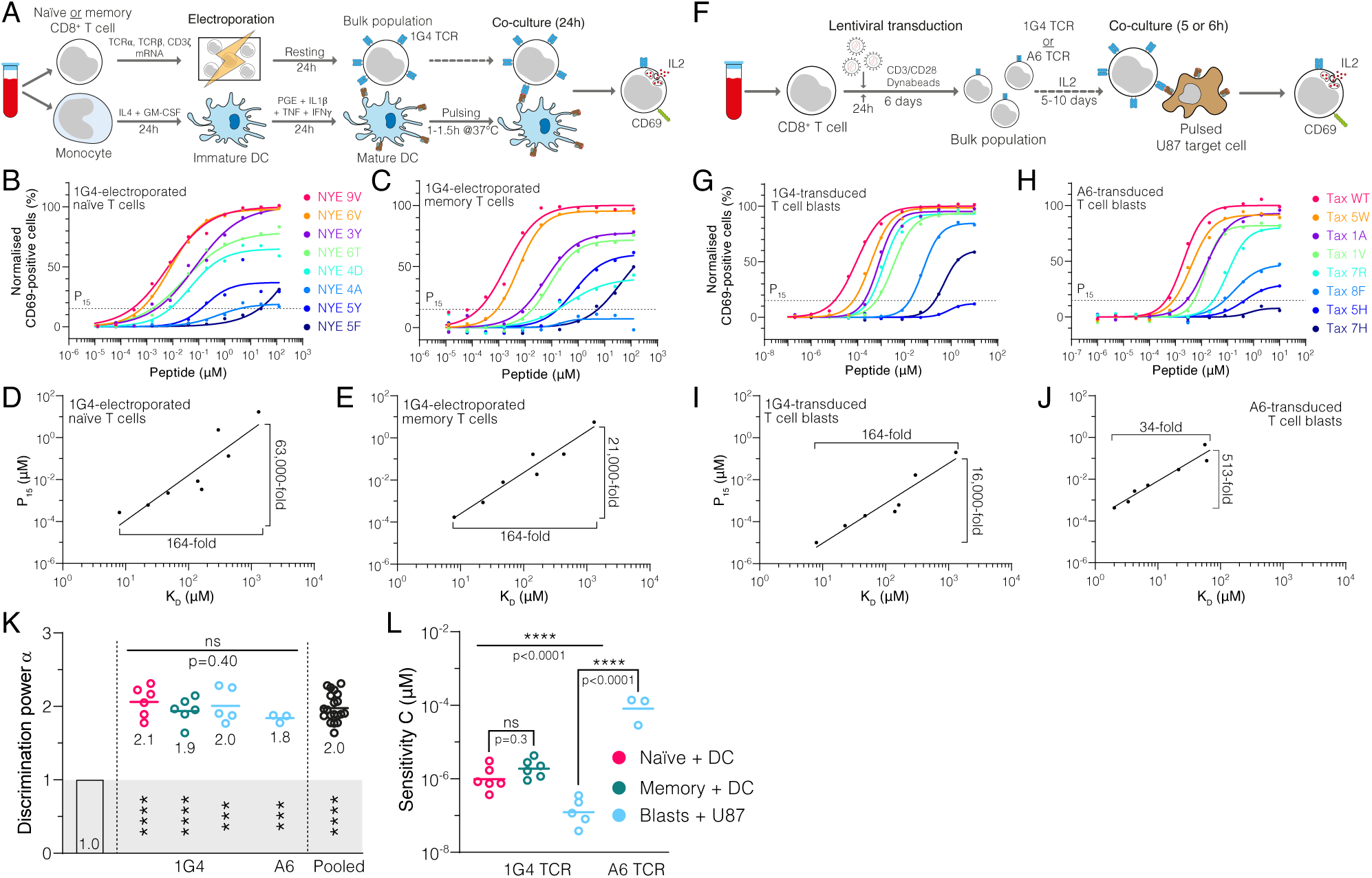
Naïve, memory, and blast human CD8^+^ T cells exhibit enhanced but imperfect discrimination. **(A)** Protocol for producing quiescent primary human naïve and memory CD8^+^ T cells interacting with autologous moDCs as APCs. **(B,C)** Example dose-responses for naïve and memory T cells. Potency (P_15_) is determined by the concentration of peptide eliciting 15 % activation. **(D,E)** Examples of potency vs. K_D_ fitted with a power law. Fold-change in K_D_ and in potency derived from fits are shown. **(F)** Experimental protocol for producing primary human CD8^+^ T cell blasts interacting with the glioblastoma cell line U87 as APCs. **(G,H)** Example dose-responses and **(I,J)** potency vs. K_D_ plots for T cell blasts expressing the indicated TCR. **(K-L)** Comparison of the fitted discrimination power (*α*) and fitted sensitivity (*C*). Shown are means with each dot representing an independent experiment (n=3–6). (K) In grey the result of a statistical test vs. 1 is shown (p<0.0001 for naïve, memory & pooled, p=0.0002 for U87/1G4, p=0.0009 for U87/A6). 95 % CI for pooled *α* in **K** is 1.9–2.1.

We observed similar results with T cell blasts (Fig. 2F,G), which serve as an *in vitro* model for effector T cells and are commonly used in adoptive cell therapy. To independently corroborate discrimination with a second TCR, we used A6-expressing T cell blasts, and again found a graded response (Fig. 2H). However, potency for all pMHCs were lower and therefore, responses were only observed for higher affinity peptides with K_D_ <100 *µ*M (Fig. 2H, Fig. S2A,B), which we attribute to the much lower expression of the A6 TCR (Fig. S2C,D). Nonetheless, potency correlated with affinity (Fig. 2I,J).

In order to quantify discrimination and sensitivity we fitted the following power law to the data,

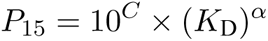

where *C* measures antigen sensitivity (y-intercept on the log-log plot) as the potency of a pMHC with K_D_ = 1 *µ*M (lower *C* values indicate higher sensitivity), and *α* measures the discrimination power (slope on the log-log plot) as it quantifies the ability of a surface receptor to amplify changes in ligand affinity into potentially larger changes in ligand potency. Mechanistically, a receptor occupancy model, where the response is proportional to the concentration of receptor/ligand complexes, produces *α* = 1 (termed baseline discrimination as there is no amplification) whereas additional mechanisms are required to produce *α >* 1 (termed enhanced discrimination). We observed enhanced discrimination powers (1.8–2.1) that were similar for naïve, memory, and blasted T cells and for both the 1G4 and A6 TCRs (Fig. 2K), and when using IL-2 as a measure of T cell activation (Fig. S3A-C). Despite these similar discrimination powers, we observed large *∼*1,000-fold variation in antigen sensitivity (Fig. 2L).

Taken together, while we found that the discriminatory power of the TCR was enhanced above baseline, we did not observe the previously reported sharp affinity threshold indicative of near-perfect discrimination.

### Systematic analysis reveals that the discriminatory power of the TCR is imperfect

Since *α* is a dimensionless measure of discrimination, we used it to compare the discriminatory power measured in this study with the apparently near-perfect discrimination suggested by earlier studies. We began by analysing the original three murine TCRs (Fig. 3A-C). In the case of the OT-I TCR (Fig. 3A), the T cell response was measured by target cell killing (9) and we defined potency as the peptide concentration producing 10 % lysis (P_10_) in order to include the E1 peptide variant. The original binding data was provided in a subsequent study (11). A plot of potency over K_D_ revealed a very large discriminatory power (*α* = 10.5), which reflects their finding that the E1 peptide variant had a 5 10^6^-fold lower potency despite apparently having only a 3.5-fold lower affinity compared to the wild-type OVA peptide. We found similar large values of *α* (12, 18, and *>* 5.1) for OT-I when using functional data from other studies (10, 15) (Table S5 ID 1-4).

**Figure 3:**
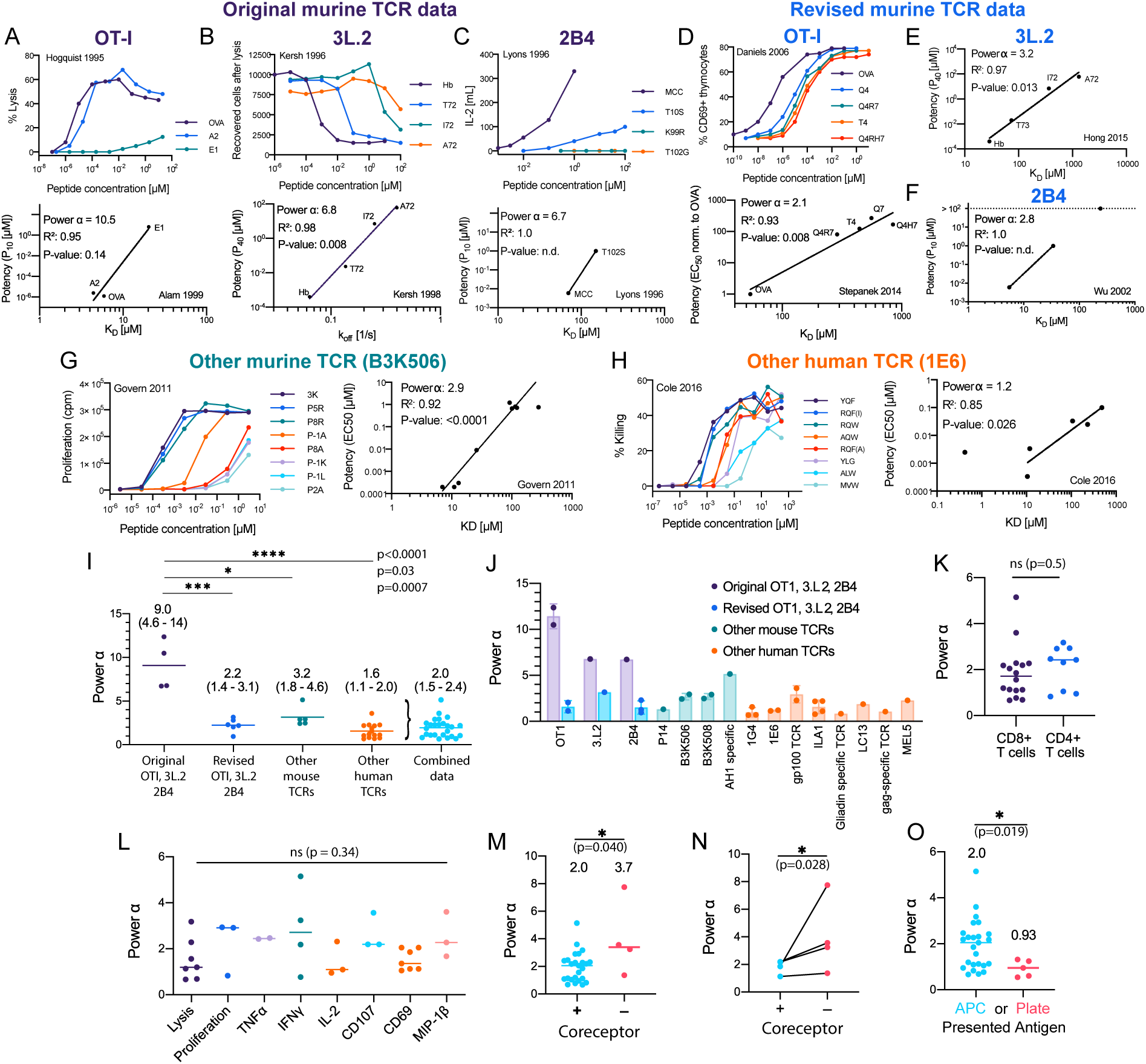
Systematic analyses shows enhanced but imperfect discriminatory powers for the TCR that depend on the antigen presenting surface. **(A-H)** T cell dose-responses and potency/affinity plots for **(A-C)** the original murine TCR data, revised analysis of the original murine TCRs using **(D)** new functional and binding data or **(E,F)** only new binding data, and examples of other **(G)** murine and **(H)** human TCRs. The highest affinity peptide (K_D_ *<* 1 *µ*M) for the 1E6 TCR was excluded because it saturated the response and would have artificially lowered the fitted *α* (see Methods for inclusion and exclusion criteria). Additional information on each panel is provided in Table S5 (ID: 2 **(A)**, 11 **(B)**, 14 **(C)**, 5 **D)**, 13 **(E)**, 17 **(F)**, 23 **(G)**, and 42 **(F)**). **(I)** Comparison of discrimination powers with mean and 95% CI (Combined data includes revised OTI, 3L.2, and 2B4 and other mouse and human data). **(J)** Discrimination powers shown in **(I)** parsed into each TCR. **(K)** Comparison between CD4^+^ and CD8^+^ T cells. **(L)** Comparison between different T cell responses. **(M)** Comparison between conditions with and without the CD4/CD8 co-receptors. **(N)** Comparison as in **(M)** but for paired data (where both conditions were present in the same study). **(O)** Comparison between the use of APCs or artificial plate surfaces to present antigens. Combined data is used in **(K,L)**, in **(M)** (+coreceptor), and **(O)** (APC data). Complete list of all 70 calculated powers can be found in Table S5 and Fig. S4-6.

Similar to OT-I, the original data for the 3.L2 (12, 13) and 2B4 (14) TCRs also produced large powers (Fig. 3B,C). In the case of 3.L2, we plotted potency over *k*_off_ instead of K_D_ because *k*_on_ was different between pMHCs (13) (Fig. 3B, bottom). Because of the small number of data points for these TCRs the correlation plots used to determine *α* only reached statistical significance (p*<*0.05) for the 3L.2 TCR. Notwithstanding this limitation, this analysis supports the conclusions of these early mouse studies that TCR discrimination was near-perfect, with *α ∼* 9 (see below).

The OT-I, 3L.2, and 2B4 transgenic mice continue to be instrumental in studies of T cell immunity and as such, substantial data has been generated relating to these TCRs over the years, including new TCR/pMHC binding measurements. Revised SPR data for OT-I revealed no binding for the E1 peptide variant (24) and therefore, we could not use the original potency data. To produce an estimate of *α* for OT-I, we combined measurements of antigen potency (29) and binding (24) that were now available for four peptides and found an appreciably lower discrimination power of 2.1 (Fig. 3D). In the case of the 3.L2 TCR, revised SPR data for the original 4 peptide variants showed a wider variation in K_D_ than originally reported (30). We re-plotted the original potency data over the revised K_D_ values (as *k*_off_ was not available for all peptides), and found a lower power of 3.2 (Fig. 3E). Similarly, re-plotting the 2B4 TCR potency data over revised binding data (31) produced a lower discrimination power of 2.8 (Fig. 3F). Although this calculation included only two data points, we identified two additional studies with 4-5 data points (32, 33) that also produced lower powers of 2.3 and 0.95 for 2B4 (Table S5 ID 18 and 19).

Thus, estimates of discrimination powers of the OT-I, 3L.2, and 2B4 TCRs based on the early binding data were much higher (mean value of *α* ~ 9) than those obtained when using more recent binding data (mean value of *α* = 2.2) (Fig. 3I), with the revised estimate being similar to the values obtained in this study for two TCRs (Fig. 2K). This strongly suggests that discrepancies between the original mouse TCR data suggesting near-perfect discrimination (*α* ~ 9) and our human TCR data suggesting imperfect discrimination (*α* = 2.0) is a consequence of issues with the original SPR measurements.

Since many other mouse and human TCRs have been characterised over the past two decades we used our approach to quantitate their discrimination powers. To be included in this study, a pMHC dose-response stimulation had to have been performed so that a measure of ligand potency could be determined and monomeric TCR/pMHC binding data (K_D_ or *k*_off_) also had to be available. We used studies that relied on different peptides that bound a single TCR, studies that relied on multiple TCRs that bound the same peptide, or studies that relied on a combination of both. We generated 51 potency plots (Fig. S5-6) and extracted the discrimination power (Table S5 ID 20-70). As representative examples, we show the mouse B3K506 TCR (Fig. 3G) and the human 1E6 TCR (Fig. 3H). Strikingly, analysis of these TCRs, and other mouse and human TCRs (Fig. 3J), produced discrimination powers that were also significantly lower than those produced using the original mouse TCR data (Fig. 3I). The variability across studies was not unexpected because they were not designed to accurately estimate *α*. Variability may be a result of the limited K_D_ range and/or issues with estimating lower-affinities. Nonetheless, combining all TCR data with the exception of the original mouse TCR data produced *α* = 2.0 (95% CI of 1.5 to 2.4), in excellent agreement with our measurements. Therefore, a 5-fold decrease in affinity can be compensated for by a 25-fold increase in antigen concentration for the TCR (*α* = 2). While this is higher than the 5-fold increase in concentration required by baseline discrimination (*α* = 1), it is far lower than the unattainable 2-million-fold increase in concentration required by near-perfect discrimination (*α* = 9). Taken together, this shows that the discriminatory power of the TCR is imperfect but enhanced above baseline.

### Factors affecting the discriminatory power of T cells

We next investigated factors that might affect the TCR discriminatory power. Using the literature data, we found no significant differences between CD4^+^ or CD8^+^ T cells (Fig. 3K) or across different T cell responses (Fig. 3L), which is consistent with a TCR proximal mechanism for discrimination. When we analysed studies where CD4/CD8 co-receptor binding was abolished (34–36) we found a significant increase in the discrimination power (Fig. 3M,N), suggesting that the well-established role of coreceptors in increasing T cell sensitivity to antigen is accompanied by a decrease in discriminatory power.

We also identified studies where the antigen was presented on artificial surfaces in isolation (e.g. recombinant pMHC immobilised on plates (37–40)) and found that *α* decreased significantly from 2.0 on APCs to 0.93 on these surfaces (Fig. 3O). Using our 1G4 T cell blasts, we confirmed that the discrimination power decreased from 2.0 when antigen was presented on APCs to 1.1 when presented as recombinant pMHC on plates (Fig. 4A-F). This suggested that other factors, beyond TCR/pMHC, may required for enhanced discrimination.

**Figure 4:**
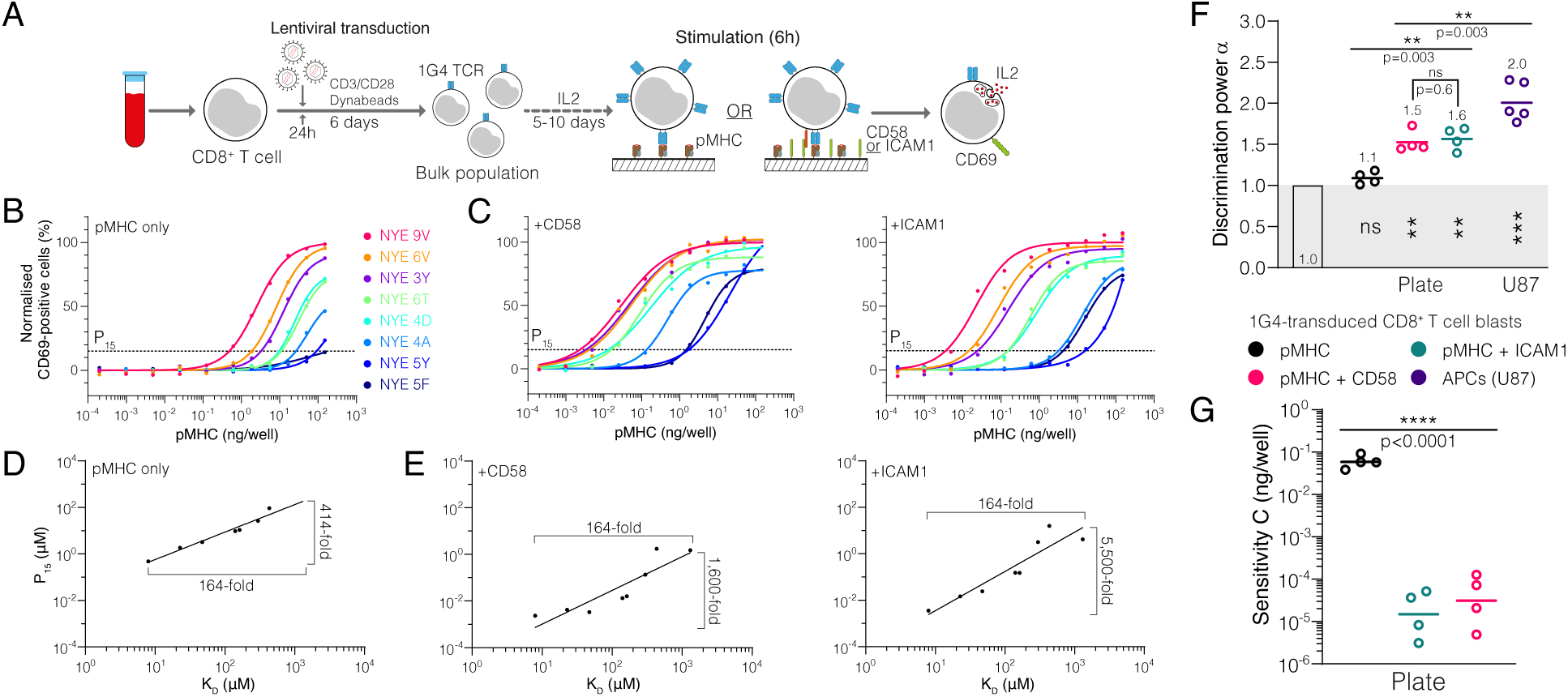
The T cell discriminatory power is enhanced by ligation of the receptors CD2 or LFA-1. **(A)** Protocol for stimulation of CD8^+^ T cell blasts with plate-bound recombinant ligands. **(B,C)** Example dose-response curve for 1G4 T cell blasts stimulated with **(B)** pMHC alone or **(C)** in combination with CD58 or ICAM1. **(D,E)** Potency derived from dose-response curves over K_D_ showing the power function fit **(D)** with pMHC alone or **(E)** in combination with CD58 or ICAM1. **(F)** Comparison of the fitted discrimination power (*α*) and fitted sensitivity (*C*). Shown are geometric means with each dot representing an independent experiment (n=4–-5). (F) In grey the result of a statistical test vs. 1 is shown (p=0.09 for pMHC, p=0.002 for CD58 & ICAM1, p=0.0002 for U87/1G4).

We hypothesised that co-signalling receptors CD2 and LFA-1 may be such factors because of their role in increasing ligand potency (41, 42). Indeed, addition of recombinant ICAM1 (a ligand of LFA-1) or CD58 (the ligand to CD2) increased TCR downregulation (Fig. S7) and antigen potency (Fig. 4C) in this experimental system, consistent with previous reports using APCs (41, 42). The potency plots highlighted that the 164-fold variation in K_D_ was now amplified into a *>* 1600-fold variation in potency (Fig. 4E) compared to only 414-fold when antigen was presented in isolation (Fig. 4D). This is reflected in the discrimination power, which increased from 1.1 to *>* 1.5 (Fig. 4F). We noted that the 100-fold increase in antigen sensitivity is appreciably larger than previous reports (41, 42) and likely reflects the reductionist system we have used where other co-signalling receptors cannot compensate (Fig. 4G). These observations were reproduced using IL-2 as a measure of T cell activation (Fig. S3). Therefore, engagement of the co-signalling receptors CD2 and LFA-1 enhance not only antigen sensitivity but also discrimination.

### The kinetic proofreading mechanism explains the discriminatory power of T cells

The KP mechanism proposes that a sequence of biochemical steps between initial pMHC binding (step 0) and TCR signalling (step N) introduces a proofreading time-delay that tightly couples TCR signalling to the *k*_off_ (or equivalently to K_D_ if *k*_on_ does not vary appreciably) of TCR/pMHC interactions (Fig. 5A). Despite being introduced more than 20 years ago (16) and underlying all models of T cell activation (43), there are no estimates for two crucial parameters in the model, namely the number of steps and the time delay for T cells discriminating antigens using APCs.

**Figure 5:**
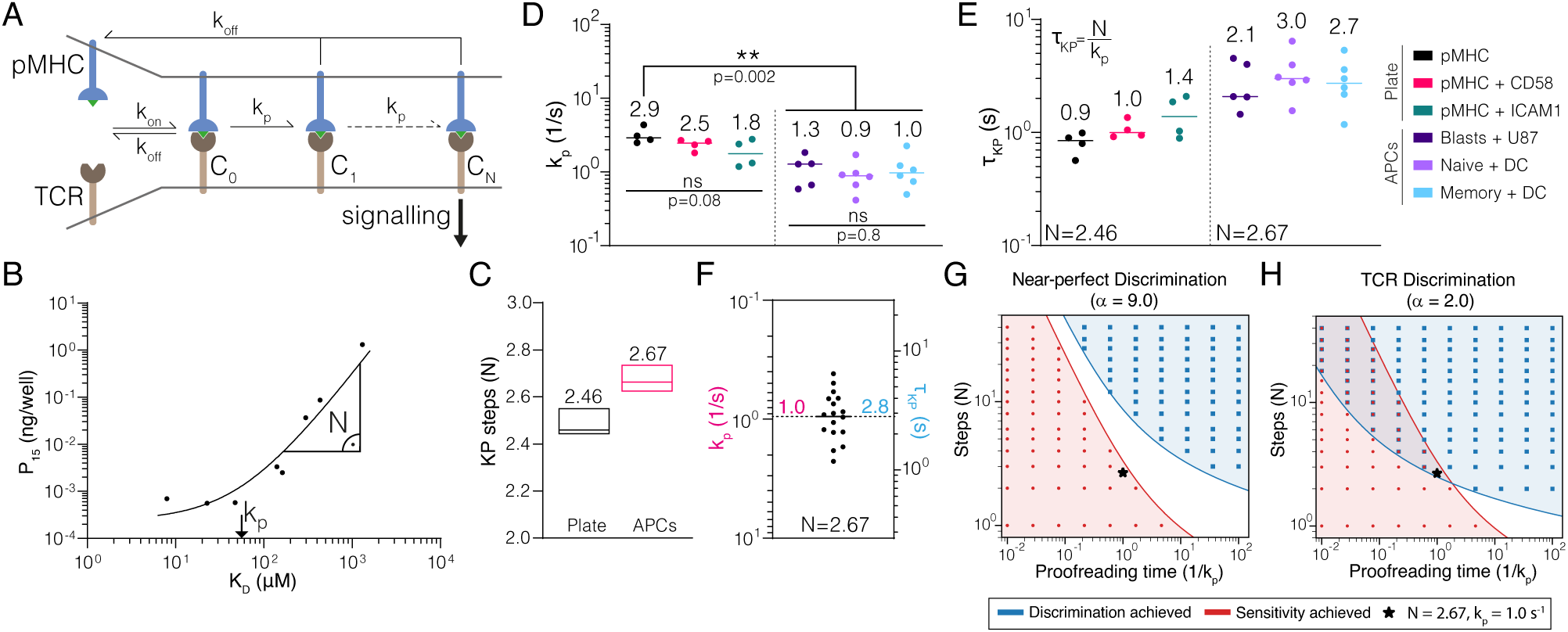
The kinetic proofreading mechanism explains TCR discrimination. **(A)** Schematic of the KP model. The KP time delay between initial binding (step 0) and signalling (step N) is *τ_KP_* = *N* /*k_p_*. **(B)** Example fit of the KP model to data generated using CD8^+^ blasts stimulated with pMHC + ICAM1 showing that the fitted *k_p_* is near the K_D_ threshold where potency saturates and *N* is the slope away from this saturation point. **(C)** The fitted number of steps (median with min/max) was a global shared parameter for all plate or APC experiments. **(D)** The fitted KP rate was a local parameter for individual experiments. **(E)** The KP time delay calculated from N in (C) and individual *k*_p_ values in (D). **(F)** Pooled APC data are used to compute means of *k_p_* and *τ*_KP_ of 1.0 (95 % CI: 0.7–1.2) and 2.8 (95 % CI: 2.2–3.6), respectively. **(G,H)** Binary heatmaps showing when sensitivity (red) and discrimination (blue) are achieved for the indicated discrimination power. Results shown using stochastic simulations (dots) or deterministic calculations (continuous colours).

To determine the KP parameters we fit the model simultaneously to all potency data from the plate experiments (27 parameters fitted to 12 experiments with a total of 89 data points) or all potency data from the APC experiments (37 parameters fitted to 17 experiments with a total of 126 data points). In both fits, we found excellent agreement (e.g. Fig. 5B, Fig. S8A,B) and, importantly, the fit method showed that *N* and *k_p_* could be uniquely determined (Fig. S8C-H). The value of *k_p_* was related to the K_D_ value where potency saturated (i.e. showed no or modest changes as K_D_ decreased) whereas the value of *N* was the slope at much larger K_D_ values (Fig. 5B). Accurately determining both parameters required potency data spanning saturation to near complete loss of responses, which can only be achieved by having a wide range of pMHC affinities down to very low affinities (high K_D_). We found an unexpectedly small number of biochemical steps when fitting the APC data (2.67), and a similar value when independently fitting the plate data (Fig. 5C). The fitted *k_p_* values were similar within the APC experiments but generally smaller than the plate experiments (Fig. 5D), and because a similar number of steps were observed in both, this translated to the time delay which was longer on APCs (Fig. 5E). Therefore, the higher discrimination power observed on APCs compared to the plate (Fig. 4F) is a result of a longer time delay produced not by more steps but rather a slower rate for each step. This made conceptual sense because the number of steps is constrained by the signalling architecture whereas the rate of each step can be regulated. We combined the similar KP parameters for the APC data to provide an average time delay of *τ*_KP_ = 2.8 s using *N* = 2.67 (Fig. 5F).

Although the KP mechanism can explain our discrimination data, it has been previously argued that it cannot simultaneously explain the observed high sensitivity of the TCR for antigen (3, 4, 15). We systematically varied the KP model parameters and determined whether discrimination and/or sensitivity were achieved for different levels of discrimination (Fig. 5G,H). As in previous reports, we found that the KP mechanism could not simultaneously achieve sensitivity and near-perfect discrimination (Fig. 5G). However, it readily achieved sensitivity and the revised imperfect discrimination that we now report and interestingly, the 2.67 steps that we determined appears to be near the minimum number required to achieve this (Fig. 5H). This may reflect the importance of maintaining a very short time-delay so that antigen recognition can proceed rapidly allowing individual T cells to rapidly scan many APCs (3, 4, 15).

### The discriminatory power of the TCR is higher than conventional surface receptors

Our finding that the discriminatory power of the TCR is only modestly enhanced above baseline raises the important question of whether it is unique in its ligand discrimination abilities. To answer this question, we identified studies that allowed us to estimate the discrimination power for Cytokine Receptors, Receptor-Tyrosine-Kinases (RTKs), G-protein Coupled Receptors (GPCRs), Chimeric Antigen Receptors (CARs), and B cell receptors (BCRs) (Fig. 6A-E). Out of 30 calculations, we found 21 significant correlations between potency and K_D_ (or *k*_off_) that allowed us to estimate *α* (Table S6). We found that the discrimination powers of Cytokine receptors, RTKs, GPCRs, and CARs were all at or below one, and as a group, their discrimination powers were significantly lower than the TCR (Fig. 6F). We identified only a single study for the BCR that could be used to compute *α* and report a preliminary discrimination power of 1.3, which is intermediate between the TCR and other receptors. Therefore, the TCR appears to be unique in its enhanced ligand discriminatory powers.

**Figure 6:**
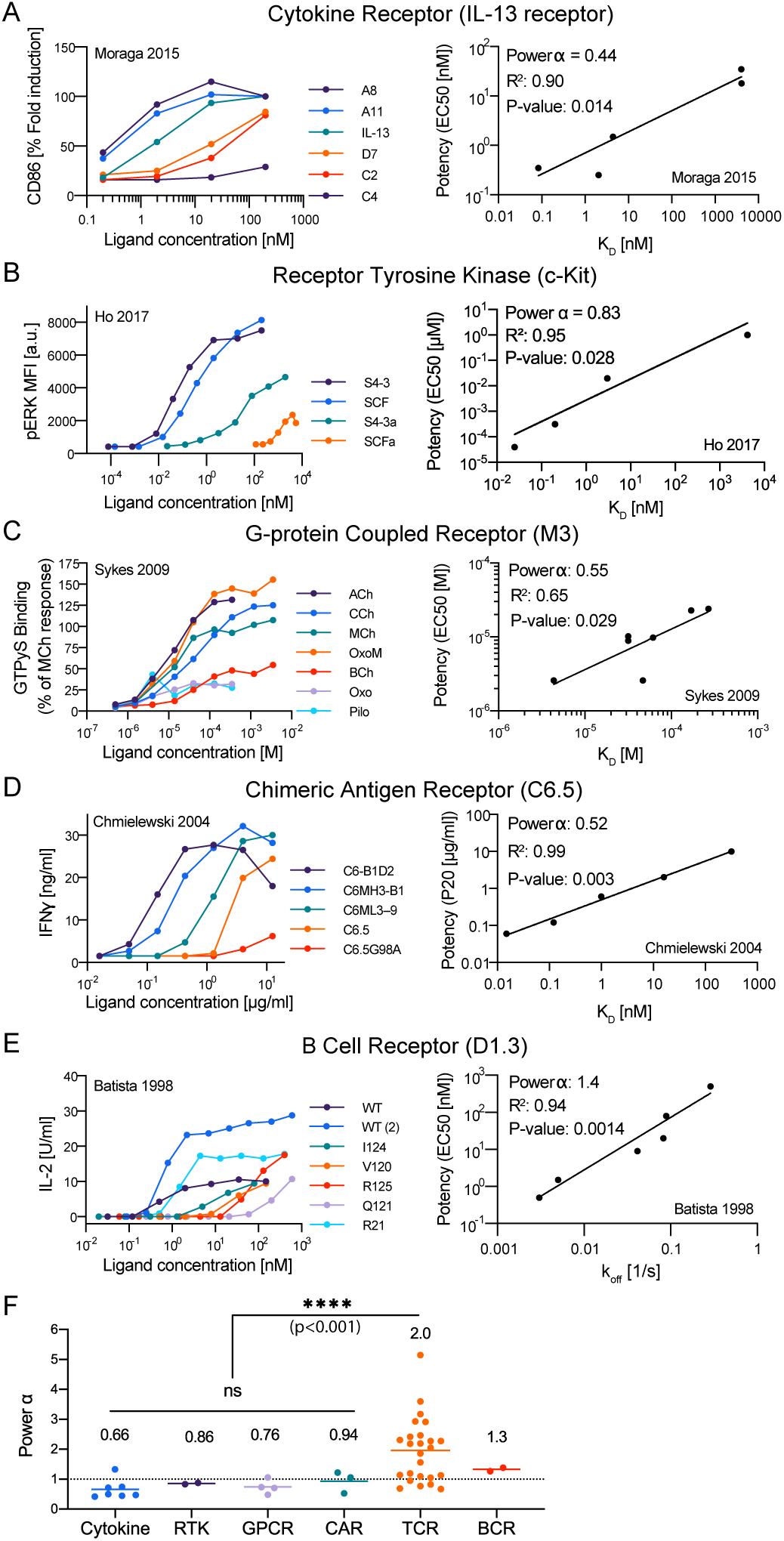
The discriminatory power of the TCR is higher than conventional surface receptors. **(A-E)** Representative dose-response (left column) and potency over K_D_ or *k*_off_ (right column) for the indicated surface receptor. **(F)** Discrimination powers for the indicated receptor. Data for the TCR as in Fig. 3I (Combined data) and data for other receptors are summarised in Table S6 and Fig. S9 (ID: 5 **(A)**, 15 **(B)**, 20 **(C)**, 25 **(D)**, 29 **(E)**).

## Discussion

In contrast to the prevailing view that the TCR exhibits near-perfect discrimination, we have shown here that the discriminatory power of the TCR is imperfect and that it is able to respond to ultra-low affinity antigens. Our estimates of TCR discrimination were facilitated by the development of a revised SPR method to accurately measure TCR/pMHC affinities.

The KP mechanism was able to explain both the high antigen sensitivity and the discrimination power of the TCR. This was achieved by a few steps (2.67) and a short proofreading time delay (2.8 s). This time delay is at the shorter end of the value estimated using pMHC tetramers (8 s with 95 % CI: 3–19 s) (44) and consistent with the 4 s time delay between pMHC binding and LAT phosphorylation (45). The small number of steps is reasonable because, although the TCR complex undergoes a large number of biochemical modifications (4), only those that must be sequential contribute. It follows that multiple ITAMs acting in parallel would not extend the proofreading chain. In support of this, the number of steps we estimate here for the TCR with 10 ITAMs is the same as the number recently reported for a CAR with 6 ITAMs (2.7 ± 0.5) (46).

The finding that the number of KP steps is fractional (2.67) may suggest that at least one intermediate proofreading step is not instantly reversible. For example, a proofreading chain with 3 steps where the 1st step can be sustained after ligand unbinding would generate a population of TCRs that required only 2 steps before productive signalling. Depending on the relative concentration of this TCR population, the apparent number of steps can be between 3 and 2. Therefore, the fractional number of steps that we have observed suggests that one (or more) KP steps may be sustained upon pMHC unbinding, which may represent the time delay between pMHC unbinding and the dephosphorylation of the TCR signalling complex and/or the unbinding of ZAP70 (47, 48).

Our finding that the discriminatory power of the TCR is enhanced compared with conventional receptors raises the question as to the underlying mechanism. One distinct feature of the TCR is that recognition occurs at a cell-cell interface and is assisted by co-signalling receptors such as CD2 and LFA-1, which appear to be required for enhanced discrimination. Our preliminary observation that the BCR may also exhibit enhanced discrimination suggests a role for ITAM-based signalling in enhanced discrimination. While our finding that ITAM-based CARs did not exhibit enhanced discrimination argues against this, CARs are artificial chimeric molecules with defects in ITAM signalling (49).

Although ligand potency usually correlates with solution or three-dimensional (3D) affinity measured by SPR, there are occasional exceptions. In one example, a structural explanation was provided for a pMHC that could bind the TCR but could not activate T cells; it exhibited an unusual docking geometry that prevented co-receptor binding (50). In another example, it was suggested that mechanical forces could affect the TCR binding affinity to different ligands in a different way (2, 51). Finally, in a third example, it was shown that the surface or two-dimensional (2D) TCR/pMHC binding parameters measured within the T cell contact interface predicted the T cell response more accurately compared to the 3D binding parameters measured in SPR (52). However, this was based on the earlier inaccurate SPR data for the OT-I system, which was the only data available at the time. A subsequent study found that the 2D and 3D binding parameters for the 1E6 TCR were equally accurate at predicting the T cell response (53). Taken together, these studies suggest that there are likely to be occasional exceptions where 3D binding properties do not correlate with potency. This may partly explain the lack of correlation between potency and 3D affinity reported in a subset of the published studies we have analysed (Table S5).

We found that the basic KP mechanism was sufficient to accurately capture antigen discrimination within the physiological affinity range and when antigens are presented in the context of self pMHCs on autologous APCs. However, it is known that the basic KP mechanism alone cannot explain the phenomena of antagonism or optimal affinity. Antagonism is a phenomena where lower-affinity pMHCs, which do not induce T cell responses on their own, are able to inhibit T cell activation by agonist pMHCs (15, 54–56). This can be explained by augmenting the KP mechanism with feedbacks (1, 15, 43). In studies that used supra-physiological TCR/pMHC affinities, it was observed that T cell responses eventually decreased as the affinity increased (39, 57–59). This optimal pMHC affinity can be explained by augmenting the KP mechanism with limited signalling (43). In the future, including data using supra-physiological and/or antagonist antigens can be used to calibrate a KP model augmented with limited signalling and/or feedbacks.

To study discrimination, we have introduced the discriminatory power (*α*) because it can quantify discrimination, independently from antigen sensitivity, from experimental studies. Previously, the term specificity has been used to refer to this discriminatory concept (1, 3, 8, 15). However, specificity is also commonly used to mean the opposite of promiscuity (i.e. the ability of T cells to respond to many different peptides). To avoid ambiguity, we suggest that specificity and promiscuity are used to refer to the tolerance of peptide sequence diversity while discrimination is used to refer to the tolerance of changes in TCR/pMHC binding parameters. Using this terminology, our analysis suggests that coreceptors decrease the discriminatory power of the TCR (Fig. 3M,N), whereas published data has demonstrated that coreceptors can increase the promiscuity of the TCR (60).

The imperfect discriminatory power of the TCR has important functional consequences. Under the assumption of near-perfect TCR discrimination, T-cell mediated autoimmunity is often viewed as a defect in thymic negative selection and/or peripheral tolerance mechanisms (19). However, with an imperfect discriminatory power of *α* = 2, the 10-100 fold lower affinity reported for autoreactive TCRs binding their self antigens (19, 20) means that they can become activated if their self antigens increase in expression by 100-10000 fold. This suggests that T-cell autoimmunity can arise by inappropriate increases in expression of self antigens, and such increases have recently been implicated in T-cell mediated autoimmunity (21, 22). T cells also have important roles in eliminating tumour cells but their therapeutic use is often limited by toxicities to lower-affinity off-tumour antigens (e.g. (61)). The factors we have identified that control antigen discrimination, together with proposed mechanisms that can generate near-perfect discrimination (1, 3, 8, 15, 62), may enable the engineering of T cells with improved discriminatory powers that selectively reduce responses to lower-affinity off-tumour antigens.

## Acknowledgements

We thank Ignacio Moraga Gonzalez, David K. Cole, David R. Greaves, Philipp Kruger, Edward Jenkins, Marcus Bridge, Samuel A. Isaacson, Marion H. Brown, and Tal Arnon for helpful discussions.

## Author contribution

Conceptualisation: EAS, MK, OD, PAvdM, JP. Investigation: JP, EAS, MK, DBW, AH. Methodology: MK, EAS, JP. Visualisation: AH, JP. Formal analysis: AH, JP, DBW. Data curation: AH, EAS, JP, DBW. Project administration: OD, JP, EAS. Funding acquisition: EAS, PAvdM, SJD, OD, MLD, JP. Writing - original draft: OD, JP, AH. Writing - review & editing: everyone. Resources: MK. Supervision: OD, PAvdM, MLD, SJD.

## Materials & Methods

### 1 Key resources

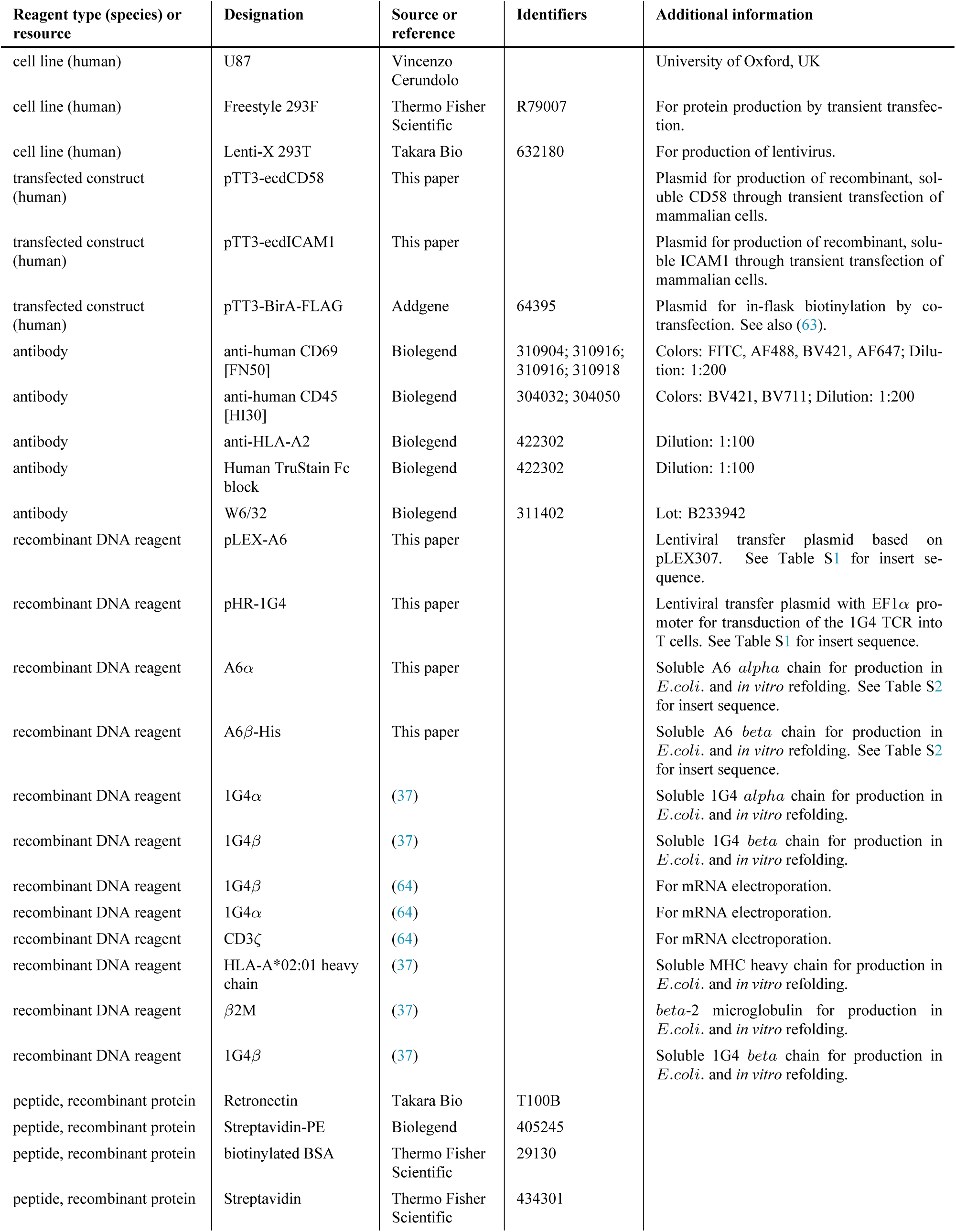
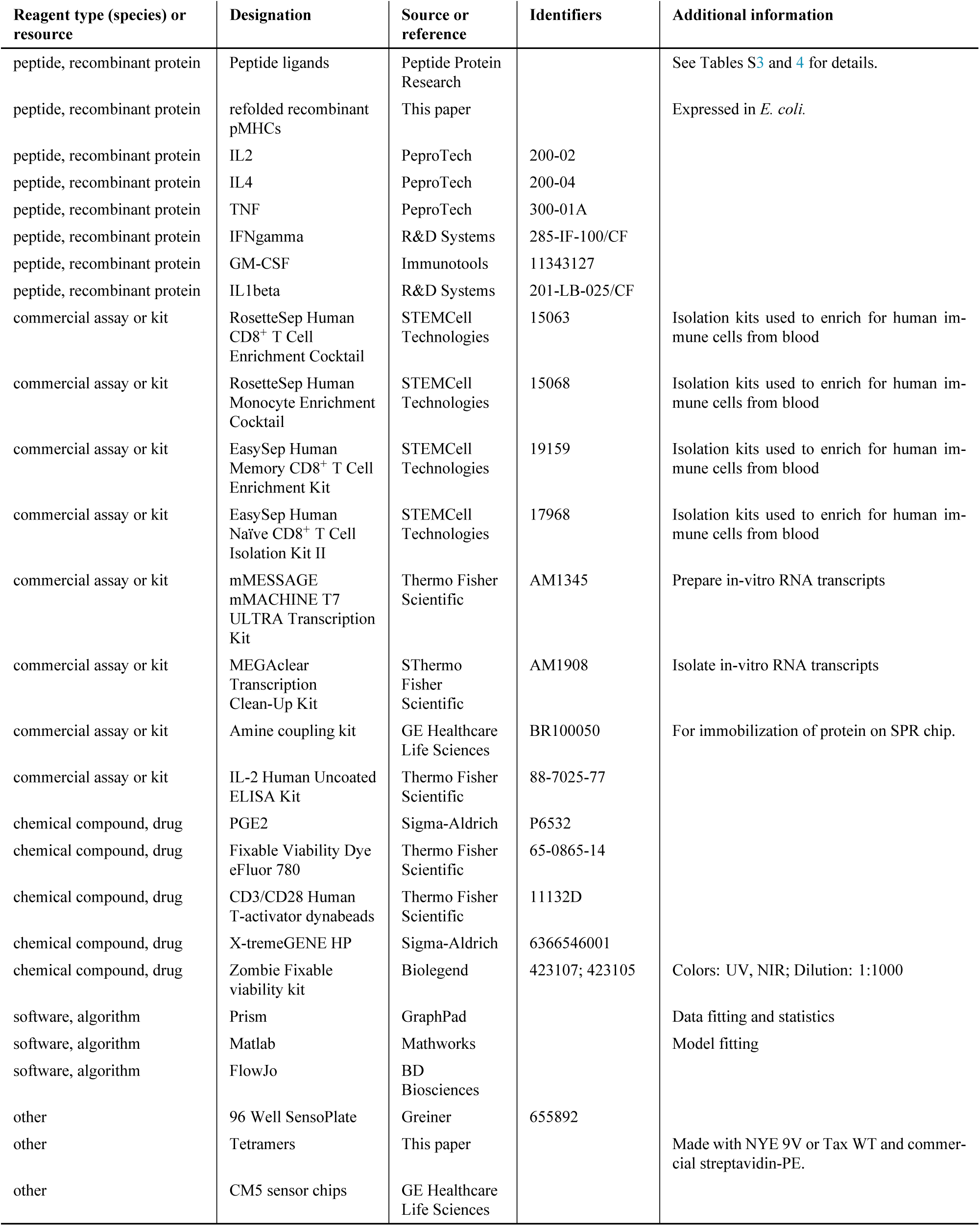

| Reagent type (species) or resource | Designation | Source or reference | Identifiers | Additional information |
| --- | --- | --- | --- | --- |
| cell line (human) | U87 | Vincenzo Cerundolo |  | University of Oxford, UK |
| cell line (human) | Freestyle 293F | Thermo Fisher Scientific | R79007 | For protein production by transient transfection. |
| cell line (human) | Lenti-X 293T | Takara Bio | 632180 | For production of lentivirus. |
| transfected construct (human) | pTT3-ecdCD58 | This paper |  | Plasmid for production of recombinant, soluble CD58 through transient transfection of mammalian cells. |
| transfected construct (human) | pTT3-ecdICAM1 | This paper |  | Plasmid for production of recombinant, soluble ICAM1 through transient transfection of mammalian cells. |
| transfected construct (human) | pTT3-BirA-FLAG | Addgene | 64395 | Plasmid for in-flask biotinylation by co-transfection. See also (63). |
| antibody | anti-human CD69 [FN50] | Biolegend | 310904; 310916; 310916; 310918 | Colors: FITC, AF488, BV421, AF647; Dilution: 1:200 |
| antibody | anti-human CD45 [HI30] | Biolegend | 304032; 304050 | Colors: BV421, BV711; Dilution: 1:200 |
| antibody | anti-HLA-A2 | Biolegend | 422302 | Dilution: 1:100 |
| antibody | Human TruStain Fc block | Biolegend | 422302 | Dilution: 1:100 |
| antibody | W6/32 | Biolegend | 311402 | Lot: B233942 |
| recombinant DNA reagent | pLEX-A6 | This paper |  | Lentiviral transfer plasmid based on pLEX307. See Table S1 for insert sequence. |
| recombinant DNA reagent | pHR-1G4 | This paper | | Lentiviral transfer plasmid with EF1α promoter for transduction of the 1G4 TCR into T cells. See Table S1 for insert sequence. |
| recombinant DNA reagent | A6α | This paper | | Soluble A6 *alpha* chain for production in *E.coli*. and *in vitro* refolding. See Table S2 for insert sequence. |
| recombinant DNA reagent | A6β-His | This paper | | Soluble A6 *beta* chain for production in *E.coli*. and *in vitro* refolding. See Table S2 for insert sequence. |
| recombinant DNA reagent | 1G4α | (37) | | Soluble 1G4 *alpha* chain for production in *E.coli*. and *in vitro* refolding. |
| recombinant DNA reagent | 1G4β | (37) | | Soluble 1G4 *beta* chain for production in *E.coli*. and *in vitro* refolding. |
| recombinant DNA reagent | 1G4β | (64) | | For mRNA electroporation. |
| recombinant DNA reagent | 1G4α | (64) | | For mRNA electroporation. |
| recombinant DNA reagent | CD3ζ | (64) | | For mRNA electroporation. |
| recombinant DNA reagent | HLA-A*02:01 heavy chain | (37) |  | Soluble MHC heavy chain for production in *E.coli*. and *in vitro* refolding. |
| recombinant DNA reagent | β2M | (37) | | *beta*-2 microglobulin for production in *E.coli*. and *in vitro* refolding. |
| recombinant DNA reagent | 1G4β | (37) | | Soluble 1G4 *beta* chain for production in *E.coli*. and *in vitro* refolding. |
| peptide, recombinant protein | Retronectin | Takara Bio | T100B |  |
| peptide, recombinant protein | Streptavidin-PE | Biolegend | 405245 |  |
| peptide, recombinant protein | biotinylated BSA | Thermo Fisher Scientific | 29130 |  |
| peptide, recombinant protein | Streptavidin | Thermo Fisher Scientific | 434301 |  |
| peptide, recombinant protein | Peptide ligands | Peptide Protein Research |  | See Tables S3 and 4 for details. |
| peptide, recombinant protein | refolded recombinant pMHCs | This paper |  | Expressed in *E. coli*. |
| peptide, recombinant protein | IL2 | PeproTech | 200-02 |  |
| peptide, recombinant protein | IL4 | PeproTech | 200-04 |  |
| peptide, recombinant protein | TNF | PeproTech | 300-01A |  |
| peptide, recombinant protein | IFNgamma | R&D Systems | 285-IF-100/CF |  |
| peptide, recombinant protein | GM-CSF | Immunotools | 11343127 |  |
| peptide, recombinant protein | IL1beta | R&D Systems | 201-LB-025/CF |  |
| commercial assay or kit | RosetteSep Human CD8<sup>+</sup> T Cell Enrichment Cocktail | STEMCell Technologies | 15063 | Isolation kits used to enrich for human immune cells from blood |
| commercial assay or kit | RosetteSep Human Monocyte Enrichment Cocktail | STEMCell Technologies | 15068 | Isolation kits used to enrich for human immune cells from blood |
| commercial assay or kit | EasySep Human Memory CD8<sup>+</sup> T Cell Enrichment Kit | STEMCell Technologies | 19159 | Isolation kits used to enrich for human immune cells from blood |
| commercial assay or kit | EasySep Human Naïve CD8<sup>+</sup> T Cell Isolation Kit II | STEMCell Technologies | 17968 | Isolation kits used to enrich for human immune cells from blood |
| commercial assay or kit | mMESSAGE mMACHINE T7 ULTRA Transcription Kit | Thermo Fisher Scientific | AM1345 | Prepare in-vitro RNA transcripts |
| commercial assay or kit | MEGAclear Transcription Clean-Up Kit | SThermo Fisher Scientific | AM1908 | Isolate in-vitro RNA transcripts |
| commercial assay or kit | Amine coupling kit | GE Healthcare Life Sciences | BR100050 | For immobilization of protein on SPR chip. |
| commercial assay or kit | IL-2 Human Uncoated ELISA Kit | Thermo Fisher Scientific | 88-7025-77 |  |
| chemical compound, drug | PGE2 | Sigma-Aldrich | P6532 |  |
| chemical compound, drug | Fixable Viability Dye eFluor 780 | Thermo Fisher Scientific | 65-0865-14 |  |
| chemical compound, drug | CD3/CD28 Human T-activator dynabeads | Thermo Fisher Scientific | 11132D |  |
| chemical compound, drug | X-tremeGENE HP | Sigma-Aldrich | 6366546001 |  |
| chemical compound, drug | Zombie Fixable viability kit | Biologend | 423107; 423105 | Colors: UV, NIR; Dilution: 1:1000 |
| software, algorithm | Prism | GraphPad |  | Data fitting and statistics |
| software, algorithm | Matlab | Mathworks |  | Model fitting |
| software, algorithm | FlowJo | BD Biosciences |  |  |
| other | 96 Well SensoPlate | Greiner | 655892 |  |
| other | Tetramers | This paper |  | Made with NYE 9V or Tax WT and commercial streptavidin-PE. |
| other | CM5 sensor chips | GE Healthcare Life Sciences |  |  |

### 2 Protein production

Class I pMHCs were refolded as previously described (65). Human HLA-A*0201 heavy chain (UniProt residues 25–298) with a C-terminal AviTag/BirA recognition sequence and human beta-2 microgolublin were expressed in *Escherichia coli* and isolated from inclusion bodies. Trimer was refolded by consecutively adding peptide, *β*2M and heavy chain into refolding buffer and incubating for 2–3 days at 4*^◦^*C. Protein was filtered, concentrated using centrifugal filters, biotinylated (BirA biotin-protein ligase bulk reaction kit [Avidity, USA]) and purified by size exclusion chromatography (Superdex 75 column [GE Healthcare]) in HBS-EP (0.01 M HEPES pH 7.4, 0.15 M NaCl, 3 mM EDTA, 0.005 % v/v Tween20). Purified protein was aliquoted and stored at −80*^◦^*C until use. Soluble *α* and *β* subunits of 1G4 and A6 TCRs were produced in *E. coli*, isolated from inclusion bodies, refolded *in vitro* and purified using size exclusion chromatography in HBS-EP, as described previously (37).

Soluble extracellular domain (ECD) of human CD58 (UniProt residues 29–204 or 29–213) was produced either in Freestyle 293F suspension cells (Thermo Fisher) or adherent, stable GS CHO cell lines. For the latter, cells were expanded in selection medium (10 % dialysed FCS, 1x GSEM supplement [Sigma-Aldrich], 20–50 *µ*M MSX, 1 % Pen/Strep) for at least 1 week. Production was performed in production medium (2–5 % FCS, 1x GSEM supplement, 20 *µ*M MSX, 2 mM sodium butyrate, 1 % Pen/Strep) continuously for a few weeks with regular medium exchanges. Human ICAM1 ECD (UniProt residues 28–480) was either produced by transient transfection or lentiviral transduction of adherent 293T, or by transient expression in 293F. Production in 293F was performed according to manufacturer’s instructions using pTT3-ecdCD58 or pTT3-ecdICAM1. All supernatants were 0.45 *µ*m filtered and 100 *µ*M PMSF was added. Proteins were purified using standard Ni-NTA agarose columns, followed by *in vitro* biotinylation as described above. Alternatively, ligands were biotinylated by co-transfection (1:10) of a secreted BirA-encoding plasmid (pTT3-BirA-FLAG) and adding 100 *µ*M D-biotin to the medium, as described before (66). Proteins were further purified and excess biotin removed from proteins biotinylated *in vitro* by size exclusion chromatography (Superdex 75 or 200 column [GE Healthcare]) in HBS-EP; purified proteins were aliquoted and stored at −80*^◦^*C until use.

Biotinylation levels of pMHC and accessory ligands were routinely tested by gel shift on SDS-PAGE upon addition of saturating amounts of streptavidin.

### 3 Surface plasmon resonance

TCR–pMHC interactions were analysed on a Biacore T200 instrument (GE Healthcare Life Sciences) at 37*^◦^*C and a flow rate of 10min. Running buffer was HBS-EP. Streptavidin was coupled to CM5 sensor chips using an amino coupling kit (GE Healthcare Life Sciences) to near saturation, typically 10000–12000 response units (RU). Biotinylated pMHCs (47 kDa) were injected into the experimental flow cells (FCs) for different lengths of time to produce desired immobilisation levels (typically 500–1500 RU), which were matched as closely as feasible in each chip. Usually, FC1 was as a reference for FC2–FC4. Biotinylated CD58 ECD (24 kDa + ~ 25 kDa glycosylation) was immobilised in FC1 at a level matching those of pMHCs. In some experiments, another FC was used as a reference. Excess streptavidin was blocked with two 40 s injections of 250 *µ*M biotin (Avidity). Before injections of soluble 1G4 or A6 *αβ*TCR (51 kDa), the chip surface was conditioned with 8 injections of the running buffer. Dilution series of TCRs were injected simultaneously in all FCs; the duration of injections (30–70 s) was the same for conditioning and TCR injections. After every 2–3 TCR injections, buffer was injected to generate data for double referencing. After the final TCR injection and an additional buffer injection, W6/32 antibody (10 *µ*g/ml; Biolegend; Lot: B233942) was injected for 10 min.

TCR Steady-state binding was measured >10 s post-injection. In addition to subtracting the signal from the reference FC with immobilised CD58 (single referencing), all TCR binding data was double referenced (67) vs. the average of the closest buffer injections before and after TCR injection. This allows to exclude small differences in signal between flow cells (e.g. drifts). TCR binding versus TCR concentration was fitted with the following model: *B* = *B_max_* [*TCR*]/(*K_D_* + [*TCR*]), where *B* is the response/binding, B_max_ the maximal binding (this parameter is either kept free or is fixed with the W6/32 derived B_max_), and [*TCR*] the injected TCR concentration. Maximal W6/32 binding (R_max_) was used to generate the empirical standard curve and to infer the B_max_ of TCRs from the standard curve. R_max_ was derived by fitting the W6/32 binding data after double referencing with the following, empirically chosen, model: *R* = *R_max_∗t*/(*K_t_* +*t*), where *t* is time (s), *R* the sensogram response after single referencing, and *K_t_* a nuisance parameter. The empirical standard curve only contained data where the ratio of the highest concentration of TCR to the fitted K_D_ value (obtained using the standard method with B_max_ fitted) was 2.5 or more. This threshold ensured that the binding response curves saturated so that only accurate measurements of B_max_ were included. All interactions were fit using both the fitted and constrained B_max_ method (Fig. 1E). For constrained K_D_ above 20 *µ*M we reported the constrained K_D_, otherwise we use the B_max_ fitted K_D_. SPR data was analysed using GraphPad Prism 8 (GraphPad software) or using a custom Python script (Python v3.7 and lmfit v0.9.13).

### 4 Co-culture of naïve & memory T cells

The assay was performed as previously described (68). Naïve and memory T cells were isolated from anonymized HLA-A2^+^ leukocyte cones obtained from the NHS Blood and Transplantation service at Oxford University Hospitals by (REC 11/H0711/7), using EasySep Human naïve CD8^+^ T Cell Isolation Kit (Stemcell) and EasySep Human Memory CD8^+^ T Cell Enrichment Kit (Stemcell), respectively. Cells were washed 3x with Opti-MEM serum-free medium (Thermo Fisher) and 2.5–5.0 Mio cells were resuspended at a density of 25 Mio/ml. Suspension was mixed with 5 *µ*g/Mio cells of 1G4*α*, 1G4*β*, and CD3*ζ* each and 100–200 *µ*l suspension was transferred into a BTX Cuvette Plus electroporation cuvette (2 mm gap; Harvard Bioscience). Electroporation was performed using a BTX ECM 830 Square Wave Electroporation System (Harvard Bioscience) at 300 V, 2 ms. T cells were used 24 h after electroporation. 1G4 TCR contained an engineered cysteine (*α*T48C and *β*S57C) to reduce mispairing (69).

Autologous monocytes were enriched from the same blood product using RosetteSep Human Monocyte Enrichment Cocktail (Stemcell), cultured at 1–2 Mio/ml in 12-well plates in the presence of 50 ng/ml IL4 (PeproTech) and 100 ng/ml GM-CSF (Immunotools) for 24 h to induce differentiation. Maturation into moDCs was induced by adding 1 *µ*M PGE_2_ (Sigma Aldrich), 10 ng/ml IL1*β* (Biotechne), 20 ng/ml IFN*γ*, and 50 ng/ml TNF (PeproTech) for an additional 24 h. MoDCs (50,000/well) were loaded for 60–90 min at 37*^◦^*C with peptide and labelled with Cell Trace Violet (Thermo Fisher) to distinguish them from T cells prior to co-culturing with 50,000 T cells/well in a 96 well plate for 24 h. T cell activation was assessed by flow cytometry and testing culture supernatant for cytokines using ELISAs.

### 5 T cell blasts

All cell culture of human T cells was done using complete RPMI (10 % FCS, 1 % penicillin/streptomycin) at 37*^◦^*C, 5 % CO_2_. T cells were isolated from whole blood from healthy donors or leukocyte cones purchased from the NHS Blood and Transplantation service at the John Radcliffe Hospital. For whole blood donations, a maximum of 50 ml was collected by a trained phlebotomist after informed consent had been given. This project has been approved by the Medical Sciences Inter-Divisional Research Ethics Committee of the University of Oxford (R51997/RE001) and all samples were anonymised in compliance with the Data Protection Act.

For plate stimulations and experiments with U87 target cells, CD8^+^ T cells were isolated using Rosette-Sep Human CD8^+^ enrichment cocktail (Stemcell) at 6 *µ*l/ml for whole blood or 150 *µ*l/ml for leukocyte cones. After 20 min incubation at room temperature, blood cone samples were diluted 3.125-fold with PBS, while whole blood samples were used directly. Samples were layered on Ficoll Paque Plus (GE) at a 0.8:1.0 ficoll:sample ratio and spun at 1200 g for 20–30 min at room temperature. Buffy coats were collected, washed twice, counted and cells were resuspended in complete RMPI with 50 U/ml IL2 (PeproTech) and CD3/CD28 Human T-activator dynabeads (Thermo Fisher) at a 1:1 bead:cell ratio. Aliquots of 1 Mio cells in 1 ml medium were grown overnight in 12- or 24-well plates (either TC-treated or coated with 5 *µ*g/cm^2^ retronectin [Takara Bio]) and then transduced with VSV-pseudotyped lentivirus encoding for either the 1G4 or the A6 TCR. After 2 days (4 days after transduction), 1 ml of medium was exchanged, and IL2 was added to a final concentration of 50 U/ml. Beads were magnetically removed at day 5 post-transduction and T cells from thereon were resuspended at 1 Mio/ml with 50 U/ml IL2 every other day. For functional experiments, T cells were used between 10–16 days after transduction.

### 6 Lentivirus production

HEK 293T or Lenti-X 293T (Takara) were seeded in complete DMEM in 6-well plate to reach 60–80% confluency after one day. Cells were either transfected with 0.95 *µ*g pRSV-Rev, 0.37 *µ*g pVSV-G (pMD2.G), 0.95 *µ*g pGAG (pMDLg/pRRE), and 0.8 *µ*g of pLEX-A6 or pHR-1G4 with 9 *µ*l X-tremeGENE 9 or HP (both Roche). Lentiviral supernatant was harvested after 20–30 h and filtered through a 0.45 *µ*m cellulose acetate filter. In an updated version, LentiX cells were transfected with 0.25 *µ*g pRSV-Rev, 0.53 *µ*g pGAG, 0.35 *µ*g pVSV-G, and 0.8 *µ*g transfer plasmid using 5.8 *µ*l X-tremeGENE HP. Medium was replaced after 12–18 h and supernatant harvested as above after 30–40 h. Supernatant from one well of a 6-well plate was used to transduce 1 Mio T cells. Sequence for the A6 TCR lacked one natural cysteine per chain and included engineered cysteines (*α*T48C and *β*S57C) to reduce the formation of mixed TCR dimers with endogenous TCR (69). The 1G4 TCR was expressed from the WT sequences without engineered cysteines.

### 7 Co-culture of T cell blasts

For co-culture experiments with U87 (a kind gift of Vincenzo Cerundolo, University of Oxford), 30,000 target cells were seeded in a TC-coated 96-well F-bottom plate and incubated overnight. Peptides were diluted in complete DMEM (10 % FCS, 1 % penicillin/streptomycin) to their final concentration and incubated with U87 cells for 1–2 h at 37*^◦^*C. Peptide-containing medium was removed and 60,000 TCR-transduced primary human CD8^+^ T cell blasts were added, spun for 2 min at 50 g, and incubated for 5 h at 37*^◦^*C. At the end of the experiment, 10 mM EDTA was added and cells were detached by vigorous pipetting. Cells were stained for flow cytometry and analysed immediately, or fixed and stored for up to 1 day before running. Supernatants were saved for cytokine ELISAs.

### 8 Plate stimulation

Glass-bottom Sensoplates (96-well; Greiner) were washed with 1 M HCl/70 % EtOH, thoroughly rinsed twice with PBS and coated overnight at 4*^◦^*C with 100 *µ*l/well of 1 mg/ml biotinylated BSA (Thermo Fisher) in PBS. Plates were washed with PBS twice and incubated for at least 1 h with 20 *µ*g/ml streptavidin (Thermo Fisher) in 1 % BSA/PBS at room temperature. Plates were washed again with PBS and biotinylated pMHC (in-house) was added for at least 1 h at room temperature or overnight at 4*^◦^*C. Plates were emptied and accessory ligand (CD58 or ICAM1, in-house) or PBS was added for the same duration as above. Upon completion, plates were washed once and stored for up to 1 day in PBS at 4*^◦^*C.

For stimulation, T cells were counted, washed once to remove excess IL2, and 75,000 cells in 180–200 *µ*l complete RMPI were dispensed per well. Cells were briefly spun down at 50 g to settle to the bottom and subsequently incubated for 4 h at 37*^◦^*C. At the end of the experiment, 10 mM EDTA was added and cells were detached by vigorous pipetting. Cells were stained for flow cytometry and analysed immediately, or fixed and stored for up to 1 day. Supernatants were saved for cytokine ELISAs.

### 9 Peptides and loading

We used peptide ligands that were either described previously (37, 39, 70–74) or designed by us based on the published crystal structures of these TCRs in complex with MHC (1G4: PDB 2BNQ, A6: PDB 1AO7).

Peptides were synthesised at a purity of >95 % (Peptide Protein Research, UK). Tax WT is a 9 amino acid, class I peptide derived from HTLV-1 Tax_11–19_ (27, 75). NYE 9V refers to a heteroclitic (improved stability on MHC), 9 amino acid, class I peptide derived from the wild type NYE-ESO_157–165_ 9C peptide (26). See Table 3 and Table 4 for a list of peptides.

Loading efficiency was evaluated by pulsing T2 cells for 1–2 h at 37*^◦^*C with a titration of peptides. Loading was assessed as upregulation of HLA-A2 (clone: BB7.2; Biolegend) by flow cytometry.

### 10 Flow cytometry

Tetramers were produced in-house using refolded monomeric, biotinylated pMHC and streptavidin-PE (Biolegend) at a 1:4 molar ratio. Streptavidin-PE was added in 10 steps and incubated for 10 min while shaking at room temperature. Insoluble proteins were removed by brief centrifugation at 13,000 *g* and 0.05–0.1 % sodium azide added for preservation. Tetramers were kept for up to 3 months at 4*^◦^*C. Cells were stained for CD69 with clones FN50 (Biolegend). Staining for CD45 (clone HI30; Biolegend) was used to distinguish target and effector cells in co-culture assays with U87 cells. Cell viability staining was routinely performed for plate stimulations and U87 co-culture using fixable violet or near-infrared viability dyes (Zombie UV fixable viability kit [Biolegend], Zombie NIR fixable viability kit [Biolegend], eBioscience fixable viability dye eFluor 780 [Invitrogen]). Samples were analysed using a BD X-20 flow cytometer and data analysis was performed using FlowJo v10 (BD Biosciences).

### 11 ELISAs

Human IL-2 Ready-SET Go! ELISA kit (eBioscience/Invitrogen) or Human TNF alpha ELISA Ready-SET-Go! (eBioscience/Invitrogen) and Nunc MaxiSorp 96-well plates (Thermo Fisher) were used according to the manufacturer’s instructions to test appropriately diluted (commonly 4–30-fold) T cell supernatant for secretion of IL2 or TNF.

### 12 TCR expression

TCR*αβ*-KO Jurkat E6.1 cells (a kind gift of Edward Jenkins) were transduced with 1G4 or A6 lentivirus and TCR expression was measured by staining for CD3 (clone: UCHT1; Biolegend) and TCR*αβ* (clone IP26; Biolegend).

### 13 Data analysis

Quantitative analysis of antigen discrimination was performed by first fitting dose-response data with a 4-parameter sigmoidal model on a linear scale in Python v3.7 and lmfit v0.9.13 using Levenberg–Marquardt:

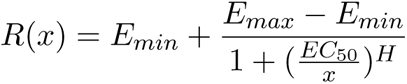

where *x* refers to the peptide concentration used to pulse the target cells (in *µ*M) or the amount of pMHC used to coat the well of a plate (in ng/well). The curve produced by this fit was used to interpolate potency as the concentration of antigen required to induce activation of 15 % for CD69 (*P*_15_) and 10 % for IL2 (*P*_10_). These percentages were chosen based on noise levels and to include lower affinity antigens in the potency plots. Potency values exceeding doses used for pulsing or coating were excluded from the analysis (i.e. no extrapolated data was included in the analysis).

To determine the discrimination power *α*, we fitted the power law in log-space to our data:

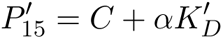

where 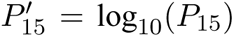 and 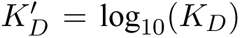. All data analysis was performed using GraphPad Prism

(GraphPad Software), if not stated otherwise.

### 14 Kinetic proofreading: fitting to data

#### Deriving the expression for ligand potency

A pMHC ligand *L* can bind with a T cell receptor *R* to create a complex *C*_0_ at a rate *k*_on_. In order for this complex to initiate TCR signalling, it undergoes a series of *N* steps. We denote by *C_i_* a TCR/pMHC complex in the *i*-th KP step. A complex *C_i_* becomes a complex *C_i_*_+1_ with rate *k_p_*, for 0 *≤ i ≤ N −* 1. At any KP step the pMHC ligand can unbind with rate *k*_off_. Let *L*(*t*), *R*(*t*), and *C_i_*(*t*) be the concentration of ligand, receptor and complex in the *i*-th KP step at time *t*, respectively. The system of ordinary differential equations that govern the temporal evolution of the concentrations is given by

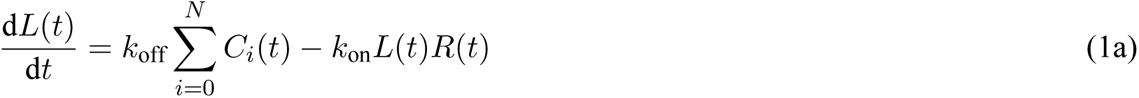

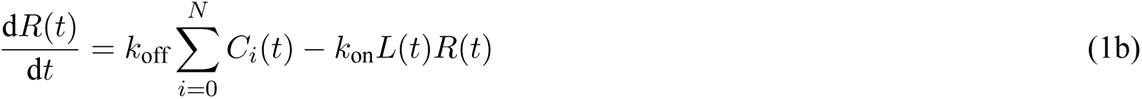

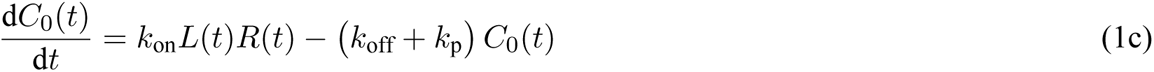

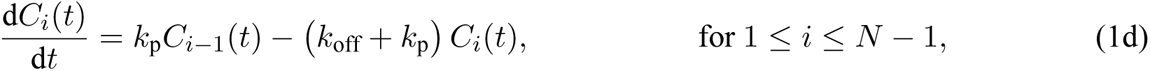

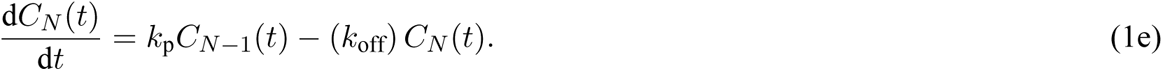

Let the initial number of pMHC ligands and T cell receptors be *L*_0_ and *R*_0_, respectively. We then define the total number of complexes at time *t* as 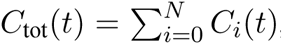, and note the two conservation equations, *L*_0_ = *L*(*t*) + *C*_tot_(*t*) and *R*_0_ = *R*(*t*) + *C*_tot_(*t*). Solving the steady state equations arising from setting the time derivatives in Eq. (1) to zero, and substituting in the conservation equations we find that

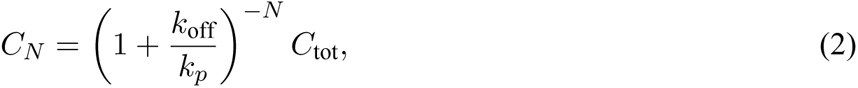

where

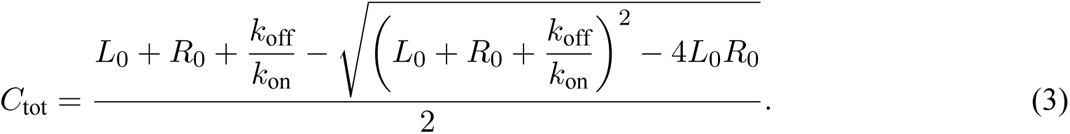

The expression in Eq. (2) determines the concentration of actively signalling TCR/pMHC complexes *C_N_* for a given number of ligands *L*_0_. To fit this model to the potency data we are interested in calculating the concentration of pMHC ligand required to initiate T cell activation for different TCR/pMHC binding parameters. We first introduce a few convenient rescalings and redefinitions. We define *x* = *L*_0_/*R*_0_ to be the potency of ligand concentration relative to the total number of receptors and let *λ* = *C_N_* /*R*_0_ be a threshold parameter that dictates how much *C_N_* complex is needed to activate a T cell response relative to the total number of receptors. Thus Eq. (2) can be rewritten as

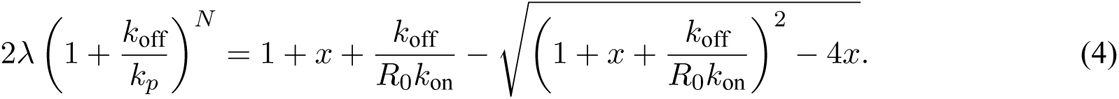

The experimental measurements of potency do not directly correspond to the potency *x* in our model as the exact number of ligand and receptor is unknown. Therefore we introduce a constant of proportionality *γ* into our model, such that *x→γx*. Similarly, the ratio *k*_off_/*k*_on_ is a measure of ligand affinity and is directly proportional to the experimental K_D_ values, thus we introduce a second constant of proportionality *δ* such that *k*_off_/(*R*_0_*k*_on_)→*δK_D_*, where we absorb the constant *R*_0_ into the new parameter. With these adjustments Equation (4) becomes

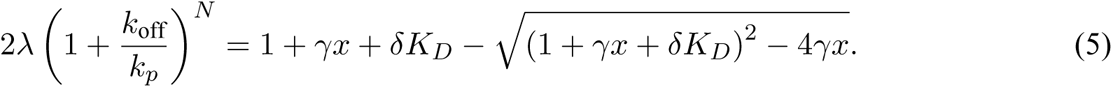

Upon rearranging Eq. (5) we find that

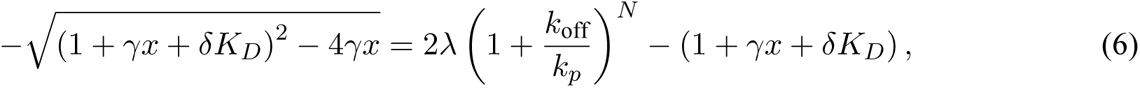

we then square^1^ both sides of Eq. (6) and find the following expression for the potency

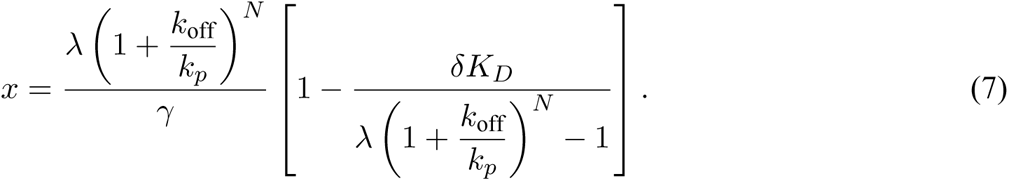

#### Fitting the potency expression using ABC-SMC parameter estimation

We used the Approximate Bayesian Computation-Sequential Monte Carlos (ABC-SMC) algorithm to determine the distribution of KP model parameters that fit the experimental data. Our KP model has five parameters, *N*, *k_p_*, *λ*, *γ* and *δ*. We fit the model parameters to the plate and the cell data separately. For both the plate and the cell data we fit *N*, *γ* and *δ* as a global parameter shared amongst all experimental repeats. The parameters *k_p_* and *λ* are fitted locally for each repeat. We fit the potency equation to the experimental data in log space and as such the log expression for potency, 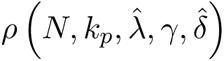, calculated from Eq. (7) is given by

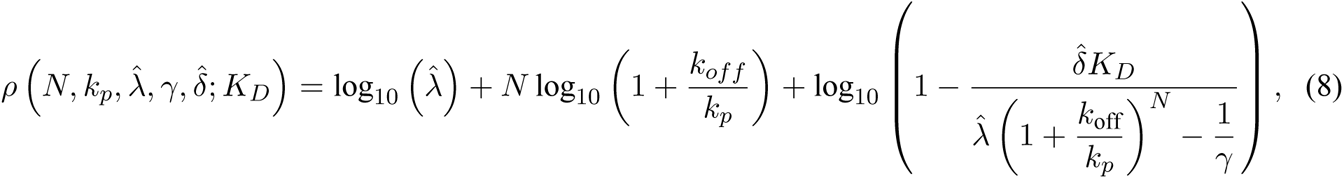

where 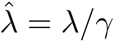 and 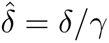 These rescalings ensure that the parameters are orthogonal and thus parameter space can be searched efficiently. The fast kinetics of the low-affinity pMHCs precluded direct measurements of *k*_off_ and instead, we noted that on-rates exhibit small variations between pMHCs that differ by few amino acids (37, 39). Therefore, we estimated *k*_off_ using K_D_ and a fixed *k*_on_ of 0.0447 *µ*M*^−^*^1^s*^−^*^1^ taken as the average *k*_on_ of NYE 9C, 9V, 3A, 3I, 3M, 3Y, and 6V previously measured at 37*^◦^*C (37).

We chose uniform prior distributions in log space for each parameter except *N*, where a uniform prior in linear space was used. This allows for efficient search through parameter space over many orders of magnitude. The priors for the plate data are as follows

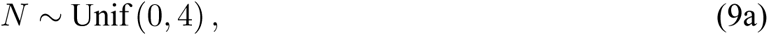

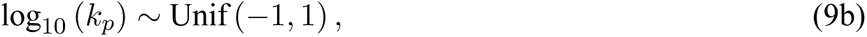

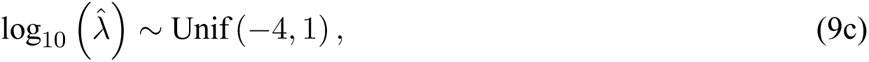

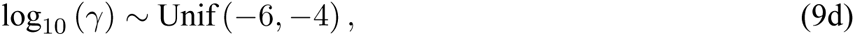

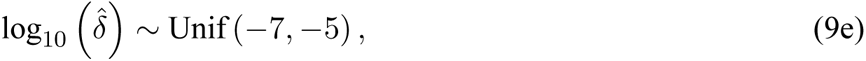

where the priors for the cell data are the same other than for 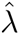 where 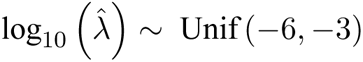.

Recall that we fit the parameters *N*, *γ*, and 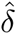 globally and 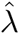 and *k_p_* are fitted locally. For the plate data this results in 27 parameters to fit whilst for the cell data there are 37 parameters. 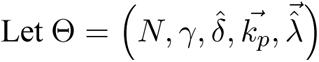 be the vector of parameters to fit such that the *i*-th entry of the vectors 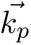 and 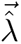 correspond to the local parameters for the *i*-th experiment. Then let 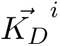 be the vector of experimentally measured K_D_ values, and 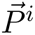 be the vector of potency measurements for the *i*-th experiment. These vectors differ in length and so we denote by *d_i_* the number of data points in the *i*-th experiment. We measure the similarity between the KP model and the experimental results via the following distance function

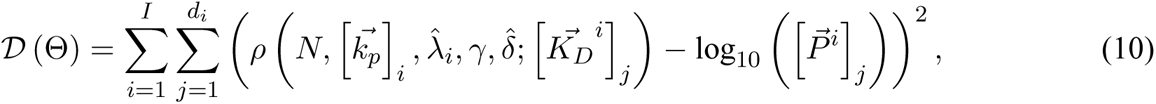

where *I* denotes the total number of experiments, *I* = 12 and *I* = 17 for the plate and cell data, respectively.

To perform a randomised search through the parameter space we employed the following Metropolis-Hastings algorithm. We sample an initial parameter set Θ_0_ from the prior distributions detailed above. Let Θ_curr_ denote the current set of parameters which initially is Θ_0_. A candidate set of parameters, Θ_cand_ is found by adding a random perturbation to Θ_curr_. The perturbation is achieved by adding a uniform random shift to each parameter in Θ_curr_ independently. The range of the uniform random shift is [−0.005, 0.005] multiplied by the width of the prior. For example we perturb the *N* parameter by adding a random uniform shift in the interval [−0.02, 0.02]. If the parameter falls outside the bounds in the prior distribution it is reflected symmetrically back within the bounds. We then have to decide whether to accept or reject the candidate set of parameters. If (Θ_cand_) *<* (Θ_curr_) then we accept the parameters as they share a greater similarity with the experimental data and set Θ_curr_ = Θ_cand_. Otherwise we only accept the candidate parameters with probability exp (((Θ_cand_) (Θ_curr_)) /*ξ*), where *ξ* is a parameter that controls how likely accepting a set of parameters with a higher distance function is. The value of *ξ* is reduced as the algorithm gets closer to a set of parameters that minimises the distance function. Initially *ξ* = 10 but is subsequently reduced to *{*1, 0.1, 0.01, 0.005, 0.001*}* when the distance function of the candidate set of parameters first reaches 50, 30, 20, 18, 17.5 for the plate data and 100, 75, 50, 40, 35 for the cell data. The algorithm continues until it reaches a final set of parameters that has a distance less than 11.08 or 39.2 for the plate and cell data, respectively. For both the plate and cell data we performed this algorithm 1000 times to capture the distribution of parameter values that fit the experimental data.

The ABC-SMC algorithm described above was implemented with custom C++ code (Apple LLVM version 7.0.0, clang-700.1.76). The distributions of the parameters are presented in Fig. S8.

### 15 Kinetic proofreading: binary heatmaps of discrimination and sensitivity

We defined measures of sensitivity and discrimination in order to test whether the KP mechanism can explain both for different KP model parameters. Recall that *λ* is the minimum threshold concentration of productively signalling TCR/pMHC complexes in the *N*-th step. To determine TCR sensitivity, we require that the number of productively signalling TCRs is above the threshold for a single agonist pMHC with the highest affinity *K_D_*_;1_ = *k*_off;1_/*k*_on_. From Eq. (3) we can make the approximation *C*_total_ *≈* min (*L*_0_*, R*_0_) when *L*_0_ + *R*_0_ ≫*K_D_*_;1_. Then, noting that min (1*, R*_0_) = 1 and using Eq. (2) we can write the sensitivity requirement as the following inequality:

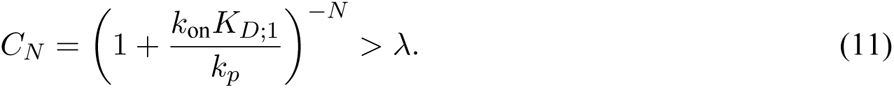

To determine TCR discrimination, we determined whether the number of productively signalling TCRs was below the same threshold *λ* for a pMHC that was expressed at 10,000-fold higher concentration but bound with a Δ-fold lower affinity. With our empirical equation for the discrimination power 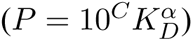 we can calculate the potency *P* for a given ligand affinity. Assuming K_D_ is proportional to *k*_off_ and *P* is a ligand concentration needed to activate the TCR *L*_0_, we can rewrite the equation as 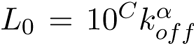. The difference in potency between the ligand interaction with the higher affinity *K_D_*_;1_ and a ligand with lower affinity *K_D_*_;2_ is hence:

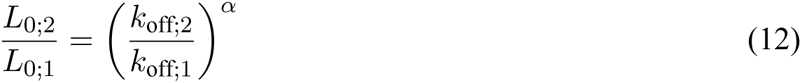

As we require *L*_0;1_ to be 1 to fulfil the sensitivity constrain the equation simplifies to *L*_0;2_ = Δ*^α^* with Δ being the difference in affinity between the two ligands. Hence a ligand with Δ-fold lower affinity than the higher affinity ligand will need a concentration of *L*_0;2_ ligands for activation. For the discrimination constraint we require that a ligand with Δ-fold lower affinity than the highest affinity ligand, needs *L*_0;2_ or more ligands to overcome the threshold of activation. The discrimination requirement can be written as the following inequality:

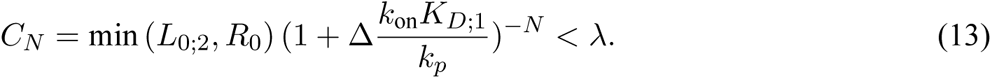

Both of these constraints must be fulfilled simultaneously for a given set of KP parameters in order for the kinetic proofreading model to explain both sensitivity and discrimination.

For the simulation of the KP model (Fig 5G–I) we choose Δ such that *L*_0;2_ = 10, 000 according to Δ*_A_* = 10000^1/^*^α^*. Given that the number of TCRs is *R*_0_ ~ 30, 000, choosing Δ*_L_ < R*_0_ means that the receptors are not saturated with ligands and potency varies linearly with affinity. The final discrimination constraint function is as follows:

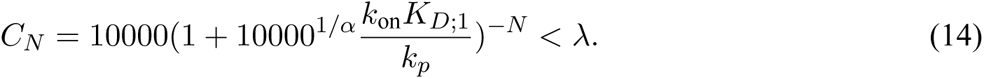

In addition to using the determinstic KP model, we also calculated these sensitivity and discrimination measures using discrete stochastic simulations. We varied *N* and *τ* = 1/*k*_p_. For each pair of parameters (*N, τ*) we simulate 250 realisations of the kinetic proofreading model using a standard Gillespie algorithm until a termination time of *t* = 100 s, which is sufficient in order for the model to have reached a steady state. From this ensemble, an average number of receptors in the final (*N*-th) proofreading step, *C_N_*, is calculated. This ensemble average is compared to the threshold for activation *λ* = 0.1.

Testing for both sensitivity and discrimination for each parameter pair (*N, τ*) requires simulating the model in two different scenarios. The first scenario is with a single ligand and unit dissociation rate, i.e. *k*_off_ = 1. If the ensemble average *C_N_ >* 0.1 then the parameter pair (*N, τ*) observes sensitivity and is shown as a red asterisk in the panels in Fig. 5G–I. For discrimination we increase the number of ligands to Δ*_L_* = 10000 and decreased the affinity of the ligand by Δ*_A_* = 10000^1/^*^α^*, i.e. *k*_off_ = 10000^1/^*^α^*. If the average number of receptors *C_N_ <* 0.1 then discrimination is observed, and the parameter pair (*N, τ*) is shown as a blue square in Fig. 5G–I. Parameter pairs that are shown with both a red asterisk and a blue square observe both sensitivity and discrimination. All stochastic simulations were performed with custom Julia code using the package *DifferentialEquations.jl*.

### 16 Analysis of ***α*** for TCRs from published studies

A supplementary table provides information on each calculation of *α* (Table S5) and specific details on the source of data underlying each calculation (see Supplementary Text).

The broad method was to obtain a measure of ligand potency from each study. If provided by the study, this was often an EC_50_, which is the concentration of ligand eliciting 50% of the maximum response. If not explicitly provided, we estimated ligand potency as *P_X_*, which was defined by the concentration of ligand that produced *X* response. To do this, we drew a horizontal line at *X* on a provided dose-response graph and estimated the ligand concentration where the data intercepted the horizontal line. The disadvantage with this method is that ligand potency was estimated based on the single representative graph provided in the study.

Each study often contained or cited a study that contained estimates of K_D_ or *k*_off_ for the specific TCR/pMHC interactions used in the study. We only included studies where monomeric SPR binding data was available to avoid multimeric binding parameters (e.g. when using tetramers). However, when analysing discrimination by other non-TCR receptors, we included binding data from various methods (e.g. SPR, radio labelled ligands) provided they were monomeric measurements. The use of SPR is important for weak interactions, such as TCR/pMHC, but various methods are available for higher affinity interactions.

The plot of potency over K_D_ or *k*_off_ was fit using linear regression on log-transformed axes. We reported the slope of the fit (i.e. the discrimination power, *α*), the goodness-of-fit measure (*R*^2^), and the P-value for the null hypothesis that the slope is zero (i.e. *α* = 0). We defined significance using the threshold of p=0.05. We found that the calculated *α* was robust to the precise definition of ligand potency so that the same slope was produced when using a different response threshold (e.g. 0.25 or 0.75 instead of the commonly used value of 0.5, not shown).

A subset of the data relied on engineered high affinity TCR/pMHC interactions. It has been observed that increasing the affinity beyond a threshold does not improve ligand potency (39, 59). To avoid underestimating the discrimination power, we found that globally removing data where K_D_ *<* 1 *µ*M avoided entering this saturation regime (with a single exception, see ID 58-61 in Supplementary Information and Fig S6). Similarly, to avoid over-estimating *α*, we did not include data where the potency was extrapolated (i.e. when EC_50_ values were larger than the highest ligand concentration tested). Some studies provided multiple measures of T cell responses and in this case, we produced potency plots for each response and hence were able to obtain multiple estimates of *α*.

We only included discrimination powers in final comparisons (Fig. 3I-O, Fig. 6F) that were statistically significant (*p <* 0.05) with the exception of the original and revised mouse TCR data (Fig. 3I) because only few data were available. We found more studies that performed functional experiments on the original mouse TCRs compared to those that measured binding and therefore, to avoid introducing a potential bias in the analysis, we included only a single calculated *α* for each independent SPR measurement. In the case of the original mouse TCR data, we included 4 calculations of *α* (Table S5, ID 1, 2, 11, 14) and in the case of the revised mouse TCR data, we included 6 calculations of *α* (Table S5, ID 5, 13, 15, 17, 18, 19). We also note that discrimination powers obtained using artificial conditions, when antigen was presented on plates as recombinant protein or when presented on APCs but co-receptors were blocked, were *not* included in aggregated analyses (Fig. 3I-O, Fig. 6F).

### 17 Analysis of ***α*** for other surface receptors from published studies

A supplementary table provides information on each calculation of *α* (Table S6) and specific details on the source of data underlying each calculation (see Supplementary Text).

The general method was similar to that used for the TCR (see previous section). We provide specific information on the analysis of each receptor family below.

Cytokine receptors transduce signals by ligand-induced dimerisation of receptor subunits. We identified 5 studies that produced ligands with mutations that modified binding to either one or both receptor subunits and either reported potency or provided dose-response curves from which potency can be extracted (76–80). As an example, Moraga et al (77) generated IL-13 variants with mutations that resulted in a broad range of affinities to the IL-13R*α*1 subunit but maintained the wild-type interface, and hence the same affinity, to the IL-4R*α* subunit. By measuring cellular responses, such as upregulation of CD86 on monocytes, doseresponse curves were generated for each IL-13 variant allowing us to determine ligand potency. We observed a significant correlation between potency and K_D_ (Fig. 6A). We repeated the analysis for each study (Table S6 ID 1-13). In studies that included ligands with mutations to both receptor interfaces, we plotted potency over the product of the dissociation constants to each interface since this serves as an estimate of the overall affinity 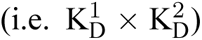 Collating these studies revealed a mean discrimination power of *α* = 0.66 (Fig. 6F).

Like cytokine receptors, RTKs transduce signals by ligand-induced dimerisation. We identified two potential studies to include in the analysis (81, 82). Ho et al (81) generated stem cell factor (SCF) ligand variants to the RTK c-Kit. SCF induces c-Kit dimersation by binding to c-Kit with one interface and binding to another SCF with a different interface generating SCF/c-Kit homodimers. Four SCF variants were used in detailed dose-response assays measuring phosphorylation of ERK (Fig. 6B, left) and AKT (not shown here). Given that the SCF variants included mutations impacting both c-Kit binding and SCF homodimersation, we plotted potency over the product of the dissociation constants for each interface finding a significant correlation for ERK (Fig. 6B, right) and AKT (Fig. S9 ID 16) with discrimination powers of 0.83 and 0.88, respectively. A significant correlation was not observed for the second study using EGFR (Table S6 ID 14) and therefore, we estimated the mean for RTK based on the c-Kit data to be *α* = 0.86 (Fig. 6F).

Although multiple ligands for a given GPCR have been described, they often bind at different GPCR sites to stabilise different receptor conformations and hence transduce qualitatively different signals. Therefore, ligand affinity may not correlate to functional potency. Instead, we focused on identifying studies that used ligands that were confirmed to bind to the same interface with different affinities. As an example, Sykes et al (83) used 7 agonists to the muscarinic M_3_ receptor and confirmed they all bound to the same interface using a binding competition assay. Using titrations of each ligand, they examined the binding of GTP*γ*S to CHO-M_3_ membrane as a measure of response (Fig. 6C, left). Plotting ligand agonist potency over K_D_ produced a significant correlation with a discrimination power of *α* = 0.55 (Fig. 6C, right). We found a similar discrimination power when using a different measure of response (Ca^2+^ mobilisation from CHO-M_3_ cells) from the same study and moreover, similar discrimination powers in other studies investigating the A2A receptor (84) and the chemokine receptors CXCR4 (85) and CXCR3 (86) (Table S6 ID 17-24). Collating these studies revealed a mean discrimination power of *α* = 0.76 (Fig. 6F).

Chimeric antigen receptors (CARs) are therapeutic receptors often expressed in T cells that fuse an extracellular antigen recognition domain to an intracellular signalling domain (often the *ζ*-chain from the TCR). Chmielewski et al (87) generated a panel of CARs that bind the ErbB2 receptor (target antigen) with different affinities. CAR-T cells were stimulated with a titration of recombinant ErbB2 and their ability to produce the cytokine IFN*γ* was used to measure T cell responses (Fig. 6D, left). We found a significant correlation between potency and K_D_ with a discrimination power of *α* = 0.52 (Fig. 6D, right). Similar results were observed using a different ErbB2 CAR (88) and a DNA-based CAR (89) (Table S6 ID 25-28). Together, we found a mean discrimination power of *α* = 0.94 (Fig. 6F).

Lastly, antigen discrimination has also been reported for the B cell receptor (BCR), which shares many structural and functional features with the TCR. Although several studies have investigated BCR ligand discrimination, we identified only a single study with the requisite dose-response curves to quantify discrimination. Batista et al (90) used two lysozyme-specific BCRs (HyHEL10 and D1.3) to perform dose-response curves to wild-type or mutated lysozyme variants measuring the production of the cytokine IL-2 (Fig. 6E, left). We estimated potency directly from the dose-response curves and found a significant correlation with *k*_off_ (Fig. 6E, right). We found the discrimination power for both HyHEL10 and D1.3 BCRs to be *>* 1 (mean of *α* = 1.3, Fig. 6F).

### 18 Statistical analyses

All statistics on discrimination power and sensitivity were performed on log-transformed data, unless stated otherwise. In Fig. 1F, data was compared using a Kruskal-Wallis test with Dunn’s multiple comparison. In Fig. 2K, conditions were compared with a 1-way ANOVA and each condition was compared to *α* = 1 with an independent 1-sample Student’s t test and in Fig. 2L, the 1G4 data was compared with a ordinary 1-way ANOVA and all data was compared using a second ordinary 1-way ANOVA with Sidak’s multi-ple comparison for a pairwise test. In Fig. 3, all comparisons were performed using parametric 1-way ANOVA and/or multiple t-tests (with the stated correction for multiple comparisons) on log-transformed data. In Fig. 4F, plate data was compared using repeated-measure 1-way ANOVA (Geisser-Greenhouse corrected) with Sidak’s comparison for the indicated pairwise comparison. CD58 and ICAM1 were compared to U87 co-culture data using ordinary 1-way ANOVA. Each condition was compared to *α* = 1 using an independent 1-sample Student’s t test. In Fig. 4G, comparison using repeated-measure 1-way ANOVA (Geisser-Greenhouse corrected). In Fig. 5D, plate data compared using a repeated-measure 1-way ANOVA (Geisser-Greenhouse corrected) and APC data and APC vs. plate data was compared using each an ordinary 1-way ANOVA. In Fig. 6F, a 1-way ANOVA compares other receptors and a t-test compares other receptors to the TCR.

## Supplementary Information

### Supplementary Tables

**Table S1:**
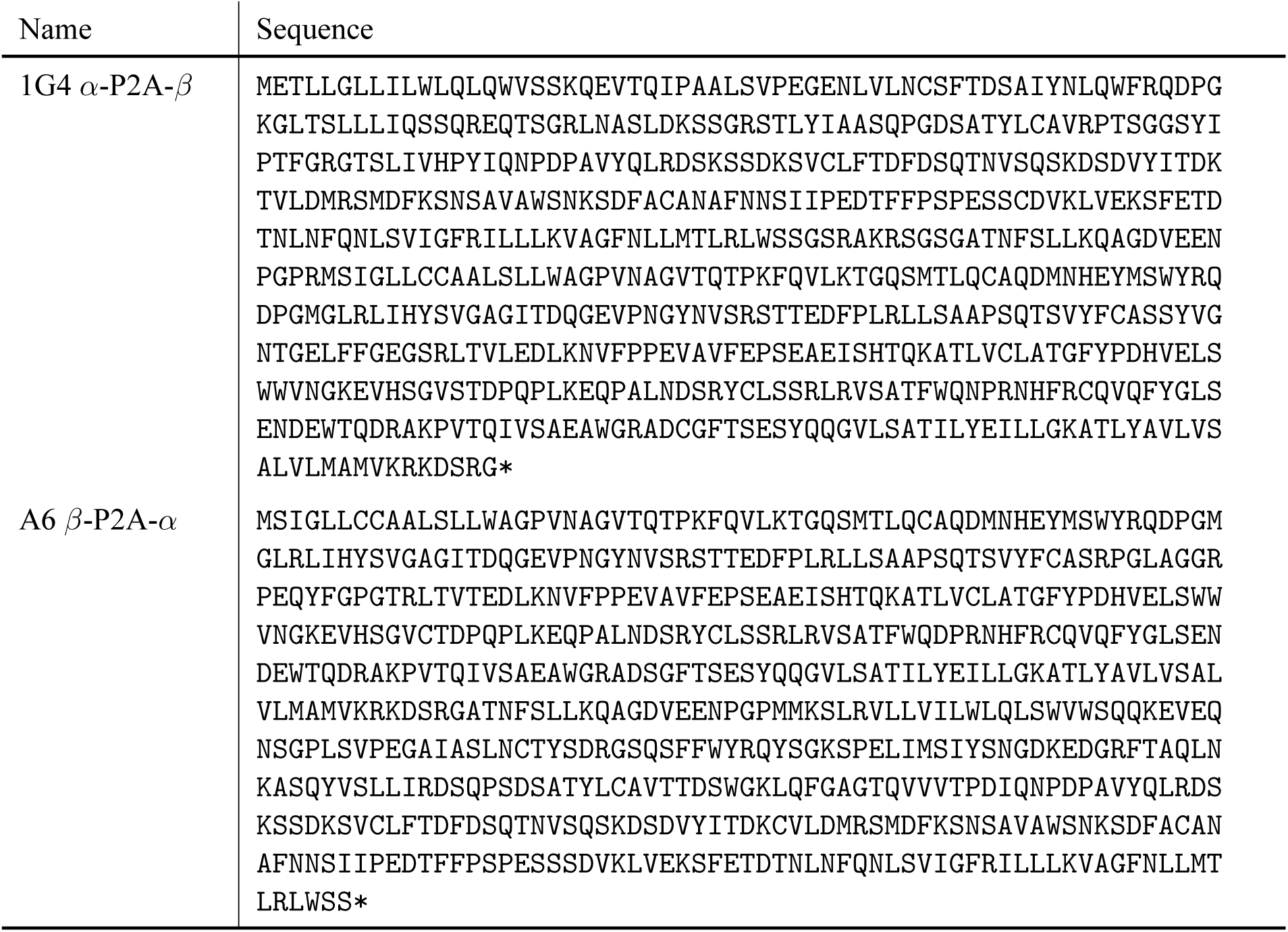
Sequences of 1G4 and A6 TCRs for lentiviral transduction.

**Table S2:** Sequences of soluble A6 TCRs for SPR.

| Name | Sequence |
| --- | --- |
| A6α | MQKEVEQNSGPLSVPEGAIASLNCTYS DRGSQSFFWYRQYSGKSPELIMSIYSNGDKEDG<br>RFTAQLNKASQYVSL LIRDSQPSDSATYLC AVTTDSWGKLQFGAGTQVVVTPDIQNPDA<br>VYQLRDSKSSDKSVCLFTDFDSQTNVSQSKSDVYITDKCVLDMRSMDFKSNSA VAWSNK<br>SDFACANAFNNSIIPEDTFFPSPRESS* |
| A6β-His | MNAGVTQTPKFQVLKTGQSMTLQCAQDMNHEYMSWYRQDPGMGLRLIHYSVGAGITDQGE<br>VPNGYNVSRSTTEDFPLRLLSAAPSQTSVYFCASRPGLAGGRPEQYFGPGTRLTVTEDLK<br>NVFPPEVAVFEPSEAEISHTQKATLVCLATGFYPDHVELSWWVNGKEVHSGVCTDPQLK<br>EQPALNDSRYALSSRLRVSATFWQDPRNHFRCQVQFYGLSENDEWTQDRAKPVTQIVSAE<br>AWGRADHHHHHH* |

**Table S3:** Fitted K_D_ with the indicated method for the 1G4 TCR in SPR at 37*^◦^*C. Includes all peptides used for SPR standard curve and functional experiments.

| Name | Sequence | $K_D$ (Bmax fitted) | | | | $K_D$ (Bmax const.) | | | |
| --- | --- | --- | --- | --- | --- | --- | --- | --- | --- |
| | | Mean (μM) | SD (μM) | CV ( %) | n | Mean (μM) | SD (μM) | CV ( %) | n |
| NYE 9V | SLLMWITQV | 7.97 | 2.55 | 32.0 | 10 | 4.93 | 1.22 | 24.8 | 7 |
| NYE 3A | SLAMWITQV | 7.04 | 1.49 | 21.1 | 5 | 6.28 | 2.16 | 34.3 | 5 |
| NYE 6V | SLLMWVTQV | 17.6 | 5.28 | 30.0 | 6 | 22.8 | 13.8 | 60.8 | 6 |
| NYE 3Y | SLYMWITQV | 36.3 | 10.4 | 28.7 | 5 | 47.3 | 8.58 | 18.1 | 3 |
| NYE 4D | SLLDWITQV | 90.8 | 41.7 | 45.9 | 4 | 140 | 18.3 | 13.0 | 4 |
| NYE 6T | SLLMWTTQV | 84.6 | 30.3 | 35.9 | 4 | 162 | 26.3 | 16.2 | 4 |
| NYE 4A | SLLAWITQV | 157 | 80.8 | 51.4 | 4 | 299 | 48.6 | 16.3 | 4 |
| NYE 5Y | SLLMYITQV | 191 | 107 | 56.3 | 3 | 433 | 103 | 23.8 | 3 |
| NYE 5F | SLLMFITQV | 756 | 1092 | 144.5 | 8 | 1309 | 505 | 38.6 | 8 |

**Table S4:**
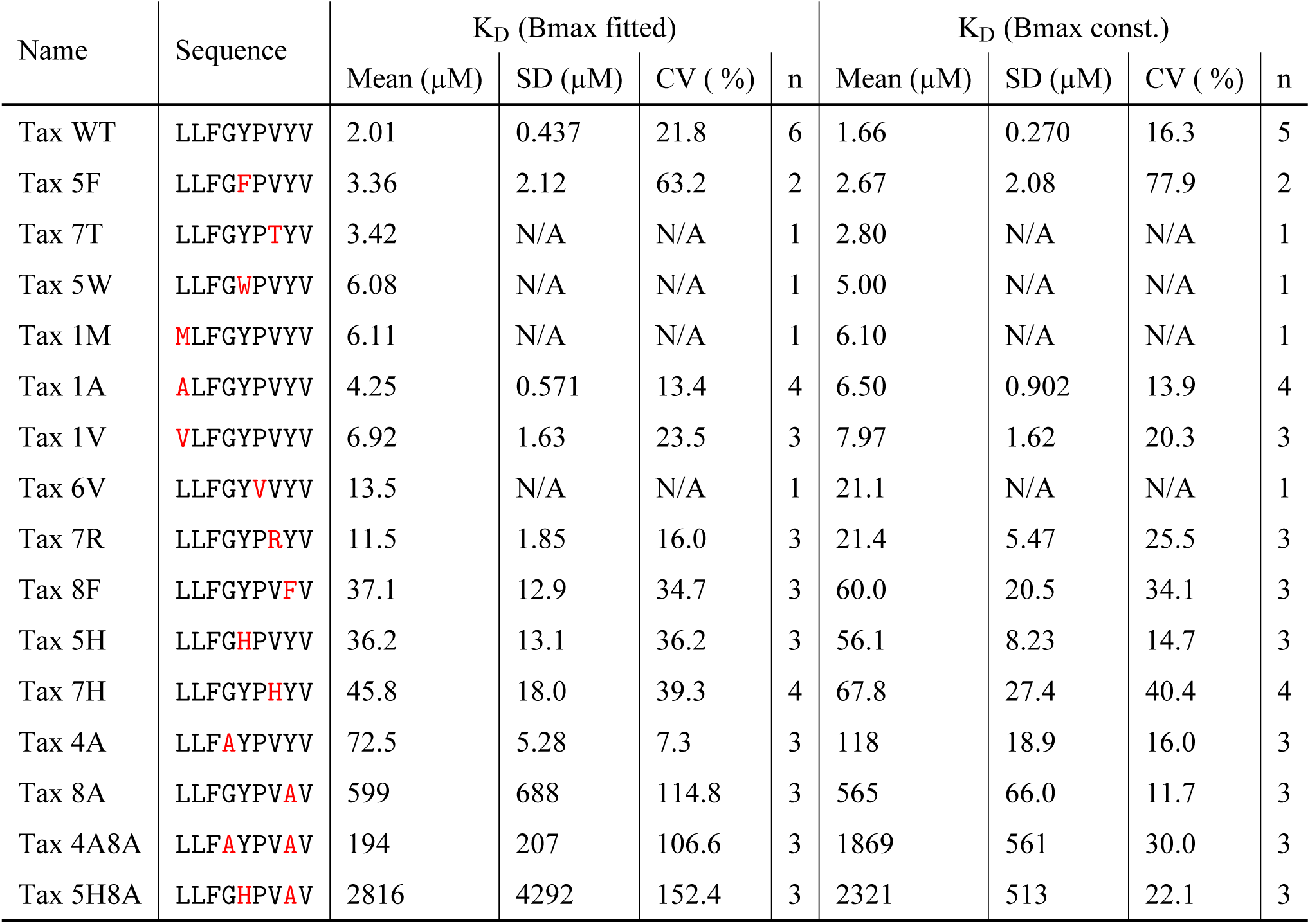
Fitted K_D_ with the indicated method for the A6 TCR in SPR at 37*^◦^*C. Includes all peptides used for SPR standard curve and functional experiments. N/A: not applicable.

**Table S5:**
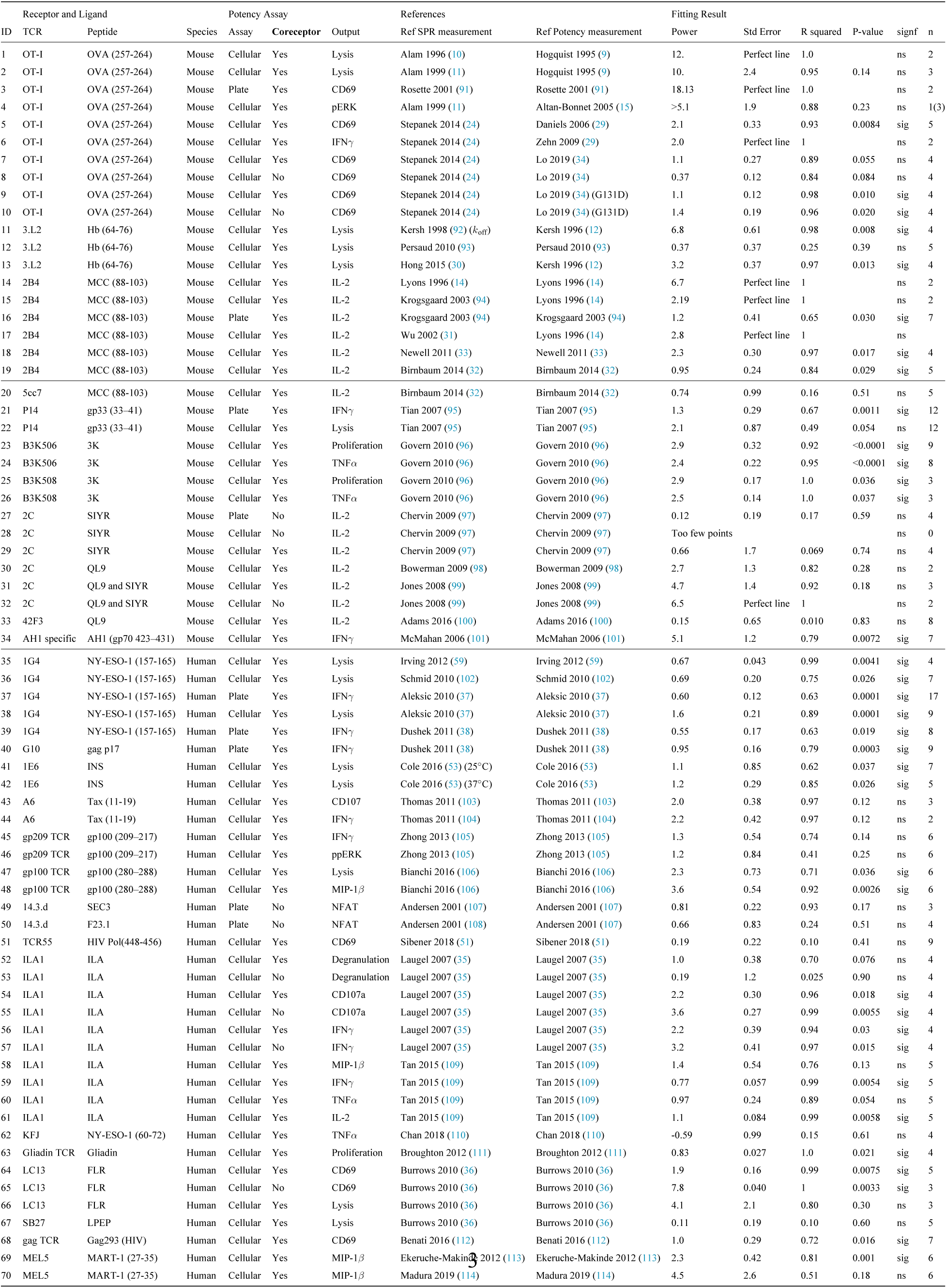
Overview of discrimination powers for TCRs. Each row is associated with an experimental ID that is linked to detailed information on how the data was extracted (see Methods & Supplementary Information text) and to potency plots (Fig. S4-6).

**Table S6:** Overview of discrimination powers for other (non-TCR) surface receptors. Each row is associated with an experimental ID that is linked to detailed information on how the data was extracted (see Methods & Supplementary Information text) and to potency plots (Fig. S9).

|  | Family | Receptor | Output | Ref Affinity measurement | Ref Potency measurement | Power | Std Error | R squared | P value | sig. | n |
| --- | --- | --- | --- | --- | --- | --- | --- | --- | --- | --- | --- |
| 1 | Cytokine | IL-2Rβ | pSTAT5 (NK cells) | Levin 2012 (76) | Levin 2012 (76) | 0.55 | 0.031 | 0.99 | 0.0032 | sig | 4 |
| 2 | Cytokine | IL-2Rβ | %pSTAT5 (T cells) | Levin 2012 (76) | Levin 2012 (76) | 0.74 | 0.047 | 0.99 | 0.0039 | sig | 4 |
| 3 | Cytokine | IL-13Rα1 | pSTAT6 | Moraga 2015 (77) | Moraga 2015 (77) | 0.47 | 0.065 | 0.80 | <0.0001 | sig | 9 |
| 4 | Cytokine | IL-13Rα2 | Proliferation | Moraga 2015 (77) | Moraga 2015 (77) | 0.39 | 0.24 | 0.27 | 0.15 | ns | 9 |
| 5 | Cytokine | IL-13Rα2 | CD86 | Moraga 2015 (77) | Moraga 2015 (77) | 0.44 | 0.085 | 0.90 | 0.0139 | sig | 5 |
| 6 | Cytokine | IL-13Rα1 | CD205 | Moraga 2015 (77) | Moraga 2015 (77) | 0.42 | 0.048 | 0.96 | 0.0031 | sig | 9 |
| 7 | Cytokine | IFNAR1/IFNAR2 | antiviral | Thomas 2011 (78) | Thomas 2011 (78) | 0.71 | 0.11 | 0.79 | <0.0001 | sig | 13 |
| 8 | Cytokine | IFNAR1/IFNAR2 | antiproliferation | Thomas 2011 (78) | Thomas 2011 (78) | 1.3 | 0.11 | 0.93 | <0.0001 | sig | 13 |
| 9 | Cytokine | IFN-γ R1-IL-10Rβ | pSTAT1 | Mendoza 2017 (79) | Mendoza 2017 (79) | 0.024 | 0.046 | 0.030 | 0.61 | ns | 11 |
| 10 | Cytokine | IFN-γ R1-IL-10Rβ | antiviral | Mendoza 2017 (79) | Mendoza 2017 (79) | 0.034 | 0.063 | 0.032 | 0.60 | ns | 11 |
| 11 | Cytokine | IFN-γ R1-IL-10Rβ | antiproliferation | Mendoza 2017 (79) | Mendoza 2017 (79) | 0.50 | 0.14 | 0.60 | 0.0054 | sig | 11 |
| 12 | Cytokine | gp130 (and IL6Rα) | pSTAT1 | Martinez-Fabregas 2019 (80) | Martinez-Fabregas 2019 (80) | 0.54 | 0.24 | 0.84 | 0.26 | ns | 3 |
| 13 | Cytokine | gp130 (and IL6Rα) | pSTAT3 | Martinez-Fabregas 2019 (80) | Martinez-Fabregas 2019 (80) | 0.52 | 0.32 | 0.57 | 0.24 | ns | 4 |
| 14 | RTK | EGFR | growth rate | Reddy 1996 (82) | Reddy 1996 (82) | 0.55 | 0.10 | 0.97 | 0.12 | ns | 3 |
| 15 | RTK | c-Kit | pERK | Ho 2017 (81) | Ho 2017 (81) | 0.83 | 0.14 | 0.95 | 0.028 | sig | 4 |
| 16 | RTK | c-Kit | pAKT | Ho 2017 (81) | Ho 2017 (81) | 0.88 | 0.14 | 0.96 | 0.023 | sig | 4 |
| 17 | GPCR | A2A receptor | whole cell | Guo 2012 (84) | Guo 2012 (84) | 0.29 | 0.20 | 0.22 | 0.19 | ns | 10 |
| 18 | GPCR | A2A receptor | cAMP | Guo 2012 (84) | Guo 2012 (84) | 0.71 | 0.29 | 0.42 | 0.041 | sig | 10 |
| 19 | GPCR | M3 muscarinic receptor | Calcium | Sykes 2009 (83) | Sykes 2009 (83) | 0.77 | 0.53 | 0.29 | 0.21 | ns | 7 |
| 20 | GPCR | M3 muscarinic receptor | GTPγS Binding | Sykes 2009 (83) | Sykes 2009 (83) | 0.55 | 0.18 | 0.65 | 0.029 | sig | 7 |
| 21 | GPCR | CXCR4 | Voltage | Guyon 2013 (85) | Guyon 2013 (85) | 0.57 | 0.40 | 0.40 | 0.25 | ns | 5 |
| 22 | GPCR | CXCR3 | Calcium mobilization | Heise 2005 (86) | Heise 2005 (86) | 0.73 | 0.033 | 1.0 | 0.029 | sig | 3 |
| 23 | GPCR | CXCR3 | GTPγS Binding | Heise 2005 (86) | Heise 2005 (86) | 1.1 | 0.0039 | 1 | 0.0024 | sig | 3 |
| 24 | GPCR | CXCR3 | Migrated Cells | Heise 2005 (86) | Heise 2005 (86) | 0.56 | Perfect line | 1 |  | ns | 2 |
| 25 | CAR | C6.5 (scFc against ErbB2) | INFγ | Chmielewski 2004 (115) | Chmielewski 2004 (115) | 0.52 | 0.028 | 0.99 | 0.0003 | sig | 5 |
| 26 | CAR | ErbB2 | CD107 | Liu 2015 (88) | Liu 2015 (88) | 1.1 | 0.14 | 0.97 | 0.017 | sig | 4 |
| 27 | CAR | ErbB2 | Proliferation | Liu 2015 (88) | Liu 2015 (88) | 0.64 | 0.18 | 0.87 | 0.068 | ns | 4 |
| 28 | CAR | DNA-CAR | pERK | Taylor 2017 (89) | Taylor 2017 (89) | 1.2 | 0.17 | 0.96 | 0.019 | sig | 4 |
| 29 | BCR | D1.3 | IL2 | Batista 1998 (90) | Batista 1998 (90) | 1.4 | 0.18 | 0.94 | 0.0014 | sig |  |
| 30 | BCR | HyHEL10 | IL2 | Batista 1998 (90) | 6 Batista 1998 (90) | 1.3 | 0.12 | 0.98 | 0.0088 | sig | 4 |

### Supplementary Figures

**Figure S1:**
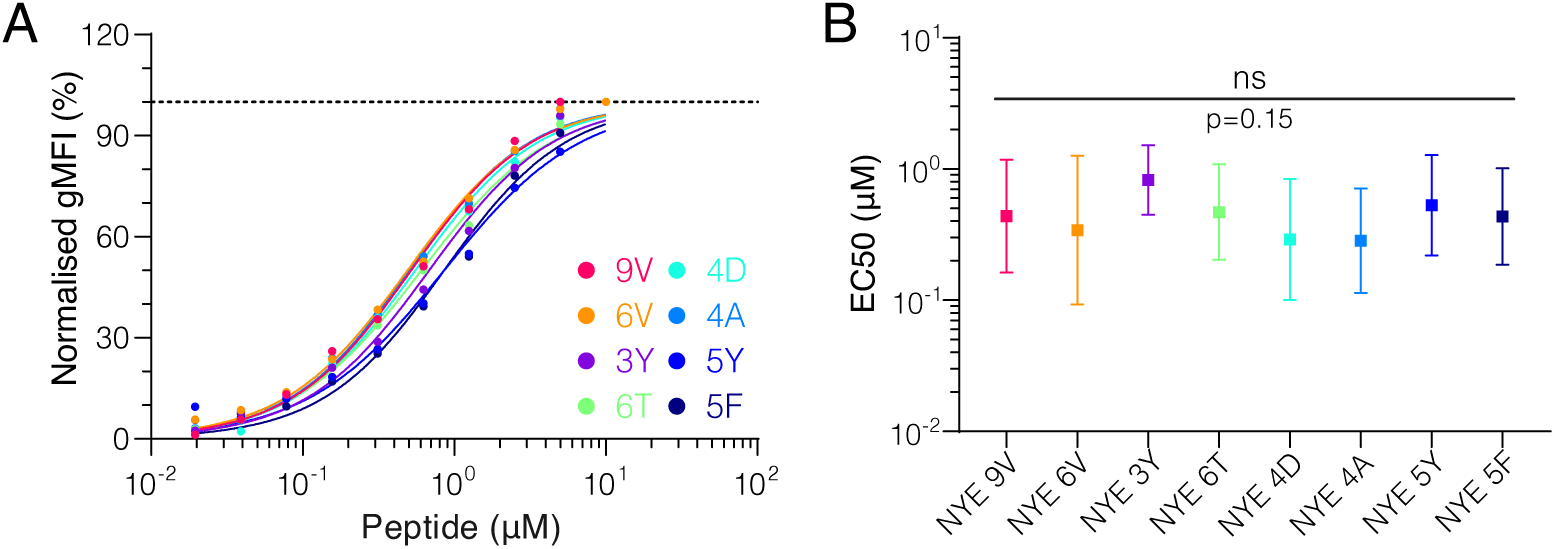
All NYE peptides load similarly on T2 cells. **(A)** Example of upregulation of HLA-A2 on T2 cells over different peptide pulsing concentrations. **(B)** Summary EC_50_s of loading on T2 cells. Shown are means with standard deviation. Pooled log-transformed data from 3 independent experiments were compared using a repeated-measure 1-way ANOVA with Geisser-Greenhouse correction.

**Figure S2:**
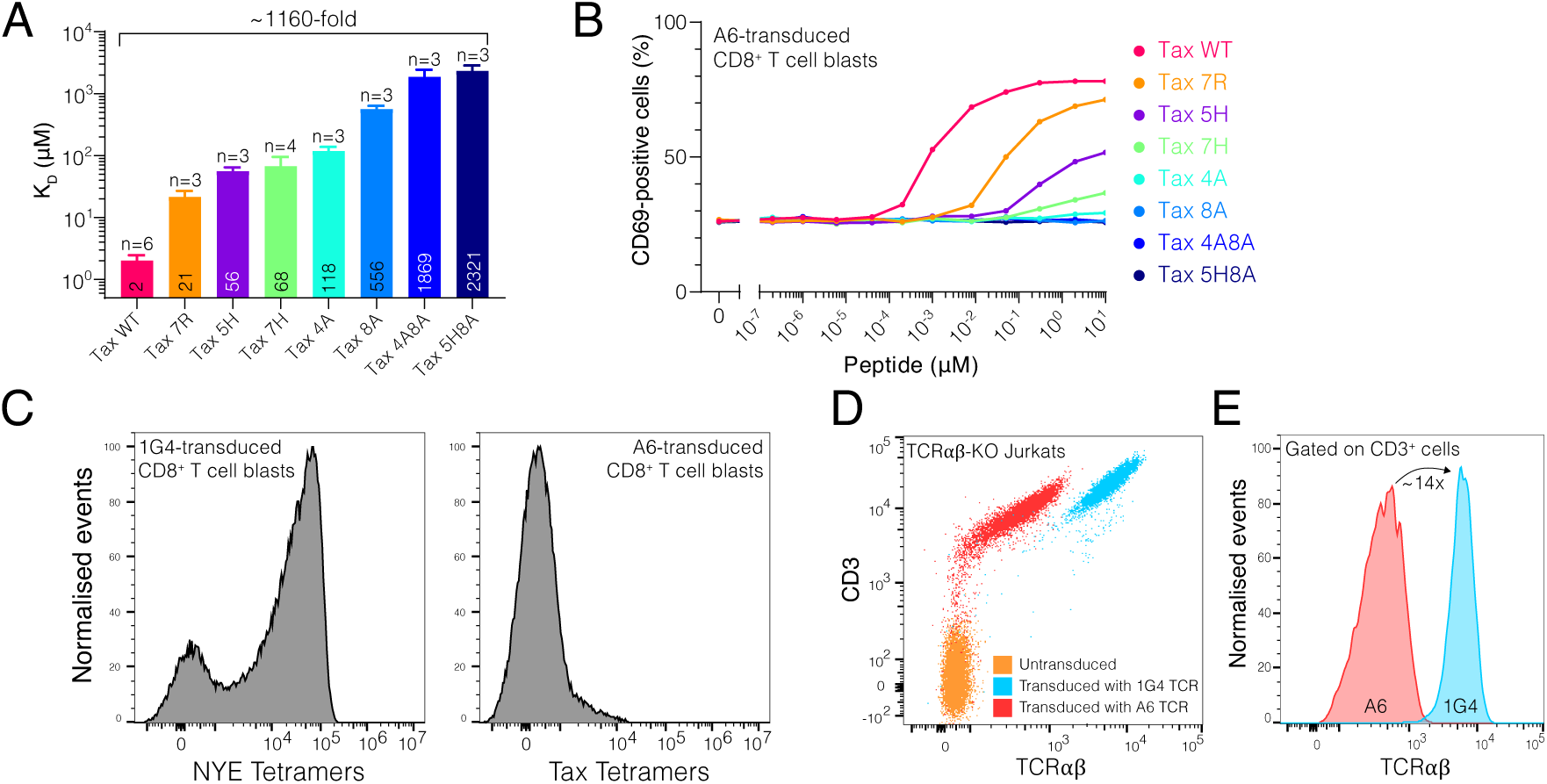
T cells transduced with A6 TCR have low expression and do not respond to ultra-low affinity pMHCs. **(A)** Affinity of the A6 TCR for the panel of Tax-related peptide, as measured by SPR. Shown are means with SD (the mean is also shown as a number wihtin the bar). **(B)** Activation of A6 expressing CD8^+^ T cell blasts by U87 APCs pulsed with the indicated concentration of Tax-related peptides. **(C)** Binding of the indicated pMHC tetramers to A6-versus 1G4-transduced CD8^+^ T cell blasts, as measured by flow cytometry. **(D)** Measurement by flow cytometry of cell surface expression of CD3 and TCR*αβ* by CD8^+^ T cell blasts following transduction with the indicated TCR into TCR*αβ*-KO Jurkats. **(E)** Relative expression of TCR*αβ* using the data in **(E)**, gated on CD3^+^ cells.

**Figure S3:**
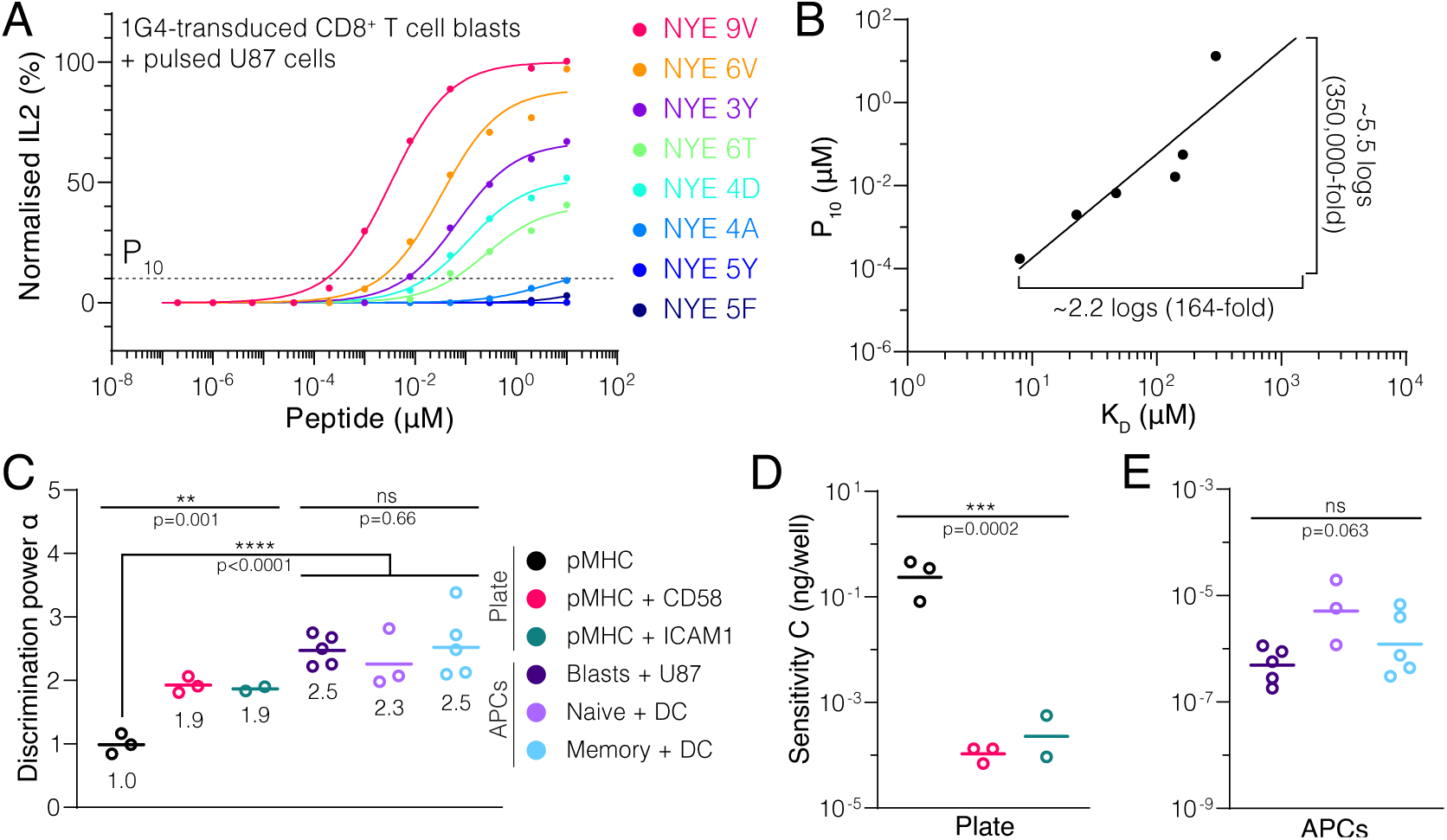
The discriminatory power based on cytokine production. **(A)** Dose-response for secretion of IL2 into the supernatant. Dotted line indicates 10 % activation threshold (*P*_10_). **(B)** A plot of potency obtained from **(A)** over affinity showing the power law fit. Lower affinity antigens that did not induce detectable IL2 (NYE 5Y and 5F) are not included in the plot. **(C)** Geometric mean of discrimination power (*α*). **(D,E)** Geometric mean sensitivity measure (*C*) for stimulation with **(D)** immobilised ligands or **(E)** APC co-culture.

**Figure S4:**
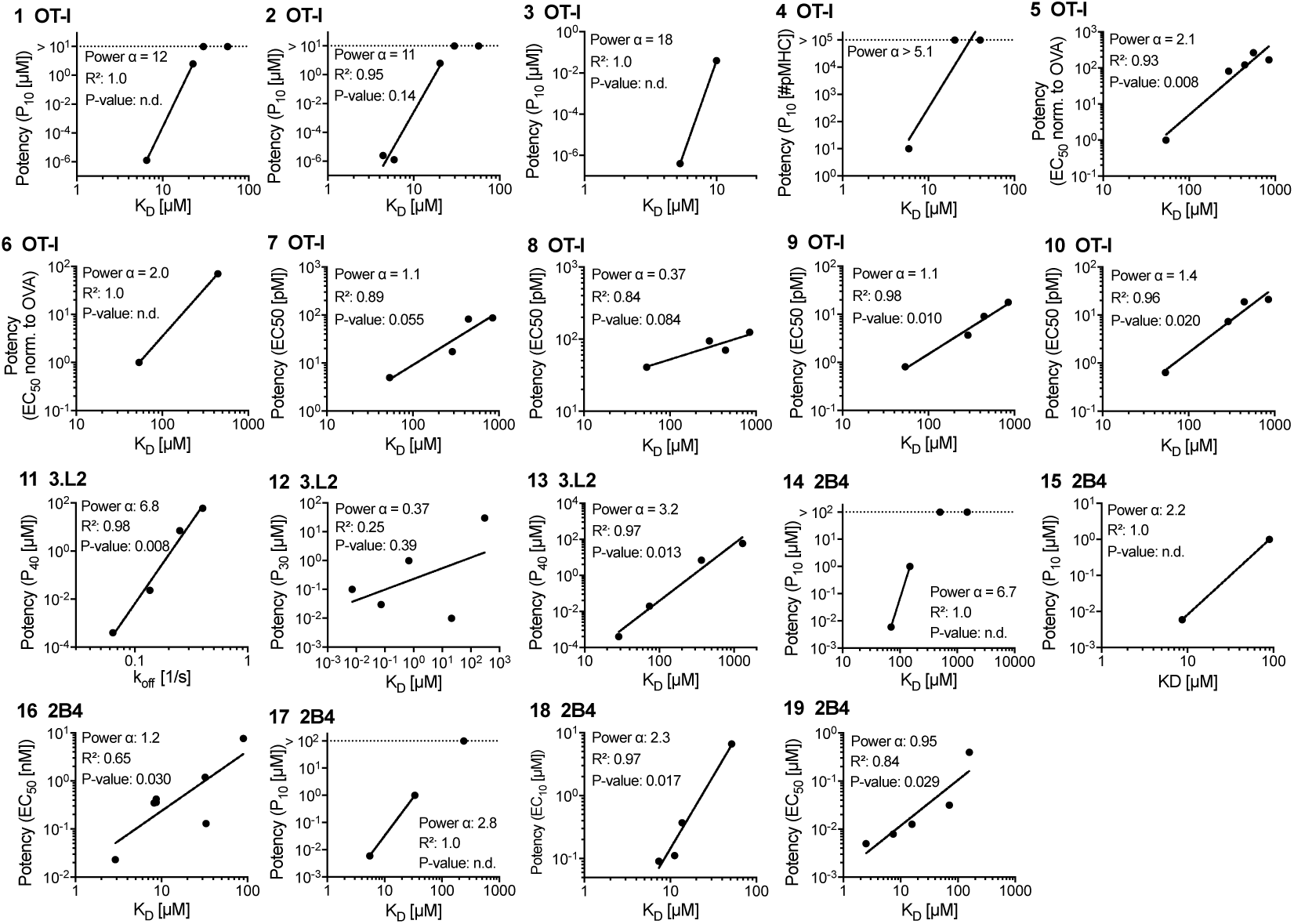
Potency over K_D_ data for the original mouse TCRs (OT-I, 3L.2, and 2B4). Each panel is linked by an ID to an entry in Table S5 and a paragraph in the Supplementary Text.

**Figure S5:**
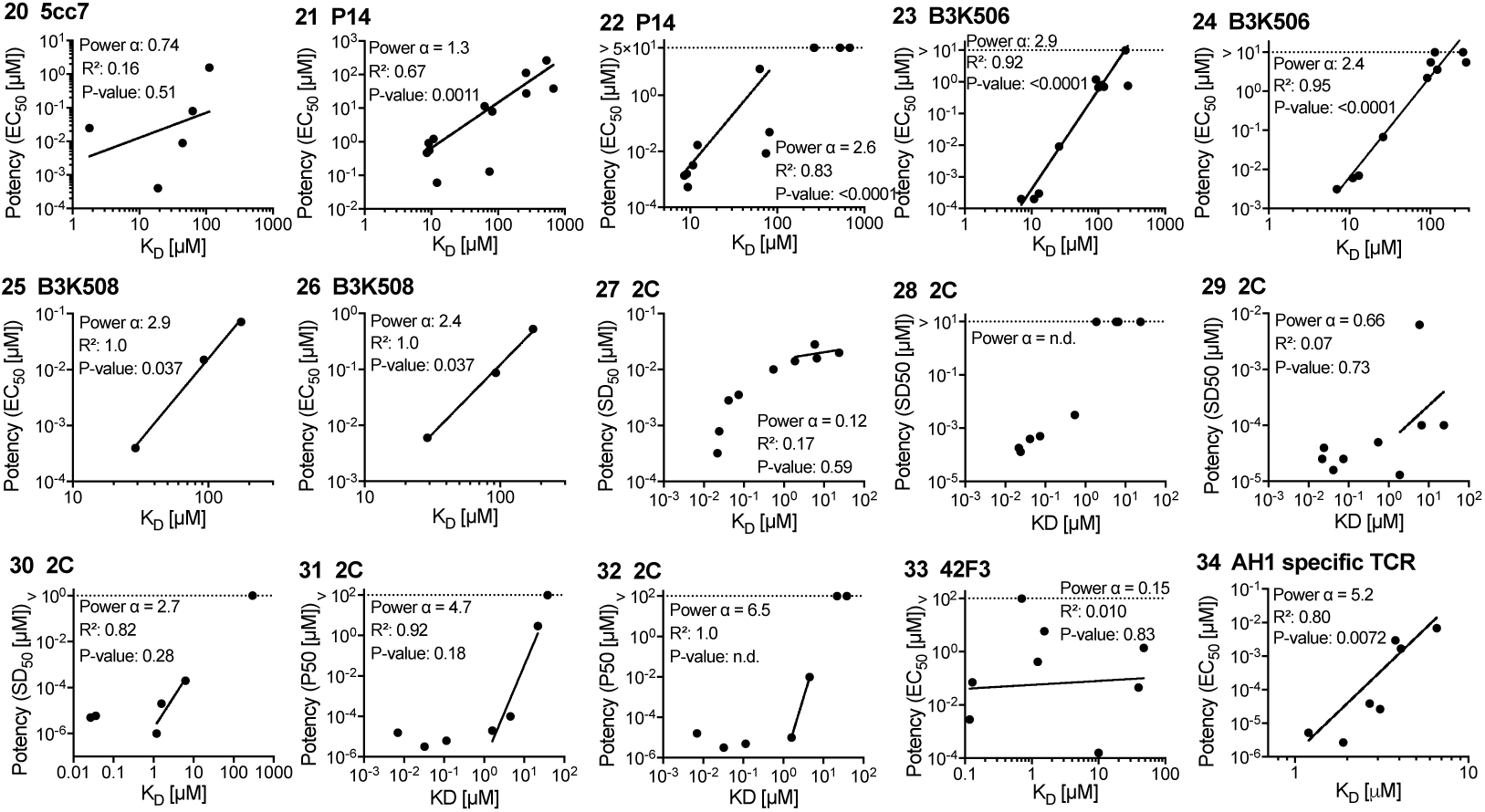
Potency over K_D_ data for other mouse TCRs. Each panel is linked by an ID to an entry in Table S5 and a paragraph in the Supplementary Text.

**Figure S6:**
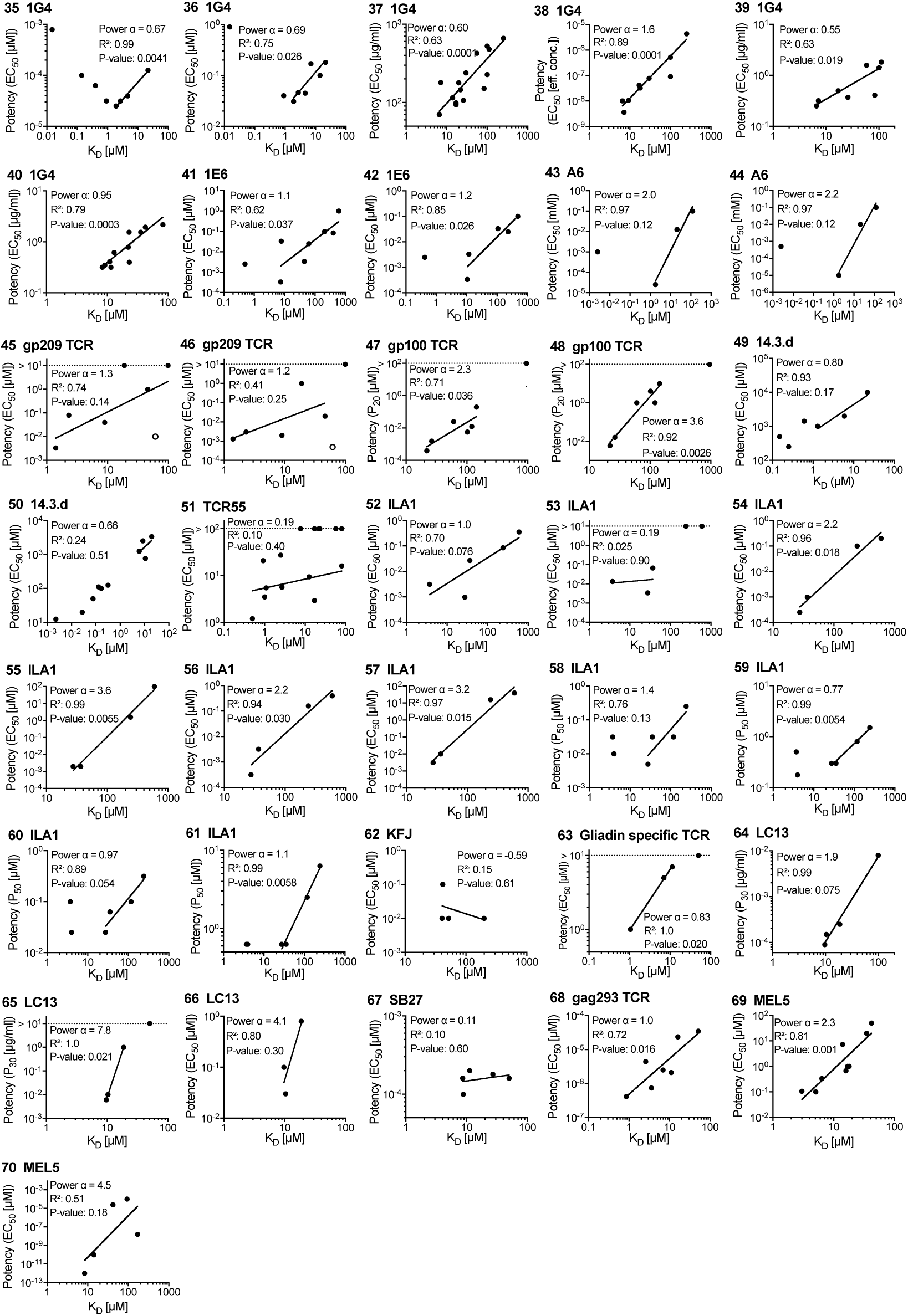
Potency over K_D_ data for other human TCRs. Each panel is linked by an ID to an entry in Table S5 and a paragraph in the Supplementary Text.

**Figure S7:**
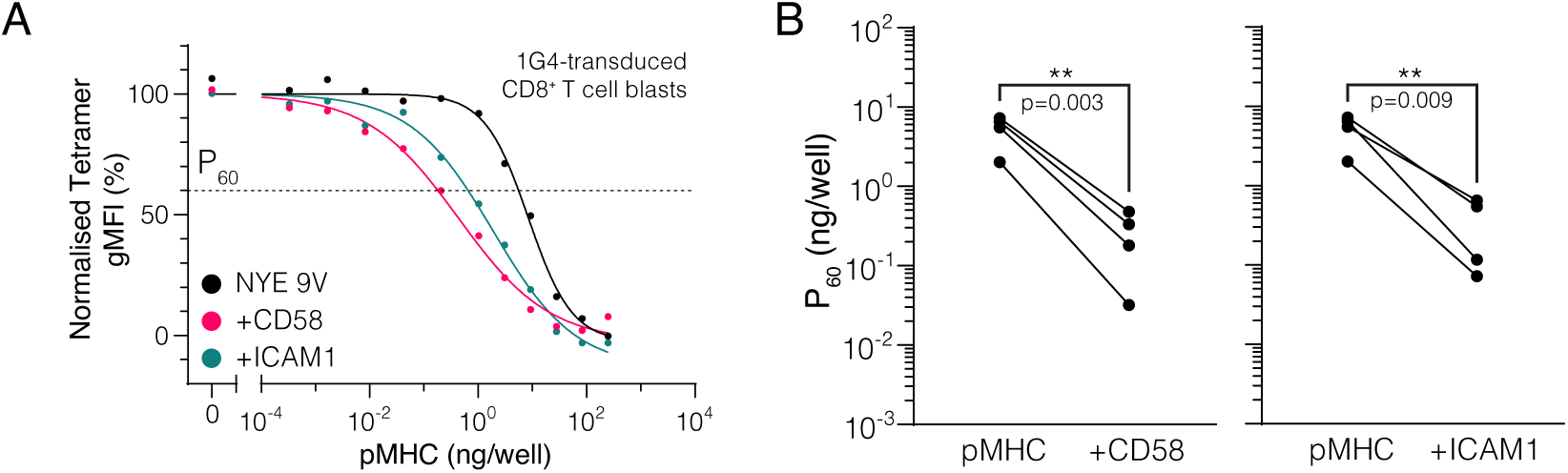
Engagement of CD2 or LFA-1 increases TCR downregulation. **(A)** Example dose-dependent for downregulation of TCR and its acceleration by engagement of CD2 by CD58 or LFA-1 by ICAM1. P_60_ is defined as the dose required to induce 40 % downregulation. Normalised to pMHC alone data. **(B)** Summary effect of CD2 or LFA-1 on TCR downregulation. Each dot represent an individual experiment. Statistics by repeated-measure 1-way ANOVA (with Geisser-Greenhouse correction) of log-transformed data (n=4) with Dunnett’s multiple comparison test.

**Figure S8:**
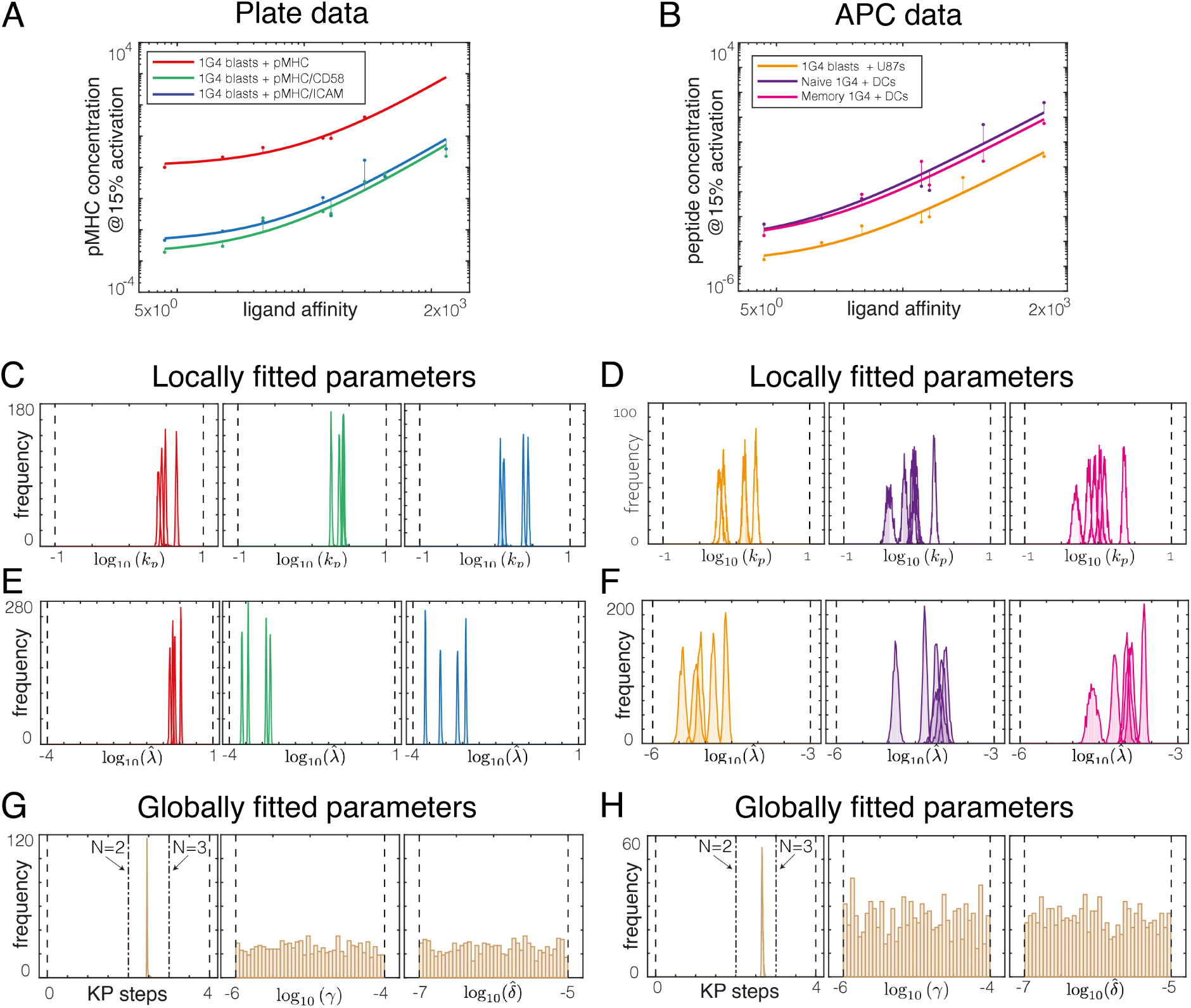
Direct fit of the kinetic proofreading model to potency data using the ABC-SMC method. **(A,B)** Examples of **(A)** plate and **(B)** APC potency data (dots) fitted with the KP model (lines). **(C-H)** The ABC-SMC method provides a distribution of all parameters that are consistent with the high quality fits presented in A and B. **(C-F)** Distribution of locally fitted parameters reveal that the proofreading rate (*k_p_*) and the sensitivity parameter (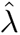) can be uniquely determined for each **(C, E)** plate experiment and **(D, F)** APC experiment. **(G-H)** Distribution of globally fitted parameters reveal that the number of KP steps (N) can be uniquely determined but that two additional parameter (*γ*, 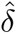) cannot be uniquely determined as they do not exhibit peaked distributions for **(G)** plate experiments and **(H)** APC experiments.

**Figure S9:**
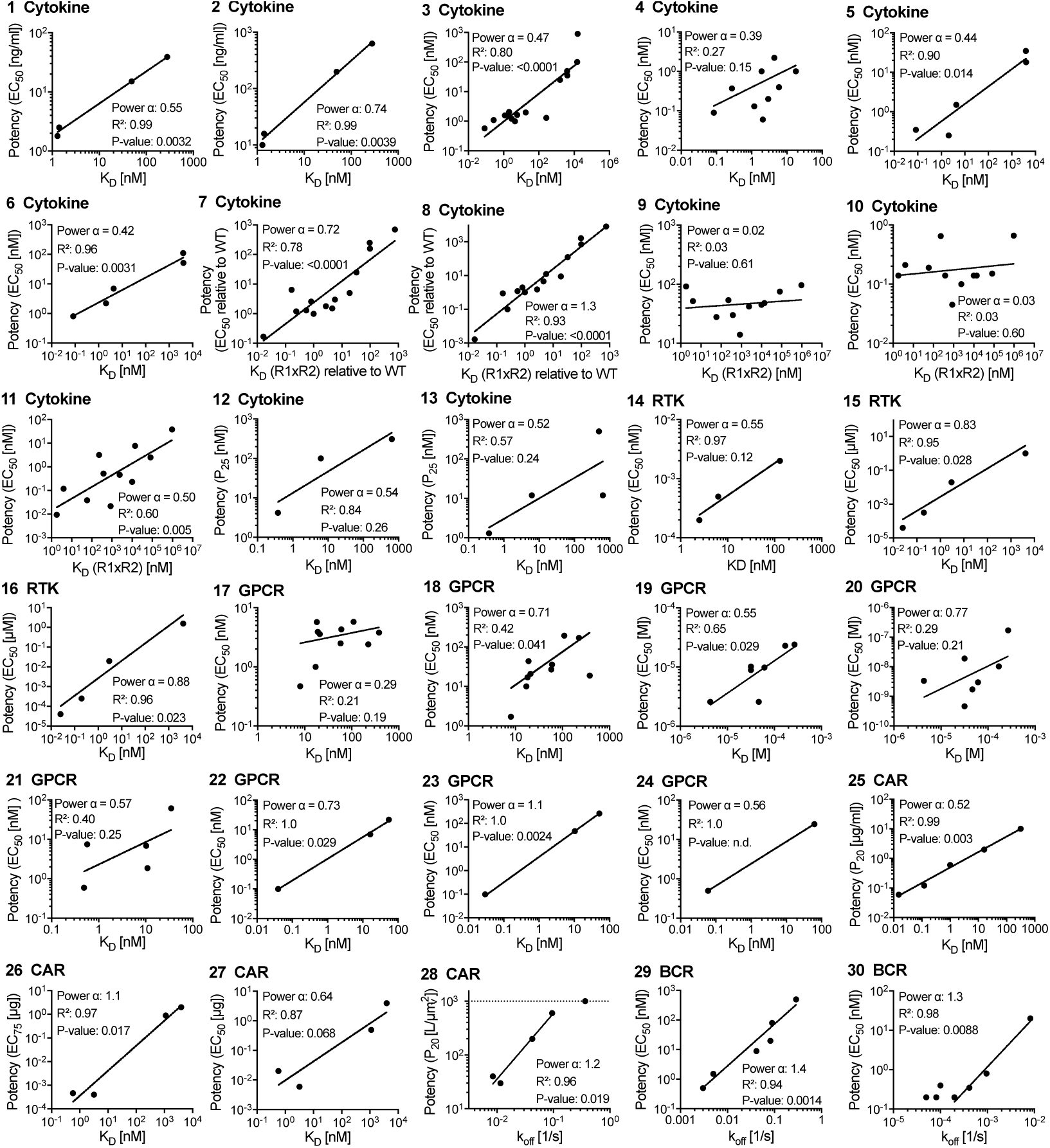
Potency over K_D_ data for other (non-TCR) receptors. Each panel is linked by an ID to an entry in Table S6 and a paragraph in the Supplementary Text.

## Supplementary Text

We provide information on each potency plot we generated in the sections that follow, including the location of the potency and K_D_ data within each study. References to Figures and Tables are to those in the cited manuscript not to those in the present study. We provide the value of *α* produced and additional information, including *R*^2^ and p-values are provided in Table S5 and S6.

### Original mouse TCR data

#### OT-I

##### ID 1 (10)

Experiments were performed with the murine transgenic TCR OT-I that binds to a peptide from ovalbumin (OVA) presented on H2Kb. Affinity was measured by SPR at 25*^◦^*C. Affinity values were taken from Table 1 and Figure 3f. For the power analysis, we used K_D_ values estimated from the binding kinetics (kinetic K_D_ values). Potency measures for the OVA peptide and peptide variants were previously measured by Hogquist et al (9). OT-I T cell responses to OVA and single amino acid peptide variants (A2 and E1) were measured in a cell lysis assay. For the power analysis we extracted the potency of the peptides by reading the *P*_10_ (peptide concentration producing 10% specific lysis) from dose response curve in Figure 2. We excluded peptides that did not result in any response. We were able to include two data points with potency and affinity values for the power analysis producing *α* = 12 (ID 1).

##### ID 2 (11)

The OT-I TCR binding to the peptides derived from OVA were used with affinity and kinetics measured by SPR at 6, 25 and 37*^◦^*C. Unusual biphasic binding was observed at 37*^◦^*C for some peptides with two *k*_on_ and two *k*_off_ values reported based on a slow first and fast second step binding. Affinity values were provided in Table 1. To avoid picking the fast or slow phase parameters, we used the monophasic affinity data measured at 25*^◦^*C for the power analysis. Potency data was taken from Hogquist et al (9). Three data points were included in the analysis producing *α* = 10 (ID 2).

##### ID 3 (91)

OT-I TCR affinity and functional activity was measured when binding its wild type ligand OVA or single amino acid variants (G4). Affinity values of TCR-pMHC interaction, measured by SPR at 25 and 37*^◦^*C, were provided in Table 1. Similar to Alam et al (11), TCR binding to MHC loaded with OVA showed biphasic binding at 37*^◦^*C. As before, we used the data measured at 25 *^◦^*C for the power analysis. Functional data was generated with T cells isolated from OT-I transgenic mice. T cells were then stimulated with peptide-MHC complexes immobilised on plates. We read off potency data from dose response curves in Figure 1. Only two data points were available for calculating the discrimination power *α* producing *α* = 18 (ID 3)

##### ID 4 (15)

In this study, the OT-I T cell response when stimulated with OVA and was determined by phosphorylation of the kinase ERK in the MAPK pathway. Responses to OVA peptide as well as two peptide variants were studied. Potency values were extracted as *P*_10_ from dose response curves in Figure 1C. Only the OVA peptide could activate T cells above background. Potency for unresponsive peptides was set to the highest concentration used in assay. Therefore, the discrimination power *α* calculated with these data points gives a lower bound on the actual value for *α*. Using the affinity data from Alam et al (11) produced *α >* 5.1 (ID 4).

#### 3.L2

##### ID 11 (92)

This paper contains affinity and kinetic data for 3.L2 TCR which recognises murine haemoglobin (Hb 64-76) measured by SPR at 25 *^◦^*C. We used *k*_off_ values for the power analysis as K_D_ values did not correlate (see main text). The D73 peptide was excluded from power analysis because this mutation impacted peptide loading to MHC. Potency data for this TCR was taken from Kersh et al (12). In the functional experiments, 3.L2 T cell hybridoma cells were incubated with antigen presenting cells pulsed with peptides. Activation was measured by lysis of target cells. *P*_40_ values (ligand concentration at 40% lysis) of T cell response were given in Figure 4, the corresponding dose response curve was shown in Figure 5. Four data points were included in the analysis and produced *α* = 6.8 (ID 11).

#### 2B4

##### ID 14 (14)

The 2B4 TCR used in this study recognises a moth cytochrome c (MCC) peptide bound to MHC class II molecule I-Ek. Table 2 provides K_D_ values using SPR at 25*^◦^*C. The potency of the peptides was determined with T cell hybridomas, stimulated by peptide-pulsed APCs, with activation determined by IL-2 production. For the power analysis we extracted the *P*_10_ from the dose response curve in Figure 1A. Two data points were available for the power analysis producing an *α* = 6.7 (ID 14).

### Revised data for the original mouse TCRs

#### OT-I

##### ID 5 (24, 29)

Revised affinity data for OT-I TCR was published by Stepanek et al (24). The K_D_ values were taken from the Table in Figure S1D. Potency data for the same set of peptide variants was measured by Daniels et al (29). Functional experiments were done with pre-selection OT-I double-positive thymocytes. T cell activation was measured by expression of CD69 after incubation with peptide-pulsed antigen presenting cells. *EC*_50_ values, corrected for small differences in peptide affinity for MHC and normalised to OVA, were given in Figure 1a. Together, these papers provide 5 data points producing *α* = 2.1 (ID 5).

##### ID 6 (24, 116)

Zehn et al (116) provided additional functional data for OT-I TCR. Potency data is measured by intracellular IFN*γ* production by OT-I T cells stimulated with peptide pulsed antigen presenting cell. The *EC*_50_ values, given in table in Supplementary Figure 2C, were normalised to OVA. To calculate the discrimination power, we used K_D_ values from Stepanek et al (24). The two data points available produced a power of *α* = 2.0 (ID 6).

##### ID 7 - 10 (24, 34)

Lo et al (34) generated additional functional data for the OT-I TCR. Functional response of CD8^+^ or CD8*^−^* Jurkat cells expressing the OT-I TCR after stimulation with peptide pulsed antigen presenting cells was measured by CD69 upregulation. The *EC*_50_ values were provided in Supplementary Figure 7C. K_D_ values were previously measured by Stepanek et al (24). The study included affinity and potency data for when one of the phosphorylation sites of LAT was mutated. The calculated discrimination power was the same (*α* = 1.1 for both wild-type LAT (ID 7) and mutated LAT (ID 9) unless CD8 was not present, in which case *α*= 0.37 (ID 8) or *α* = 1.4 (ID 10) using Jurkats expressing wild-type or mutated LAT, respectively.

#### 3.L2

##### ID 12 (93)

In this study, the 3.L2 TCR as well as the M15 TCR, a high-affinity TCR engineered from the 3.L2 TCR system, were used, both TCRs bind to murine hemoglobin (Hb 64-76). Table 1 provides K_D_ values using SPR with at 25*^◦^*C. Functional data was generated by incubating T hybridoma cells with peptide-pulsed APCs and measuring IL-2 production. We extracted potency values from dose-response curves in Figure 1b and c. Potency values from both TCR systems produces *α* = 0.37 (ID 12).

##### ID 13 (30)

This paper contain binding and potency data for the 3.L2 TCR interacting with the WT haemoglobin peptide and a panel of altered peptide ligands. Table 2 provides K_D_ values using SPR at 25*^◦^*C. The paper does not contain new potency measurement and therefore, we used potency values measured by Kersh and Allen (12) for the power analysis. This dataset produces *α* = 3.2 (ID 13).

#### 2B4

##### ID 17 (31)

This paper contains affinity data for 2B4 TCR binding to its cognate MCC antigen and a set of variant peptides. Table 1 contains K_D_ values determined by SPR at 25*^◦^*C. To compare with the original discrimination power, we used the original potency data (14) to produce *α* = 2.8 (ID 17).

##### ID 15 - 16 (94)

Revised data for 2B4 TCR is provided by Krogsgaard (94). Table 1 provides K_D_ values measured by SPR at 25*^◦^*C. For potency measurement, T cells from transgenic 2B4 mice were incubated peptide MHC molecules immobilised on plates, activation was measured by IL-2 production (*EC*_50_ values were given in Table 1). All ligands were included in the analysis, including those initially labelled as outliers in the publication. The resulting *α* is 1.2 (ID 16). We also calculated *α* with affinity data from this study and potency data from Lyons et al (14) (*α* = 2.2, ID 15).

##### ID 18 (33)

Newell et al (33) studied the 2B4 and the 226 TCRs that bind to MCC. The K_D_ values were measured by SPR at 25*^◦^*C and provided in Figure 5D (2B4) and 6B (266). T cell hybridomas were incubated with peptide-pulsed cells and T cell activation measured by IL-2 production. *P*_10_ (concentration at 10% maximal IL-2 produced by wild-type 2B4) values given in Figure 5C (2B4) and 6 (266). Data for 2B4 produces *α* = 2.3 (ID 18). The 266 TCR was not included in the analysis, because not enough data points were available.

##### ID 19- 20 (117)

The affinity and potency of the 2B4 and the related 5cc7 TCR, which both interact with MCC, were reported. As before, SPR was used to report K_D_ values. Functional assays were done with blasted transgenic T cell incubated with peptide pulsed cells. To determine potency, IL-2 production was measured. We extracted both K_D_ values and *EC*_50_ values from Figure 4C. Data for 2B4 produced *α* = 0.95 (ID 19) and for 5cc7 produced *α* = 0.74 (ID 20).

### Other mouse TCRs

#### P14

##### ID 21 - 22 (95)

The mouse P14 TCR that recognises a set of altered peptides from the lymphocytic choriomeningitis virus epitope gp33–41 on murine class I MHC Db. All binding parameters were measured by SPR at 25*^◦^*C. In functional assays, T cell cytotoxicity, and IFN-*γ* production of blasted splenocytes from P14 TCR transgenic mice was measured when binding peptide-MHC. Cytotoxicity was measured in a cellular assay, IFN*γ* production in a plate assay. The *EC*_50_ is used as potency measurement. All affinity and potency data were provided in Table 2. The *α* value for this TCR system is 2.1 for cytotoxicity assay (ID 21) and 1.3 for IFN*γ* assay (ID 22).

#### B3K506 and B3K508

##### ID 23 - 26 (118)

The MHC-II restricted B3K506 and B3K508 TCRs that recognise the 3K peptide were studied. The K_D_ values were measured by SPR at 25*^◦^*C. T cell response was measured by T cell proliferation and cytokine production after stimulation with peptide-pulsed APCs. All K_D_ and *EC*_50_ values were given in Table S1. The B3K506 system produced *α* = 2.9 (ID 23) and *α* = 2.4 (ID 24) and the B3K508 system produced *α* = 2.9 (ID 25) and *α* = 2.5 (ID 26) for proliferation and TNF*α* production, respectively.

#### 2C

##### ID 27 - 29 (119)

A panel of TCRs, derived from the murine 2C TCR, that differed in their affinity to the SIYR peptide presented on H-2Kb were used. The K_D_ values were measured by SPR at 25*^◦^*C and provided in Table 1. Functional experiments were done with T cell hybridomas with or without CD8 expression. T cells were either incubated with peptides immobilised on plates or with antigen presenting cells pulsed with peptides. For cellular experiments *EC*_50_ values are given in Figure 3B and D (with and without CD8 respectively), for plate assays in Figure 4B (only CD8 negative data). Most of the ligands have a K_D_ *<* 1 *µ*M, hence the data points were excluded from the analysis (see inclusion/exclusion criteria in Methods) and therefore, only few data points remained for the power analysis. CD8 negative T cell expressing TCRs stimulated in a plate assay produced *α* = 0.12 (ID 27), however in the cellular assay TCRs binding to the antigen with a K_D_ *>* 1 were not activated in CD8 negative T cells (no data points to calculate *α*) (ID 28). TCRs in CD8 positive T cells stimulated in the cellular assay produce *α* = 0.66 (ID 29).

##### ID 30 (98)

The 2C high affinity TCR and variants thereof binding to the QL9 and the altered QL9 peptide F5R were studied. The K_D_ values were measured by SPR at 25*^◦^*C. Functional data was generated with T cell hybridomas stimulated by peptide-pulsed APCs with T cell activation assessed by IL-2 production. K_D_ and *EC*_50_ values were taken from Table 1. K_D_ values below 1*µ*M were excluded from our power analysis. This data produces *α* = 2.7 (ID 30).

##### ID 31-32 (99)

The authors report binding and functional responses of high-affinity 2C TCR variants interacting with SIY peptide on MHC Kb and QL9 peptide on Ld. In total, 8 different TCR/pMHC ligand pairs were included. The K_D_ values were measured by SPR at 25*^◦^*C and provided in Table 1. K_D_ values lower than 1 *µ*M were excluded from the analysis. Functional assays were done with T cell hybridoma with and without CD8 expression with T cell activation assessed by IL-2 production in response to peptide-pulsed APCs. We extracted potency values as *P*_50_ from dose response curves in Figure 3. TCR variants m6 and m13 when binding to SIY-Kb showed no activation (*P*_50_ *>* 100*µ*M). The calculated discrimination power is *α* = 4.7 for CD8 positive (ID 31) and *α* = 6.5 for CD8 negative T cells (ID 32).

#### Not included (120)

This study provided binding and affinity data for the 2C TCR with and without CD8. However, when applying our inclusion/exclusion criteria only a single data point was available and therefore, we were unable to calculate *α*. The reason is that only few interactions were measured by SPR and the majority of these produced K_D_ values below 1 *µ*M.

#### 42F3

##### ID 33 (100)

The 42F3 TCR recognises the class I MHC molecule H2-Ld presenting the peptide p2Ca. The K_D_ values were measured by SPR at 25*^◦^*C and potency data (*EC*_50_ of of IL2 production after cellular stimulation) were taken from Table 1 and Supplementary Figure 3C. The resulting *α* is 0.15 (ID 33).

#### Gp70 (AH1)-specific TCR

##### ID 34 (121)

The TCR used in this study recognises the AH1 peptide which is derived from the endogenous retroviral protein gp70(423–431), a MHC class I restricted tumor-associated antigen. The authors used a set of AH1 variants with optimised affinities. The K_D_ values were measured by SPR at 25*^◦^*C and provided in Figure 1B. Functional data was generated with a T cell line incubated with peptide-pulsed APCs. *EC*_50_ values of a proliferation assay are provided in Figure 2B. The calculated discrimination power was *α* = 5.2 (ID 34).

### Other human TCRs

#### 1G4

##### ID 35 (59)

The 1G4 TCR used in this study binds the NY-ESO-1 (157–165) peptide loaded on MHC class I HLA-A2. The authors generated a panel of TCRs derived from the human 1G4 TCR that bind with higher affinity than the wild type TCR. The K_D_ values were measured by SPR at 25*^◦^*C and provided in a table in Figure 1A. Potency was measured with a cytotoxicity assay and we extracted the mean *EC*_50_ values from Figure 5E. A decrease in potency was observed for TCRs with an affinity of K_D_ < 1 *µ*M, which were excluded as per our exclusion criteria (see Methods). This data produced *α* = 0.67 (ID 35).

##### ID 36 (122)

Here, TCR–peptide–MHC binding parameters and T cell function was investigated with a panel of 1G4 TCR variants binding to the the NY-ESO-1 peptide. The K_D_ values were measured by SPR and provided in Table 1. The functional response of T cells was determined in a cytotoxic T cell assay. We extracted the mean *EC*_50_ values from Figure 4B. Data points with K_D_ *<* 1*µ*M are excluded from the power analysis. The resulting *α* is 0.69 (ID 36).

##### ID 37 - 38 (37)

Here, the interaction between 1G4 TCR binding a set of variant NY-ESO-1 (157–165) peptides on MHC class I was studied. The K_D_ values were measured by SPR at 37*^◦^*C. The potency was determined by IFN*γ* production of T cell after stimulation by plate-immobilised pMHC or cytotoxicity by peptide-pulsed T2 APCs. The 1G4 TCR clone was used for both experiments. All affinity and *EC*_50_ values were given in Table S1. Discrimination power *α* for the 1G4 system is 0.6 (IFN*γ*, ID 37) and 1.6 (Cytotoxicity assay, ID 38).

#### 1G4 and G10

##### ID 39 - 40 (38)

Experimental data was generated with the 1G4 and G10 TCR clones binding to a panel of peptide variants. The 1G4 TCR recognises the NY-ESO-1 antigen and the G10 TCR recognises the HIV gag p17 antigen in the context of MHC class I HLA-A2. The K_D_ values were measured by SPR at 37*^◦^*C. Potency was determined by measuring IFN*γ* production in response to plate-immobilised recomvbinant pMHC. All K_D_ and *EC*_50_ values were given in Table S1 and S2. For the 1G4 system we found *α* = 0.55 (ID 39) and for the G10 system we found *α* = 0.95 (ID 40).

#### 1E6

##### ID 41-42 (53)

The MHC-I restricted 1E6 TCR reactive to preproinsulin (INS) and variants were studied. The K_D_ values were measured by SPR at 25 and 37*^◦^*C and provided in Figure 2. All K_D_ values lower than 1*µ*M were excluded from the power analysis (see Methods). Functional assays were done with primary T cells responding to peptide-pulsed APCs and target cell lysis was measured for T cell activation. The *EC*_50_ was determined from the data in Figure 2K. We calculated *α* = 1.1 for K_D_ values measured at 25*^◦^*C (ID 41) and *α* = 1.2 for K_D_ values measured at 37*^◦^*C (ID 42).

#### A6

##### ID 43 −44 (103)

The A6 and engineered variants recognising the Tax or HuD peptides were used. The K_D_ values were measured by SPR at 25*^◦^*C and provided in Figure 1A. T cell activation in response to peptide-pulsed APCs was assessed by CD107a expression and IFN*γ* production. Potency data was extracted as *P*_20_ for CD107a assay from dose response curve in Figure 4C and as *P*_10_ for IFN*γ* assay from dose response curve in Figure 5A. Data point with K_D_ *<* 1*µ*M were not included in the power analysis. The resulting *α* is 2.0 (ID 43) and 2.2 (ID 44) for CD107 and IFN*γ* readout, respectively.

#### Gp100-specific TCR (Melanoma)

##### ID 45-46 (105)

Seven TCRs specific to human melanoma gp209–2M epitope (modified from gp100 (209–217)) were isolated from patients vaccinated with gp209– 2M. The K_D_ values of these TCRs measured by SPR at 25*^◦^*C was provided in Table 1. Functional activity was determined by IFN*γ* production and ERK phosphorylation of transduced CD8^+^ splenocytes mixed with peptide-pulsed APCs. Potency values were extracted from Figure S3A and C as *P*_10_. The L2G2 TCR, which appeared as an extreme outlier showing the highest potency despite having the lowest affinity, was excluded from the analysis. This data point is shown in the plots as an open circle and including it would have further reduced the estimates *α*. The calculated powers were *α* = 1.3 for IFN*γ* production (ID 45) and *α* = 1.2 for ERK phosphorylation assay (ID 46).

##### ID 47 −48 (106)

T cell responses of a TCR specific to melanoma epitope gp100(280–288) were studied using a set of altered peptides. The K_D_ values were measured by SPR at 25*^◦^*C and provided in Table 2. Functional assays used gp100 TCR-transduced CD8+ T-cells stimulated by peptide-pulsed APCs with T cell activation assessed by cytotoxic lysis and MIP-1*β* production. We extracted the potency data as *P*_10_ from dose response curves in Figure 6. The resulting *α* values were 2.3 (lysis assay, ID 47) and 3.6 (MIP-1*β* production, ID 48).

#### 14.3.d

##### ID 49 - 50 (107)

T cell responses were measured using variants of the Staphylococcus enterotoxin C3 (SEC3) super antigen. In addition, binding of a panel of mutated variants of the antibody F23.1 were also used. The K_D_ values of SEC3 were measured by SPR and provided in Table 1. The K_D_ values of the antibodies were provided in Table 1 of different publication (108). T cell hybridomas, containing a NFAT-GFP expression cassette, were stimulated with SEC3 or antibody molecules immobilised onto plate surfaces to observe functional responses. We extracted all potency values as EC20 from Figure 4. According to our exclusion criteria (see Methods), we did not include any data point where K_D_ *<* 1 *µ*M. The remaining data points generated with the SEC3 variants produced *α* = 0.81 (ID 49) and with the F23.1 antibody variants produced *α* = 0.66 (ID50).

#### TCR55

##### ID 51 (51)

This study used TCRs specific for HLA-B35-HIV(Pol448–456) binding to a set of variant peptides. The K_D_ values were measured by SPR at 25*^◦^*C. T cell activation after stimulation with peptide pulsed on APCs was measured by CD69 expression. All K_D_ and *EC*_50_ values were given in Figure S5C. We calculated *α* = 0.19 (ID 51).

#### ILA1

##### ID 52 - 57 (35)

The MHC-class I restricted ILA1 TCR is specific for the human telomerase reverse transcriptase (hTERT) epitope ILAKFLHWL (hTERT540-548). The K_D_ values were measured by SPR at 25*^◦^*C and provided in Table 1. Three different assays were used to measure T cell activation: Degranulation assay, CD107a expression, and IFN*γ* production. Each assay was performed using APCs expressing either WT MHC or CD8-null MHC which cannot bind CD8. Potency values for degranulation were given in Table 1, CD107a and IFN*γ* potency data was extracted from dose response curves in Figure 7. For potency data measured with wild-type (WT) and CD8 null MHC respectively, we calculated an *α* of 1.5 (WT, ID 52) and 2.5 (CD8 neg., ID 52) for degranulation, 2.2 (WT, ID 54) and 3.6 (CD8 neg., ID 55) for CD107a, and 2.2 (WT, ID 56) and 3.2 (CD8 neg., ID 57) for IFN*γ* production.

##### ID 58 - 61 (109)

The ILA1 TCR was studied interacting with peptide variants. The K_D_ values were measurred by SPR at 25*^◦^*C and provided in Table 1. T cell activation was measured by peptide-pulsed APCs and dertermined by MIP-1*β*, IFN*γ*, TNF*α*, and IL-2 production using intracellular cytokine staining. The potency values were read of as *P*_50_ from the dose response curves in Figure 2. Authors suggested that the TCR shows a plateau at K_D_ values < 5 *µ*M. Therefore we decided to exclude K_D_ values < 5 *µ*M from the power analysis to avoid underestimating the discrimination power *α*. The data produces *α* = 1.4 (ID 58), 0.77 (ID 59), 0.97 (ID 60) and 1.1 (ID 61) for MIP-1*β*, IFN*γ*, TNF*α* and IL-2 production respectively.

#### NY-ESO-1(60–72)-specific TCR

##### ID 62 (110)

Four TCRs binding to the tumorigenic antigen NY-ESO-1(60–72) were obtained from patients with melanomas expressing NY-ESO-1. The K_D_ values were measured by SPR 25*^◦^*C and given in Figure 2C. Functional response of TCRs to exogenous peptide stimulation was assessed by measuring IFN*γ* production of T cells incubated with NY-ESO-1-expressing melanoma cells. We extracted *EC*_50_ values from dose response curves in Figure 1F. We calculated *α* = *−*0.59 (ID 62).

#### Gliadin-specific TCRs (Celiac disease)

##### ID 63 (111)

Seven DQ8-glia-a1-restricted T cell receptors isolated from celiac disease patients were characterised for their binding affinity to a-I-gliadin and their functional response. The K_D_ values were measured by SPR at 25*^◦^*C and provided in Figure 2 and S5. T cell activation was assessed by proliferation in response to peptide-pulsed APCs. We extracted *P*_20_ values from the dose-response curves in Figure 1 (Black curve Q-Q). We calculated an *α* = 0.83 (ID 63).

#### LC13

##### ID 64 - 67 (36)

The LC13 and SB27 TCRs were studied using an alanine scan. The K_D_ values were measured by SPR and provided in Table S2. T cell activation was measured using Jurkat T cells expressing the TCR with CD69 and cytoxicity assessed in response to peptide-pulsed APCs. Figure 1C and 1D showed the dose response curves for CD69 upregulation for either CD8 positive or CD8 negative cells. We extracted the *P*_30_ as potency measure. *EC*_50_ of cytotoxicity assay was given in Figure 2 for LC13 and Figure S2 for SB27. Potency values from CD69 produced *α* = 1.9 (ID 64) for CD8 positive cells and *α* =7.8 (ID 65) for CD8 negative cells. Lysis assays produced *α* = 4.1 (ID 66) for the LC13 and *α* = 0.11 (ID 67) for SB27 TCR.

#### HIV-Gag293-specific TCRs

##### ID 68 (112)

TCRs specific to HIV Gag293 protein were isolated from patients infected with HIV. The K_D_ values were measured by SPR and provided in Table 3. T cell activation was assessed using TCR-transduced J76 cells measuring CD69 expression in response to peptide-pulsed APCs. We extracted the mean *EC*_50_ values from Figure 6D. We calculated *α* = 1.0 (ID 68).

#### MEL5

##### ID 69 (113)

The MEL5 and MEL187.c5 TCRs were studied that bind the MART-1 antigen and variants thereof. The K_D_ values were measured by SPR at 25*^◦^*C and provided in Table 1. T cell activation was measured by MIP-1*β* production in response to peptide-pulsed APCs. We extracted potency values as *P*_50_ from dose response curves in Figure 2 and S1. Because responses to peptides were measured in separate experiments, potency data is normalised to wild type peptide. This produced *α* = 2.3 (ID 69).

##### ID 70 (114)

The MEL5, MEL187.c5, DMF4, and DMF5 were studied that recognise the MART-1 antigen. Two overlapping peptides were used: nonapeptide MART-1(27–35) and decapeptide MART-1(26–35). The K_D_ values were measured by SPR at 25*^◦^*C and provided in Table 1. T cell activation was assessed using primary human T cells responding to peptide-pulsed APCs with MIP-1*β* used as a marker of T cell activation. We determined *P*_30_ directly form does response curves in Figure 1A. Data produced *α* = 4.5 (ID 70)

### Other (non-TCR) surface receptors

#### Cytokine receptors

##### ID 1-2 (76)

Engineered IL-2 variants with increased binding affinity for the interleukin-2 (IL-2) receptor subunit *β* (IL-2Rb) were studied. The K_D_ values for IL-2 variants to IL-2R*β* are given in Supplementary Figure 3 and determined by SPR at 25*^◦^*C. As only the affinities to a single subunit were varied between ligands, potency was plotted over these K_D_ values. Functional experiments were performed with either CD25 negative human Natural Killer cells or CD25 negative murine T cells. We extracted the *EC*_50_ values as a measure of potency from dose-response curves in Figure 3a and 3e. We calculated *α* = 0.55 (ID 1) for experiments done with Natural Killer cells and *α* = 0.74 (ID 2) for T cells.

##### ID 3-6 (77)

The relationship between the interaction of interleukin 13 (IL-13) with its cytokine receptor and the resulting downstream cellular responses were investigated. A panel of IL-13 variants with a range of binding affinities for the receptor subunit IL-13Ra1 was generated. Binding affinities of these ligands were given in a table in Figure 2C. Here, only the affinity for the *α* subunit of the receptor dimer was varied and therefore, we plotted potency over these K_D_ values. Functional responses of binding were determined by measuring STAT6 phosphorylation, CD86 and CD209 production, and proliferation after receptor stimulation. We extracted *EC*_50_ values for pSTAT6 from Figure 5B. To avoid extrapolating potencies, ligands with *EC*_50_ larger than highest concentration used in the dose-response (in Figure 5A) were excluded. The mean proliferation *EC*_50_ values were taken from Figure 5G. CD86 *EC*_50_ values were extracted from dose response curve in Figure 5H, CD209 *EC*_50_ values from the dose response curve in Figure S7C. *EC*_50_ values for CD86 and CD209 extracted from the dose response curves did not exactly match *EC*_50_ values given in Figure S7 D and E, but both values resulted in similar *α* values. The *α* values calculated for the IL-13 receptor are 0.47 (ID 3), 0.39 (ID 4), 0.44 (ID 5), and 0.42 (ID 6) for potency values from pSTAT6, proliferation, CD86, and CD205 assays, respectively.

##### ID 7-8 (78)

Study uses a set of mutated cytokines derived from IFN*α*2 and IFN*ω*, binding cytokine receptors IFNAR1 and IFNAR2. All binding affinities of mutants normalised to WT are provided in Supplemental Table 2. Because mutations change the affinities to both IFNAR1 and IFNAR2 we calculated an effective binding affinity by multiplying K_D_ of IFNAR1 with K_D_ of IFNAR2 (R1xR2). Functional response of cells to cytokine mutants was determined by their antiviral activity in a Hepatitis C Virus Replication Assay, their antiproliferation activity on WISH cells. Mean *EC*_50_ values normalised to WT obtained from Figure 7A. We calculated *α* = 0.71 (ID 7) for antiviral potency and *α* = 1.3 (ID 8) for antiproliferation potency.

##### ID 9-11 (79)

Study of IFN1 receptor activation with engineered higher-affinity type I IFNs. Affinity constants for peptides to each receptor subunit were measured by SPR. To get the effective K_D_ we multiplied K_D_ of IFN-*α*R1 bindning with K_D_ of IFN-*α*R2 binding (R1xR2) Ligand activity was measured by STAT phosphorylation, antiviral activity and antiproliferation activity. All affinity and *EC*_50_ values were provided in Table S2. The data produced *α* = 0.024 for STAT1 phosphorylation (ID 9), *α* = 0.034 for antiviral activity (ID 10), and *α* = 0.50 (ID 11) for the anti-proliferation assay.

##### ID 12-13 (80)

In this study, the authors engineered IL-6 variants with different affinities to the IL-6 receptor subunit gp130. Cytokine gp130 binding kinetics were measured with a switchSENSE chip, binding parameters were given in Supplementary Figure 1D. The influence of IL-6 variants on functional activity of the receptor was determined by the amount of STAT1 and STAT3 phosphorylation at different ligand concentrations. We read off the potency of each ligand as *P*_25_ directly from dose response curves in Figure 2A and B. We calculated *α* = 0.54 for pSTAT1 (ID 12) and *α* = 0.52 for pSTAT 3 (ID 13).

#### Receptor Tyrosine Kinase (RTK)

##### ID 14 (82)

In this study the effect of three mutated epidermal growth factor on epidermal growth factor receptor (EGFR) was studied. Affinity values of growth factor to receptor were measured with radioactive labelled ligands binding to receptors on cells. Data are given in Table 1. Functional response of cells to ligands was determined by measuring the specific growth rate after stimulation. We extracted the *EC*_50_ values from dose response curves in Figure 4. This produced *α* = 0.55 (ID 14).

##### ID 15 - 16 (81)

Paper contains data on the c-Kit receptor tyrosine kinase which is activated by the Stem cell factor (SCF). Affinity and functional response of the receptor to SCF variants was studied. Binding parameters were measured by SPR and provided in Figure 1F. Cell activation after stimulation with ligands was determined by the amount of ERK and AKT phosphorylation (pERK and pAKT). We extracted the potency data for each variant as *EC*_50_ from dose response curves in Figure 2D and 2E. We calculated *α* = 0.83 (ID 15) and *α* = 0.88 (ID 16) for pERK and pAKT measurements respectively.

#### GPCRs

##### ID 17 −18 (84)

The binding parameters of the GPCR adenosine A2A receptor to various agoinist and their functional effects were studied. Association and dissociation rates, and hence K_D_ values, were determined with a kinetic radioligand binding assay. Functional activity of HEK293 expressing the A2A receptor was measured by detecting cAMP production and changes in cell morphology. The binding data was provided in Table 3 and *EC*_50_ values from functional experiments were given in Table 4. The discrimination power calculated with cell morphology data is *α* = 0.29(ID 17) and with the cAMP assay produced *α* = 0.71 (ID 18).

##### ID 19-20 (83)

The M3 muscarinic receptor was studied using a set of agaonist. The binding kinetics were determined with competition binding assay and were provided in Table 1. Agonist potency was measured by guanosine 5’-O- (3-[35S]thio) triphosphate (GTP*γ*S) binding to G*α*D subunits, and by intracellular calcium levels after receptor stimulation. Potency data measured as *EC*_50_ values were provided in Table 2. The resulting power was *α* = 0.77 (ID 19) and *α* = 0.55 (ID 20) for calcium response and GTP*γ*S binding assay, respectively.

##### ID 21 (85)

The CXCR4 receptor is activated by the chemokine CXCL12. In this paper, the interaction of Baclofen and other GABA ligands were tested on their abilities to activate CXCR4. The affinity of ligands to the receptor was measured by back-scattering interferometry and K_D_ values given in Figure 7. Functional response of oocytes expressing CXCR4 to ligands was determined by measuring the inward currents at different ligand concentrations. *EC*_50_ values were provided in Table 1. We calculated *α*D = 0.57 for this system (ID 21).

##### ID 22 −24 (86)

Characterisation of binding properties and potencies of CXC chemokine receptor 3 antagonists. Binding properties of antagonist were determined using kinetic radioligand binding assay. Affinity values were in Table 1 measured for different cell lines. Functional responses after ligand binding were measured guanosine 5’-O-(3-[35S]thio)triphosphate (GTPyS) binding, calcium release, and cellular chemotaxis. All *EC*_50_ values of assays were given in the text. We calculated *α*D = 0.72 (ID 22), 1.1 (ID 23), and 0.56 (ID 24) for calcium release, GTPyS binding, chemotaxis assays respectively.

#### CARs

##### ID 25 (115)

This study contain affinity and potency data for a CAR binding the ErbB2 surface antigen. The authors generated a series of anti-ErB2 single-chain variable fragments fused to the CD3*ζ* cytoplasmic domain. The K_D_ values are reported in Table 1. Functional experiments were done in a plate assay, with ErbB2 immobilised to a surface. Potency of receptors was measured by IFN*γ* production of T cells after stimulation. We extracted *P*_20_ values from dose response curve in Figure 4A. The resulting *α* is 0.52 (ID 25).

##### ID 26-27 (88)

This study characterised a panel of CARs that bind to the ErbB2 surface protein. CARs were constructed by linking the various anti-ErB2 single-chain variable fragments to the CD8*α*D hinge and transmembrane domain followed by the 4-1BB and CD3*ζ* intracellular signalling domains. The K_D_ values were measured by SPR and provided in Table S1. For functional experiments CAR T cells were incubated with ErbB2 expressing cells. We obtained potency data by using CD107a expression and proliferation assay data in Figure 2A and C to the respective plot dose response curves. *P*_50_ values were extracted from these plots. The resulting *α* values are 1.1 for CD107 (ID 26) and 0.64 for proliferation assay (ID 27).

##### ID 28 (89)

Taylor et al (89) developed a synthetic CAR signaling system in which the extracellular domains of the CAR and its ligand antigen were exchanged with short hybridizing strands of DNA. The DNA-CAR*ζ* consist of a ssDNA covalently attached to a SNAP tag protein which was fused to a transmembrane domain and the CD3*ζ* chain. Stands of different length and sequence were designed to vary the affinity of the CAR to the ligand. Binding was measured as the lifetime (*τ_corr_*) of single ligand-CAR interactions using microscopy and corrected for photobleaching and provided in Figure 2D. The dissociation rate *k_o_ff* was calculated from the lifetimes with *k*_off_ = *ln*(2)/*τ*. To measure T cell responses, ligands, consisting of the complimentary strand of ssDNA, were anchored in planar supported lipid bilayer where they can freely diffuse. The DNA-CAR*ζ* was expressed in TCR negative Jurkat cells. Cell activation after incubation with ligands was measured by phosphorylation of ERK. Potency data was extracted as *P*_20_ from dose response curves in Figure 2C. This CAR system produced *α* = 1.2 (ID 28).

#### BCRs

##### ID 29-30 (90)

The study used the HyHEL10 and D1.3 BCRs, which have a high affinity to the hen egg lysozyme (HEL) and variants thereof. The K_D_ values were measured by SPR at 25*^◦^*C and dissociation rates were provided in Table 1. For functional experiments, the ability of B cells to mediate HEL presentation to T cell hybridomas after stimulation with mutant lysozymes was determined by measuring IL-2 production of T cells specific to HEL. We extracted the potency data from dose response curves in Figure 3 and 4 as *EC*_50_. The authors described an affinity floor for the B cell receptor when the dissociation rate was below 10*^−^*^4^ s*^−^*^1^ so that potency did not longer decrease for these interactions. To avoid underestimating *α*, we did not include these higher affinity ligands in the power analysis. The resulting *α* values were *α* = 1.4 for the D1.3 BCR (ID 29) and *α* = 1.3 for the HyHEL10 BCR (ID 30).

## Footnotes

1 Squaring both sides will not introduce a false solution so long as 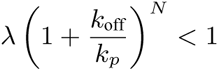.

## References

1. Francois P, Voisinne G, Siggia ED, Altan-Bonnet G, Vergassola M (2013) Phenotypic model for early T-cell activation displaying sensitivity, specificity, and antagonism. Proceedings of the National Academy of Sciences.

2. Liu B, Chen W, Evavold BD, Zhu C (2014) Accumulation of Dynamic Catch Bonds between TCR and Agonist Peptide-MHC Triggers T Cell Signaling. Cell 157:357–68.

3. Dushek O, van der Merwe PA (2014) An induced rebinding model of antigen discrimination. Trends in Immunology 35:153–158.

4. Chakraborty AK, Weiss A (2014) Insights into the initiation of TCR signaling. Nature Immunology 15:798–807.

5. Hong J, et al. (2018) A TCR mechanotransduction signaling loop induces negative selection in the thymus. Nature Immunology 19:1379–1390.

6. Fernandes RA, et al. (2019) A cell topography-based mechanism for ligand discrimination by the T cell receptor. Proceedings of the National Academy of Sciences of the United States of America 116:14002–14010.

7. Wu P, et al. (2019) Mechano-regulation of Peptide-MHC Class I Conformations Determines TCR Antigen Recognition. Molecular Cell 73:1015–1027.e7.

8. Ganti RS, et al. (2020) How the T cell signaling network processes information to discriminate between self and agonist ligands. Proceedings of the National Academy of Sciences p 202008303.

9. Hogquist KA, Jameson SC, Bevan MJ (1995) Strong agonist ligands for the t cell receptor do not mediate positive selection of functional cd8+ t cells. Immunity 3:79–86.

10. Alam SM, et al. (1996) T-cell-receptor affinity and thymocyte positive selection. Nature 381:616–20.

11. Alam S, et al. (1999) Qualitative and Quantitative Differences in T Cell Receptor Binding of Agonist and Antagonist Ligands. Immunity 10:227–237.

12. Kersh GJ, Allen PM (1996) Structural basis for T cell recognition of altered peptide ligands: a single T cell receptor can productively recognize a large continuum of related ligands. J Exp Med 184:1259–1268.

13. Kersh GJ, Kersh EN, Fremont DH, Allen PM (1998) High- and low-potency ligands with similar affinities for the TCR: the importance of kinetics in TCR signaling. Immunity 9:817–26.

14. Lyons DS, et al. (1996) A TCR binds to antagonist ligands with lower affinities and faster dissociation rates than to agonists. Immunity 5:53–61.

15. Altan-Bonnet G, Germain RN (2005) Modeling T cell antigen discrimination based on feedback control of digital ERK responses. PLoS biology 3:e356.

16. McKeithan TW (1995) Kinetic proofreading in T-cell receptor signal transduction. Proceedings of the National Academy of Sciences of the United States of America 92:5042–6.

17. Huang J, et al. (2013) A Single Peptide-Major Histocompatibility Complex Ligand Triggers Digital Cytokine Secretion in CD4+ T Cells. Immunity pp 1–12.

18. Siller-Farfán JA, Dushek O (2018) Molecular mechanisms of T cell sensitivity to antigen. Immuno-logical Reviews 285:194–205.

19. Yin Y, Li Y, Mariuzza RA (2012) Structural basis for self-recognition by autoimmune T-cell receptors. Immunological Reviews 250:32–48.

20. Bridgeman JS, Sewell AK, Miles JJ, Price DA, Cole DK (2012) Structural and biophysical determinants of *αβ* T-cell antigen recognition. Immunology 135:9–18.

21. Korem Kohanim Y, Tendler A, Mayo A, Friedman N, Alon U (2020) Endocrine Autoimmune Disease as a Fragility of Immune Surveillance against Hypersecreting Mutants. Immunity 52:872–884.e5.

22. Wang J, et al. (2020) HLA-DR15 Molecules Jointly Shape an Autoreactive T Cell Repertoire in Multiple Sclerosis. Cell 2.

23. van der Merwe PA, Davis SJ (2003) Molecular interactions mediating T cell antigen recognition. Annual review of immunology 21:659–84.

24. Stepanek O, et al. (2014) Coreceptor scanning by the T cell receptor provides a mechanism for T cell tolerance. Cell 159:333–345.

25. Liu X, et al. (2015) Affinity-tuned ErbB2 or EGFR chimeric antigen receptor T cells exhibit an increased therapeutic index against tumors in mice. Cancer Research 75:3596–3607.

26. Chen JL, et al. (2005) Structural and kinetic basis for heightened immunogenicity of T cell vaccines. The Journal of experimental medicine 201:1243–55.

27. D.N. G, et al. (1996) Structure of the complex between human T-cell receptor, viral peptide and HLA-A2. Nature 384:134–141.

28. Brodskys FM, Parham P (1982) Monomorphic anti-HLA-A, B, C monoclonal antibodies detecting molecular subunits and combinatorial determinants. Journal of immunology 128:129–135.

29. Daniels MA, et al. (2006) Thymic selection threshold defined by compartmentalization of Ras/MAPK signalling. Nature 444:724–729.

30. Hong J, et al. (2015) Force-Regulated In Situ TCR–Peptide-Bound MHC Class II Kinetics Determine Functions of CD4 + T Cells. The Journal of Immunology 195:3557–3564.

31. Wu L, Tuot D, Lyons D, Garcia K, Davis M (2002) Competing interests statement Two-step binding mechanism for T-cell receptor recognition of peptide–MHC. Nature 418:552–556.

32. Birnbaum ME, et al. (2014) Deconstructing the peptide-MHC specificity of t cell recognition. Cell 157:1073–1087.

33. Newell EW, et al. (2011) Structural Basis of Specificity and Cross-Reactivity in T Cell Receptors Specific for Cytochrome c –I-E k. The Journal of Immunology 186:5823–5832.

34. Lo WL, et al. (2019) Slow phosphorylation of a tyrosine residue in LAT optimizes T cell ligand discrimination. Nature Immunology 20:1481–1493.

35. Laugel B, et al. (2007) Different T cell receptor affinity thresholds and CD8 coreceptor dependence govern cytotoxic T lymphocyte activation and tetramer binding properties. Journal of Biological Chemistry 282:23799–23810.

36. Burrows SR, et al. (2010) Hard wiring of t cell receptor specificity for the major histocompatibility complex is underpinned by tcr adaptability. Proceedings of the National Academy of Sciences 107:10608–10613.

37. Aleksic M, et al. (2010) Dependence of T cell antigen recognition on T cell receptor-peptide MHC confinement time. Immunity 32:163–74.

38. Dushek O, et al. (2011) Antigen potency and maximal efficacy reveal a mechanism of efficient T cell activation. Science Signaling 4:ra39.

39. Lever M, et al. (2016) A minimal signalling architecture explains the T cell response to a 1,000,000-fold variation in antigen affinity and dose. Proc Natl Acad Sci USA pp E6630–E6638.

40. Abu-Shah E, et al. (2020) Human CD8 + T Cells Exhibit a Shared Antigen Threshold for Different Effector Responses. The Journal of Immunology 205:1503–1512.

41. Bachmann MF, et al. (1997) Distinct Roles for LFA-1 and CD28 during Activation of Naive T Cells : Adhesion versus Costimulation. Immunity 7:549–557.

42. Bachmann MF, Barner M, Kopf M (1999) CD2 sets quantitative thresholds in T cell activation. Journal of Experimental Medicine 190:1383–1392.

43. Lever M, Maini PK, van der Merwe PA, Dushek O (2014) Phenotypic models of T cell activation. Nature Reviews Immunology 14:619–629.

44. Yousefi OS, et al. (2019) Optogenetic control shows that kinetic proofreading regulates the activity of the T cell receptor. eLife 8:1–33.

45. Huse M, et al. (2007) Spatial and temporal dynamics of T cell receptor signaling with a photoactivatable agonist. Immunity 27:76–88.

46. Tischer DK, Weiner OD (2019) Light-based tuning of ligand half-life supports kinetic proofreading model of T cell signaling. eLife 8:1–25.

47. Wang H, et al. (2010) ZAP-70: an essential kinase in T-cell signaling. Cold Spring Harbor Perspectives in Biology 2.

48. Goyette J, et al. (2020) Regulated unbinding of ZAP70 at the T cell receptor by kinetic avidity. bioRxiv.

49. Gudipati V, et al. (2020) Inefficient CAR-proximal signaling blunts antigen sensitivity. Nature Immunology 21:848–856.

50. Adams JJ, et al. (2011) T cell receptor signaling is limited by docking geometry to peptide-major histocompatibility complex. Immunity 35:681–693.

51. Sibener LV, et al. (2018) Isolation of a Structural Mechanism for Uncoupling T Cell Receptor Signaling from Peptide-MHC Binding. Cell 174:672–687.e27.

52. Huang J, et al. (2010) The kinetics of two-dimensional TCR and pMHC interactions determine T-cell responsiveness. Nature 464:932–6.

53. Cole DK, et al. (2016) Hotspot autoimmune T cell receptor binding underlies pathogen and insulin peptide cross-reactivity. Journal of Clinical Investigation 126:2191–2204.

54. Daniels Ma, Schober SL, Hogquist Ka, Jameson SC (1999) Cutting edge: a test of the dominant negative signal model for TCR antagonism. Journal of immunology (Baltimore, Md. : 1950) 162:3761–4.

55. Yang W, Grey HM (2003) Study of the mechanism of TCR antagonism using dual-TCR-expressing T cells. Journal of immunology 170:4532–8.

56. Stone JD, et al. (2011) Opposite effects of endogenous peptide-MHC class I on T cell activity in the presence and absence of CD8. Journal of immunology (Baltimore, Md. : 1950) 186:5193–200.

57. Kalergis aM, et al. (2001) Efficient T cell activation requires an optimal dwell-time of interaction between the TCR and the pMHC complex. Nature immunology 2:229–34.

58. Corse E, Gottschalk Ra, Krogsgaard M, Allison JP (2010) Attenuated T cell responses to a high-potency ligand in vivo. PLOS BIOLOGY 8:1–12.

59. Irving M, et al. (2012) Interplay between T cell receptor binding kinetics and the level of cognate peptide presented by major histocompatibility complexes governs CD8+ T cell responsiveness. The Journal of biological chemistry 287:23068–78.

60. Wooldridge L, et al. (2010) CD8 Controls T Cell Cross-Reactivity. The Journal of Immunology 185:4625–4632.

61. Cameron BJ, et al. (2013) Identification of a titin-derived HLA-A1-presented peptide as a crossreactive target for engineered MAGE A3-directed T cells. Science Translational Medicine 5.

62. Chan C, George AJ, Stark J (2003) T cell sensitivity and specificity - Kinetic proofreading revisited. Discrete and Continuous Dynamical Systems - Series B 3:343–360.

63. Bushell KM, Sollner C, Schuster-Boeckler B, Bateman A, Wright GJ (2008) Large-scale screening for novel low-affinity extracellular protein interactions. Genome Res 18:622–30.

64. Abu-Shah E, et al. (2019) A tissue-like platform for studying engineered quiescent human t-cells’ interactions with dendritic cells. eLife 8.

65. Achour A, et al. (1999) Murine class i major histocompatibility complex h-2dd: expression, refolding and crystallization. Acta Crystallogr D Biol Crystallogr 55:260–2.

66. Parrott MB, Barry MA (2001) Metabolic biotinylation of secreted and cell surface proteins from mammalian cells. Biochemical and biophysical research communications 281:993–1000.

67. Myszka DG (1999) Improving biosensor analysis. Journal of Molecular Recognition 12:279–284.

68. Abu-Shah E, et al. (2019) A tissue-like platform for studying engineered quiescent human T-Cells’ interactions with dendritic cells. eLife 8.

69. Cohen CJ, et al. (2007) Enhanced antitumor activity of t cells engineered to express t-cell receptors with a second disulfide bond. Cancer research 67:3898–903.

70. Ding YH, et al. (1998) Two human t cell receptors bind in a similar diagonal mode to the hla-a2/tax peptide complex using different tcr amino acids. Immunity 8:403–11.

71. Ding YH, Baker BM, Garboczi DN, Biddison WE, Wiley DC (1999) Four a6-tcr/peptide/hla-a2 structures that generate very different t cell signals are nearly identical. Immunity 11:45–56.

72. Gagnon SJ, et al. (2006) T cell receptor recognition via cooperative conformational plasticity. J Mol Biol 363:228–43.

73. Borbulevych OY, et al. (2009) T cell receptor cross-reactivity directed by antigen-dependent tuning of peptide-mhc molecular flexibility. Immunity 31:885–896.

74. Borbulevych OY, Piepenbrink KH, Baker BM (2011) Conformational melding permits a conserved binding geometry in tcr recognition of foreign and self molecular mimics. J Immunol 186:2950–8.

75. Utz U, Banks D, Jacobson S, Biddison WE (1996) Analysis of the t-cell receptor repertoire of human t-cell leukemia virus type 1 (htlv-1) tax-specific cd8+ cytotoxic t lymphocytes from patients with htlv-1-associated disease: evidence for oligoclonal expansion. J Virol 70:843–51.

76. Levin AM, et al. (2012) Exploiting a natural conformational switch to engineer an interleukin-2 ‘su-perkine’. Nature 484:529–533.

77. Moraga I, et al. (2015) Instructive roles for cytokine-receptor binding parameters in determining signaling and functional potency. Science Signaling 8:1–17.

78. Thomas C, et al. (2011) Structural linkage between ligand discrimination and receptor activation by type i interferons. Cell 146:621–632.

79. Mendoza JL, et al. (2017) The IFN-*λ*-IFN-*λ*R1-IL-10R*β* Complex Reveals Structural Features Underlying Type III IFN Functional Plasticity. Immunity 46:379–392.

80. Martinez-Fabregas J, et al. (2019) Kinetics of cytokine receptor traffi 1 cking determine signaling and functional selectivity. eLife 8:1–32.

81. Ho CCM, et al. (2017) Decoupling the functional pleiotropy of stem cell factor by tuning c-kit signaling. Cell 168:1041–1052.

82. Reddy CC, Niyogi SK, Wells A, Wiley HS, Lauffenburger DA (1996) Engineering epidermal growth factor for enhanced mitogenic potency. Nature biotechnology 14:1696–1699.

83. Sykes DA, Dowling MR, Charlton SJ (2009) Exploring the mechanism of agonist efficacy: a relationship between efficacy and agonist dissociation rate at the muscarinic m3 receptor. Molecular pharmacology 76:543–551.

84. Guo D, Mulder-Krieger T, IJzerman AP, Heitman LH (2012) Functional efficacy of adenosine a2a receptor agonists is positively correlated to their receptor residence time. British journal of pharmacology 166:1846–1859.

85. Guyon A, et al. (2013) Baclofen and other GABAB receptor agents are allosteric modulators of the CXCL12 chemokine receptor CXCR4. Journal of Neuroscience 33:11643–11654.

86. Heise CE, et al. (2005) Pharmacological characterization of CXC chemokine receptor 3 ligands and a small molecule antagonist. Journal of Pharmacology and Experimental Therapeutics 313:1263–1271.

87. Chmielewski M, Hombach A, Heuser C, Adams GP, Abken H (2004) T cell activation by antibody-like immunoreceptors: increase in affinity of the single-chain fragment domain above threshold does not increase t cell activation against antigen-positive target cells but decreases selectivity. The Journal of Immunology 173:7647–7653.

88. Liu X, et al. (2015) Affinity-tuned erbb2 or egfr chimeric antigen receptor t cells exhibit an increased therapeutic index against tumors in mice. Cancer research 75:3596–3607.

89. Taylor MJ, Husain K, Gartner ZJ, Mayor S, Vale RD (2017) A DNA-based T cell receptor reveals a role for receptor clustering in ligand discrimination. Cell 169:108–119.

90. Batista FD, Neuberger MS (1998) Affinity dependence of the b cell response to antigen: a threshold, a ceiling, and the importance of off-rate. Immunity 8:751–759.

91. Rosette C, et al. (2001) The impact of duration versus extent of tcr occupancy on t cell activation: a revision of the kinetic proofreading model. Immunity 15:59–70.

92. Kersh EN, Shaw aS, Allen PM (1998) Fidelity of T cell activation through multistep T cell receptor zeta phosphorylation. Science (New York, N.Y.) 281:572–5.

93. Persaud SP, Donermeyer DL, Weber KS, Kranz DM, Allen PM (2010) High-affinity t cell receptor differentiates cognate peptide-mhc and altered peptide ligands with distinct kinetics and thermodynamics. Molecular immunology 47:1793–1801.

94. Krogsgaard M, Prado N, Adams E, He Xl (2003) Evidence that structural rearrangements and/or flexibility during TCR binding can contribute to T cell activation. Molecular cell 12:1367–1378.

95. Tian S, Maile R, Collins E, Frelinger J (2007) CD8+ T cell activation is governed by TCR-peptide/MHC affinity, not dissociation rate. The Journal of Immunology 179:2952.

96. Govern CC, Paczosa MK, Chakraborty AK, Huseby ES (2010) Fast on-rates allow short dwell time ligands to activate T cells. Proceedings of the National Academy of Sciences 107:8724–9.

97. Chervin AS, et al. (2009) The impact of TCR-binding properties and antigen presentation format on T cell responsiveness. Journal of immunology 183:1166–78.

98. Bowerman NA, et al. (2009) Engineering the binding properties of the t cell receptor: peptide: Mhc ternary complex that governs t cell activity. Molecular immunology 46:3000–3008.

99. Jones LL, Colf LA, Stone JD, Garcia KC, Kranz DM (2008) Distinct cdr3 conformations in tcrs determine the level of cross-reactivity for diverse antigens, but not the docking orientation. The Journal of Immunology 181:6255–6264.

100. Adams JJ, et al. (2016) Structural interplay between germline interactions and adaptive recognition determines the bandwidth of tcr-peptide-mhc cross-reactivity. Nature immunology 17:87–94.

101. McMahan RH, et al. (2006) Relating TCR-peptide-MHC affinity to immunogenicity for the design of tumor vaccines. Journal of Clinical Investigation 116:2543–2551.

102. Schmid Da, et al. (2010) Evidence for a TCR affinity threshold delimiting maximal CD8 T cell function. Journal of immunology (Baltimore, Md. : 1950) 184:4936–4946.

103. Thomas S, et al. (2011) Human t cells expressing affinity-matured tcr display accelerated responses but fail to recognize low density of mhc-peptide antigen. *Blood*, The Journal of the American Society of Hematology 118:319–329.

104. Thomas S, et al. (2011) Human T cells expressing affinity-matured TCR display accelerated responses but fail to recognize low density of MHC-peptide antigen. Blood 118:319–29.

105. Zhong S, et al. (2013) T-cell receptor affinity and avidity defines antitumor response and autoimmunity in T-cell immunotherapy. Proceedings of the National Academy of Sciences.

106. Bianchi V, et al. (2016) A Molecular Switch Abrogates Glycoprotein 100 (gp100) T-cell Receptor (TCR) Targeting of a Human Melanoma Antigen. Journal of Biological Chemistry 291:8951–8959.

107. Andersen PS, Geisler C, Buus S, Mariuzza Ra, Karjalainen K (2001) Role of the T cell receptor ligand affinity in T cell activation by bacterial superantigens. The Journal of biological chemistry 276:33452–7.

108. Andersen PS, Menné C, Mariuzza Ra, Geisler C, Karjalainen K (2001) A response calculus for immobilized T cell receptor ligands. The Journal of biological chemistry 276:49125–32.

109. Tan MP, et al. (2015) T cell receptor binding affinity governs the functional profile of cancer-specific CD8+ T cells. Clinical & Experimental Immunology 180:255–270.

110. Chan KF, et al. (2018) Divergent T-cell receptor recognition modes of a HLA-I restricted extended tumour-associated peptide. Nature Communications 9.

111. Broughton SE, et al. (2012) Biased t cell receptor usage directed against human leukocyte antigen dq8-restricted gliadin peptides is associated with celiac disease. Immunity 37:611–621.

112. Benati D, et al. (2016) Public t cell receptors confer high-avidity cd4 responses to hiv controllers. The Journal of clinical investigation 126:2093–2108.

113. Ekeruche-Makinde J, et al. (2012) T-cell receptor-optimized peptide skewing of the t-cell repertoire can enhance antigen targeting. Journal of Biological Chemistry 287:37269–37281.

114. Madura F, et al. (2019) Tcr-induced alteration of primary mhc peptide anchor residue. European journal of immunology 49:1052–1066.

115. Chmielewski M, Hombach A, Heuser C, Adams GP, Abken H (2004) T Cell Activation by Antibody-Like Immunoreceptors: Increase in Affinity of the Single-Chain Fragment Domain above Threshold Does Not Increase T Cell Activation against Antigen-Positive Target Cells but Decreases Selectivity. The Journal of Immunology 173:7647–7653.

116. Zehn D, Lee SY, Bevan MJ (2009) Complete but curtailed t-cell response to very low-affinity antigen. Nature 458:211.

117. Birnbaum ME, Dong S, Garcia KC (2012) Diversity-oriented approaches for interrogating T-cell receptor repertoire, ligand recognition, and function. Immunological reviews 250:82–101.

118. Govern CC, Paczosa MK, Chakraborty AK, Huseby ES (2010) Fast on-rates allow short dwell time ligands to activate t cells. Proceedings of the National Academy of Sciences 107:8724–8729.

119. Chervin AS, et al. (2009) The impact of tcr-binding properties and antigen presentation format on t cell responsiveness. The Journal of Immunology 183:1166–1178.

120. Holler PD, Kranz DM (2003) Quantitative analysis of the contribution of TCR/pepMHC affinity and CD8 to T cell activation. Immunity 18:255–64.

121. McMahan RH, et al. (2006) Relating tcr-peptide-mhc affinity to immunogenicity for the design of tumor vaccines. The Journal of clinical investigation 116:2543–2551.

122. Schmid DA, et al. (2010) Evidence for a tcr affinity threshold delimiting maximal cd8 t cell function. The Journal of Immunology 184:4936–4946.

